# From Structure to Immunogenicity: Decoding Correlated Dynamics at the Peptide MHC interface to Understand TCR Recognition

**DOI:** 10.1101/2025.05.18.654692

**Authors:** Tom Resink, Benedetta Maria Sala, Renhua Sun, Xiao Han, Evren Alici, Flavio Salazar-Onfray, Tatyana Sandalova, Cheng Zhang, Hans-Gustaf Ljunggren, Adnane Achour

## Abstract

The interaction between a class I peptide-major histocompatibility complex (pMHC) and a T cell receptor (TCR) plays a central role in the elicitation of CD8^+^ T cell immune responses. As a result, considerable effort has been invested in understanding the structural, dynamic, and biophysical parameters that govern this recognition event, including designing altered peptide ligands (APLs) which seek to modulate the downstream signaling outcomes. However, dynamic links between modified peptide positions and distant residues have until yet been ill resolved. Using an integrative approach combining crystallographic ensemble and single models with atomistic molecular dynamics simulations and correlational analysis, we have established an approach that allows us to identify coupled dynamics between spatially distant residues at the pMHC interface. Furthermore, we constructed a network encoding the inter-residue couplings observed throughout the simulations. This computational workflow corroborates well with experimental data and leads to novel insights regarding the differential immunogenicity of the closely related peptides analyzed in this study. Ultimately, we present an intuitive and comprehensive strategy for decoding the linked dynamics at the pMHC interface allowing for mechanistic insights into the biophysical bases governing immunogenicity.

**One Sentence Summary:** The dynamics at the pMHC interface can be encoded as a biophysically relevant network to yield molecular insights into immunogenicity

## INTRODUCTION

The binding and recognition of a cognate class I major histocompatibility complex presenting a peptide antigen (pMHC) by a T cell receptor (TCR) is a critical step in the induction of CD8^+^ cytotoxic T lymphocyte (CTL) responses (*1*). Class I pMHCs are exposed on the surface of all nucleated cells, each a trimer consisting of a heavy chain, β_2_-microglobulin (β_2_m), and a peptide bound within a peptide-binding cleft created by the α1 and α2 domains of the heavy chain. The surface formed by the solvent-exposed residues of the presented peptide and the α1 and α2 helixes create the interface for TCR engagement (*2*). Given the essential nature of the pMHC/TCR interaction in the elicitation of CD8^+^ CTL activity, considerable effort has been invested in understanding the mechanisms of antigen processing and presentation (*3*), determining the structural and biophysical principles governing recognition (*1*, *4*), and designing altered peptide ligands (APLs) to modulate the immune response (*5*).

Structurally, the formation of a complementary surface between the TCR and pMHC is crucial for stable binding (*6*, *7*). The presented peptide antigen (*8*, *9*), which can be derived from oncogenic, autogenic, or pathogenic, including viral, sources, and the identity of the heavy chain allele (*10–13*) contribute to the likelihood of TCR recognition (*1*, *14*). Consequently, various pMHC properties have been used as predictors of immunogenicity, such as the thermal and kinetic stability of pMHCs (*15–22*). Additionally, the sequence of the TCR complementarity-determining regions (CDRs) restricts the population of stimulatory pMHC molecules for a specific TCR (*23*, *24*). TCRs are described as adopting a “canonical” diagonal docking topology when binding to pMHCs, and changes to the binding geometry can result in differential downstream responses (*25*, *26*). Regarding biophysical interaction properties, it is challenging to devise a general rule that applies to all immunogenic pMHC/TCR pairings. Kinetically, the importance of specific association rates, dissociation rates, or the resulting dwell time differs between systems (*27–32*). The thermodynamics of different stimulatory pairings can be driven by a combination of enthalpic and entropic contributions, as opposed to the historical model which dictated that generally unfavorable entropic effects were overcome by favorable enthalpic interactions (*13*, *33–44*). Furthermore, other factors may also affect CD8^+^ CTL activation, such as the successful engagement of CD8 (*45*, *46*) and CD3 (*47*) co-receptors or the presence of catch bonds at the TCR-pMHC interface, a requirement for the mechanosensing model of TCR recognition (*48–52*). In practice, TCR-pMHC affinity generally correlates with the induction of CTL responses (*27*, *44*, *45*, *53–62*). While exceptions to this rule exist, they typically violate other proposed requirements such as the canonical docking topology (*26*) or the presence of catch bonds (*63*).

In this study, we focused on a set of four highly related H-2D^b^-restricted peptides as a highly characterized model system to demonstrate the utility of our methods (**Table 1**). Within the context of murine lymphocytic choriomeningitis virus (LCMV) infection, a strong CTL response is induced against at least three immunodominant peptides (*64–67*). 50% of this response will be targeted towards the nonameric epitope gp33 (KAVYNFATM) derived from residues 33-42 of the viral glycoprotein (*68*, *69*). In this context, the sidechains of peptide residues p4Y and p6F are positioned centrally at the TCR-pMHC interface, engaging the CDR loops (*70–73*). Moreover, residue p1K is also crucial for P14 TCR recognition specifically (*72*). The selective pressure applied by CD8^+^ CTL activity promotes the emergence of common immune escape LCMV variants, which primarily manifest as single, relatively conserved, amino acid substitutions of p3V, p4Y, and p6F (*67*, *74*, *75*). One such peptide variant, in which p4Y is mutated to phenylalanine (Y4F; KAV**<u>F</u>**NFATM), escapes recognition by altering the respective TCR contact (*71*, *73*). The two other peptides investigated in this study are proline-APLs, where p3V has been substituted with a proline in gp33 (V3P; KA**<u>P</u>**YNFATM) and Y4F (PF; KA**<u>PF</u>**NFATM). Using circular dichroism and surface plasmon resonance measurements (*62*), we have previously demonstrated that the V3P modification of gp33 and Y4F enhances the thermal stability of the pMHC, improves its 3D affinity for the P14 TCR, and promotes TCR internalization (**Table 1**). Importantly, the p3P substitution does not significantly affect the overall pMHC conformation at the interface. Simultaneously, the ternary pMHC complex is stabilized through CH-π interactions between p3P and Y159 (*62*, *76*), a residue which is nearly perfectly conserved among human and murine alleles (*77*, *78*). Proline-APLs display enhanced immunogenicity compared to their wild-type counterparts in various systems (*62*, *76*, *79–82*), and CD8^+^ CTL cross-reactivity is induced against wild-type peptides following APL vaccination (*62*, *79*, *81*). Crystallographic structures of H-2D^b^-restricted gp33, Y4F, V3P, and PF highlight that specific H-2D^b^ (R62, E163, H155) and peptide (p1K, p6F) residues adopt a different conformation after p3P substitution. The altered conformation seemingly primes the pMHC for TCR engagement as the sidechain rotamers closely match those of the P14 TCR-bound structures. Despite the extensive characterization of this set of peptides, a mechanistic insight into the conformational coupling of these residues remains elusive.

**Table 1.**
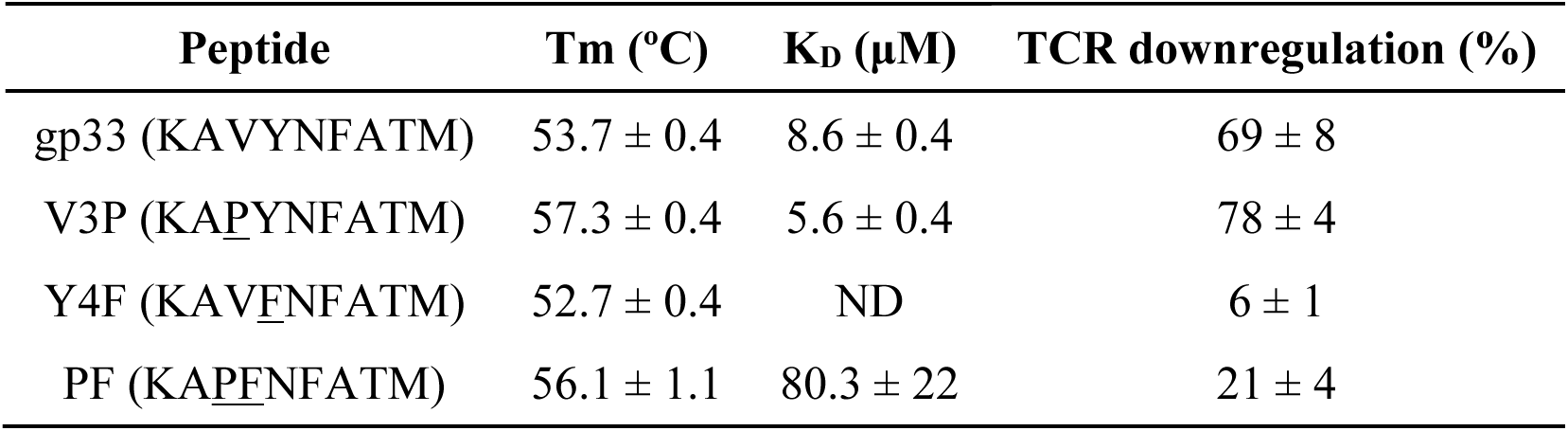
MHC stability and P14 affinity to MHC binding gp33 variants. . The thermal stability of the pMHC was measured with circular dichroism (CD); T_m_ is the temperature where 50% of the molecules are denatured. The dissociation constant (K_D_) was determined from steady state surface plasmon resonance (SPR) data. TCR downregulation on P14 T cells exposed to 10^-8^ M peptide-pulsed RMA cells. Values in the table have been published previously (*62*).

Given the marked reduction in the permitted and favored ranges of dihedral angles of proline compared to other amino acids, we set out to explore the ways in which the dynamics at the TCR-pMHC interface were altered following its introduction. Furthermore, other studies have begun to highlight the importance of localized residue flexibility, peptide dynamics, conformational rearrangements, and the overall pMHC free energy landscape in influencing TCR affinity and antigen presentation (*2*, *83–94*). As such, we used an integrative approach, combining crystallographic ensemble refinement with molecular dynamics (MD) simulations and correlational analysis, to uncover dynamic relationships between the p3P substitution and other residues at the TCR-pMHC interface. We further expand on these analyses by encoding the observed correlational couplings as a network to intuitively and holistically explore the dynamics at the pMHC interface within the context of our chosen peptides.

## RESULTS

### Similarities in packing between crystals allow for informative ensemble refinement

As the ensemble refinement is run using restraints derived from X-ray data (*95*), it was important to analyze and evaluate the potential effects and artifacts introduced by the crystal geometry of each complex. The published structures (*62*, *71*, *73*) of H-2D^b^/gp33, H-2D^b^/Y4F, and H-2D^b^/PF reveal similarities in crystal packing. Specifically, the three TCR-unbound crystals are in the same space group (**Table S1**). An analysis of residues involved in polar interactions less than 4.0 Å apart (**Table S2**) and visual inspection of the peptide-bound α1 and α2 domains further reveal the full extent of crystal contact conservation (**Fig. S1**). All complex copies in each crystal form extensive contacts at the distal ends of the α1 and α2 helixes towards the C-terminal end of the peptide-binding groove. Two copies of each complex also form contacts through the N-terminal half of the α1 helix. Some crystal contacts are also found along the flexible loops connecting the antiparallel strands of the β-sheet that forms the base of the peptide cleft. The crystal packing around the peptides is also conserved between crystals, with two copies of each complex displaying more freedom around either p4Y/F or p6F (**Fig. S1**). It should be noted that p4 in all unbound complexes was identified as a contact. However, we still expect to be able to gain some biological insight, beyond crystal artifacts, on the dynamics at this position as different p4Y/F rotamers are observed in each copy of each crystal. The arrangement of the peptide contacts does, however, hinder our ability to assess concerted conformational transitions observed in the peptide. We decided to consider all crystallographic copies of the peptide when assessing the conformational space occupied by each complex due to the differences in crystal contacts between copies.

In terms of the other complexes, H-2D^b^/V3P is not found in the same space group as the other unbound pMHC complexes (**Table S1**), and only one copy is found in the asymmetric unit cell; thus, no alternative contact geometries were sampled. Fortunately, similar regions of the complex are involved in crystal contacts compared to the other TCR-unbound pMHC complexes (**Table S2; Fig. S1**). For completeness, H-2D^b^/V3P was included in the ensemble refinement analyses. The TCR-bound complexes H-2D^b^/gp33/P14, H-2D^b^/V3P/P14, and H-2D^b^/PF/P14 also revealed conservation of regions involved in crystal packing (**Tables S1, S3**). As expected, no contacts involving the peptides were identified, and contacts at the interface were limited by the presence of the P14 TCR.

Given the similarities in crystal packing identified between the different published structures, we proceeded with the ensemble refinement of the previously published structures (**Table S1**), which in most cases led to a slight improvement in the R-factors.

### The p3P substitution alters the conformational dynamics of MHC-restricted peptides

The published crystal structures of TCR-unbound pMHC complexes H-2D^b^/gp33, H-2D^b^/V3P, H-2D^b^/Y4F, and H-2D^b^/PF have been previously compared with the TCR/pMHC complexes (*62*). The peptide conformation in H-2D^b^/Y4F closely resembles that of the H-2D^b^/gp33 peptide. While, in H-2D^b^/V3P and H-2D^b^/PF, the p6F and p1K sidechains are observed in a different rotameric state, presumably due to the p3P substitution, leading to an overall conformation resembling the TCR-bound form (**Fig. 1A**). The binding of the P14 TCR results in conformational changes in the p4Y/F and p6F sidechains of gp33, V3P, and PF. Additionally, the sidechain of p1K in gp33 moves towards the N-terminal while it is already correctly positioned for P14 TCR engagement in V3P and PF (**Fig. 1A**).

**Fig 1.**
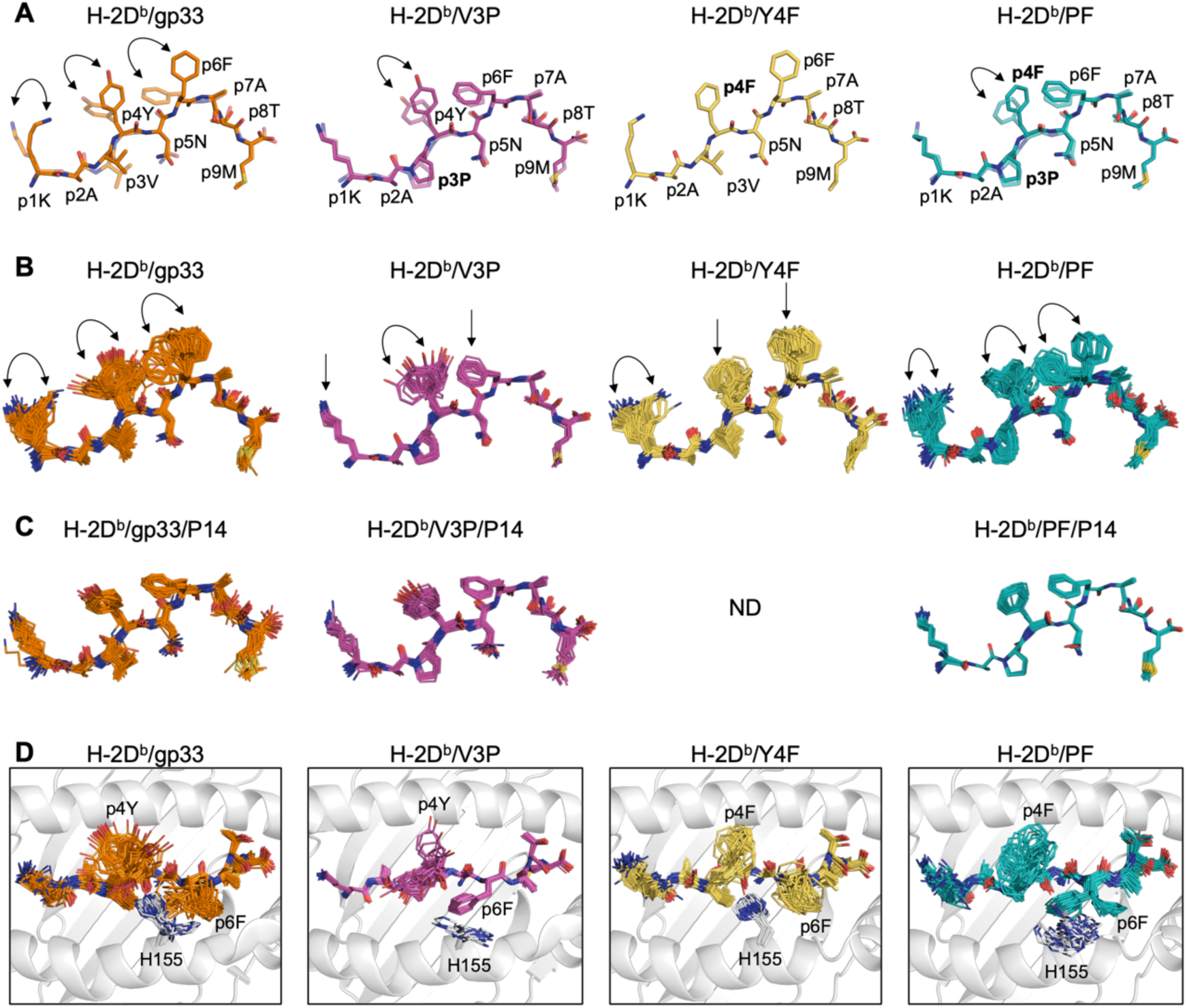
The conformational populations of gp33, V3P, Y4F, and PF peptides, calculated by ensemble refinement, unveil dynamic information from crystal structures. Stick representation of the peptides bound to the MHC. **A** Full-color peptides represent the peptide conformation before TCR binding in the crystallographic structure. In contrast, transparent peptides represent the conformation that the peptides assume after TCR binding. No crystallographic data are available for H-2D^b^/Y4F/P14. The arrows indicate the changes in sidechain conformation. **B** Polymorphic peptide conformations of gp33, V3P, Y4F, and PF of the TCR-unbound pMHC complex calculated by ensemble refinement. The arrows indicate variations in the conformation of the sidechain. **C** After TCR binding, the peptides rigidify, and all peptide ensembles assume the same restricted conformation. No crystallographic data are available for H-2D^b^/Y4F/P14. **D** Concerted and opposing motions between H155 and p6F sidechains were observed in the TCR-unbound pMHC complexes, particularly in H-2D^b^/gp33 and H-2D^b^/PF. H-2D^b^ (white) is shown in cartoon representation and the H155 sidechain is shown in stick representation.

Ensemble refinement allowed us to gain insights into the polymorphic characteristics of peptides (**Figs. 1B, 1C**). Residues p1K, p4Y/F, and p6F in the gp33 and PF peptides as well as residue p4Y in the V3P peptide display the greatest structural fluctuations. These sidechains explore different conformations, including those assumed after TCR binding in the crystal structure. In contrast, the p4F and p6F sidechains of the immune escape Y4F peptide ensemble do not sample a conformation similar to the TCR-bound conformation observed in other peptides, suggesting more restricted structural dynamics (**Fig. 1B**). Therefore, it appears that introducing p3P to the Y4F peptide in PF enables the peptide to sample a wider range of sidechain conformations, some of which are permissive to P14 TCR engagement, including the “prohibited” p6 flip. Our ensemble analyses imply that the lack of downward rotation of p4F in the Y4F peptide could be essential for hindering the formation of the TCR/pMHC complex. The TCR-bound pMHC ensembles display a decreased diversity of adopted peptide conformations, implying that TCR binding rigidifies the restricted peptide, stabilizing its conformation and reducing the accessible conformational space (**Fig. 1C**).

### Mechanistic insights into proline-altered dynamic areas are hindered by crystal contacts

In addition to comparing the presented peptides, we comprehensively analyzed the ensemble dynamics at the pMHC-TCR interface. The RMSF profiles of each ensemble copy were compared (**Figs. S2, S3**), in addition to various sidechain ensembles of MHC and TCR residues of interest (**Figs. 1D, S4-S6**). Most notably, a high degree of concerted motion was observed between the H155 and p6F sidechains, facilitated by their proximity in the 3D structure (**Fig. 1D**). While the sidechains of these two residues alternate between two main conformations in H-2D^b^/gp33, these same residues adopt only one conformation in H-2D^b^/Y4F. The observed inability of p6F to transition between sidechain rotamers, presumably due to steric hindrance induced by the H155 sidechain, may explain the lack of recognition by the P14 TCR.

In contrast, the introduction of a proline in H-2D^b^/PF, recognized by P14 (**Table 1**), allows the formation of two main conformations for both p6F and H155 (**Fig. 1D**). This observation suggests a plausible mechanism by which the conformational sampling of the p6F sidechain is altered through the transmission of dynamics from p3P through the α2 helix and finally to p6F and H155. Interestingly, substituting p3V to a proline in gp33 is associated with only one conformation for p6F in H-2D^b^/V3P (**Fig. 1D**), leading to substantially increased recognition by the P14 TCR (*62*).

Furthermore, the TCR-unbound complexes displayed a diverse distribution of RMSF values along the α1 and α2 domains (**Fig. S2**). Increased disorder in loop regions was observed, particularly around residues 13-21 and 38-44. The introduction of p3P led to the rigidification of H-2D^b^, namely in the α1 and α2 helix residues surrounding p3P, but also in some loop regions. Similar overall rigidification of proline-APL pMHCs has been shown in other studies (*76*). As expected (*96*, *97*), the dynamics at the TCR-pMHC interface are considerably reduced after P14 TCR binding (**Fig. S3**). While a relative reduction in dynamics in regions surrounding p3 is still observed, no relationship between dynamics at the TCR interface and affinity was identified, partly due to the lack of an available H-2D^b^/Y4F/P14 structure. H155 assumes a stable conformation in all ternary complexes (**Fig. S4**). Other residues, including E63, K66, E163, R75, R79, and K146, exhibit higher conformational variability in H-2D^b^/gp33/P14 and H-2D^b^/V3P/P14 complexes compared to H-2D^b^/PF/P14 (**Figs. S5, S6**). Unfortunately, the confirmation of the proposed transmission of dynamics and the overall reliability of the dynamic analysis were hindered by the prevalence of crystal contacts, primarily in the unbound pMHC ensembles, along the α1 and α2 helixes (**Tables S2, S3; Fig. S1**).

### MD simulations of the pMHC molecules agree with the crystallographic data

We ran unbiased MD simulations of each pMHC copy in the TCR-unbound crystallographic models to account for potential crystal artifacts in the ensembles. It must be noted that only the comparison between H-2D^b^/gp33 and H-2D^b^/V3P or H-2D^b^/Y4F and H-2D^b^/PF is stringently valid, as we only included an additional set of simulations where p3V was mutated to p3P or vice versa. Therefore, comparisons between H-2D^b^/gp33 and H-2D^b^/Y4F or H-2D^b^/V3P and H-2D^b^/PF do not adequately control for initial conFiguration biases. As a result, we cannot assess the dynamic effect of the p4Y to p4F mutation. We were, however, able to draw conclusions on the effects of p3P-altered ligands on antigen presentation and thus immunogenicity, given that the peptide conformations throughout each trajectory are comparable with the crystallographic ensembles and single models (**Figs. 2, S7**). We observed that the backbone dynamics and conformations are highly constrained and similar between all trajectories and that p1K, p4Y/F, and p6F sample the largest range of conformational space with their sidechains (**Figs. 2C, S7C**). In the case of the gp33 and Y4F peptides, the p3V sidechain is also relatively more mobile than the rest of the peptide. Critically, the per-residue RMSF data for p3V/P and p6F are the strongest correlates of pMHC immunogenicity (**Fig. S8**).

**Fig 2.**
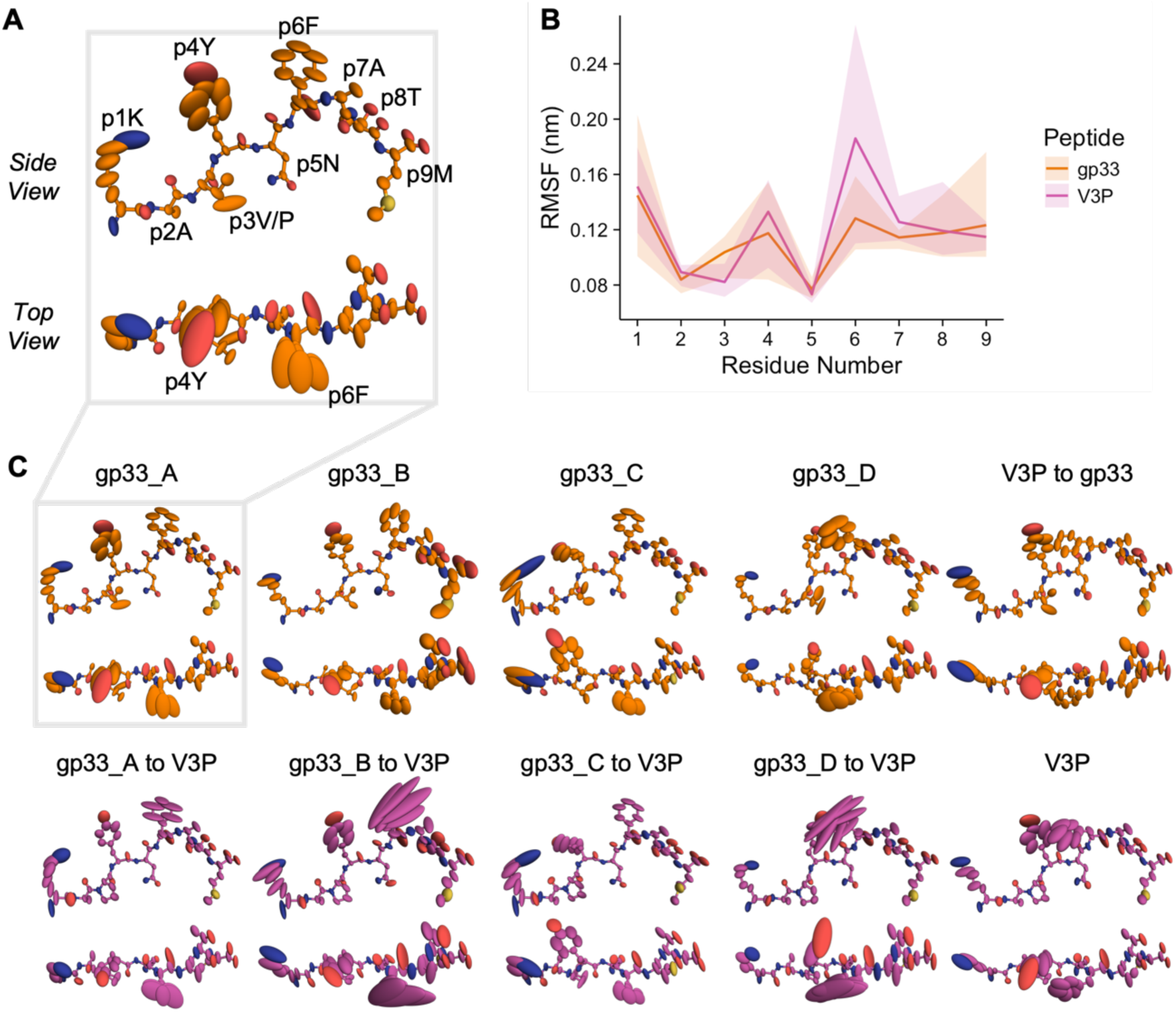
Overview of the conformational sampling of the gp33 and V3P peptides throughout the MD trajectories reveals that proline substitution leads to rigidification at p3 and increased dynamics at p6F. Stick and ellipsoid representation of MHC-bound gp33 (orange) and V3P (purple). Ellipsoids are scaled and shaped according to the anisotropic atomic temperature factors derived from each trajectory and mapped to their respective crystallographic models. **A** Residue labeling of the side and top views of the peptide as reference. **B** Average per-residue RMSF values over all trajectories for gp33 and V3P. The colored transparent envelopes depict the full range of observed trajectory RMSF values for each peptide. **C** Conformational sampling of the peptides observed in each trajectory. Each column represents a set of paired simulations. One derived from the initial crystallographic model and the other derived from an *in silico* mutation between p3V and p3P.

Two main insights are gained from these trajectories regarding the observed biophysical and functional effects of the p3P substitution. Firstly, the overall dynamics, both backbone and sidechain, are remarkably comparable between p3V and p3P trajectories derived from the same crystallographic pMHC copy (**Figs. 2C, S7C**). This may explain why overall pMHC melting temperature remains above 50 °C, regardless of peptide (**Table 1**). Moreover, these similarities provide a basis for the observed P14 TCR cross-reactivity upon proline-APL vaccination via molecular mimicry (*62*). Secondly, although p3P is consistently less dynamic than p3V, in some cases p4Y/F and p6F display greater RMSF values in the presence of p3P, which is observed both visually (**Figs. 2C, S7C**) and in the per-residue RMSF data (**Figs. 2B, S7B**). This was further analyzed statistically by considering each corresponding p3V and p3P trajectory as paired observations and performing an exact Wilcoxon signed-rank test on the per-residue RMSF data of p3V/P, p4Y/F, and p6F from all simulations. The uncorrected p-values of 0.013, 0.047, and 0.095 were obtained, with the alternative hypotheses that p3P is less dynamic than p4V, p4Y/F is more dynamic in p3P trajectories, and p6F is more dynamic in p3P trajectories, respectively. The link between p3V/P and p6F dynamics cannot be readily explained by altered conformational sampling of the peptide backbone (**Figs. S9-S11**). These observations, coupled with the previously published structures and presented ensemble refinements, may be sufficient to explain the differences in 3D TCR affinity and downstream P14 T cell activation (*62*). Importantly, these MD trajectories, in which the influence of crystal contacts has been minimized, agree with the crystallographic ensemble and single model data.

As with the ensemble refinement, the H-2D^b^ dynamics were also assessed through visual inspection (**Figs. S12, S13**) and RMSF profile comparison (**Fig. S14)**. H155 displayed increased conformational sampling, particularly in trajectories where the p6F sidechain also sampled a greater range of conformations (**Figs. 2C, S7C**). Visually, E163, which is positioned near p3, was consistently less dynamic in the presence of p3P. On the other hand, no dynamic relationship between R62 mobility and p3V to p3P mutation could be observed. It should be noted that the latter does not rule out the presence of an altered conformation upon introducing p3P (*62*). The other residues of interest (E63, K66, R75, R79, K146) did not display consistent sampling differences when comparing the paired trajectories (**Fig. S13**). This aligns with observations that the dynamics of these residues at the TCR-pMHC interface throughout the ensemble refinements did not relate to overall interaction affinity (**Figs. S4-S6**). One difference between the ensemble refinements and MD trajectories was identified between the overall per-residue RMSF profiles of H-2D^b^ (**Fig. S14**). Specifically, the relative differences in H-2D^b^ dynamics in loop regions, such as the 38-44 and 86-93 loops, do not concur with the ensemble refinements (**Fig. S2**). It cannot easily be ascertained whether this deviation is due to crystal artifacts, differential crystal packing, forcefield artifacts, stochastic assignment of initial velocities, or some other effect.

### Correlational analyses unveil dynamically linked clusters of residues

Correlational substructure is visible in the pairwise Spearman rank correlation matrix of the per-residue RMSF data (**Fig. 3A**). Structural visualization is critical to identify groups of residues that cluster closely together in real space. Thus, the significant correlations of each peptide residue were mapped to the structure of H-2D^b^/gp33 (**Fig. 3C**). Consequently, the chance of deriving conclusions from false positives and spurious correlations is also reduced, as is likely the case between p1K and R121. While correlational analyses cannot typically infer directionality, our RMSF data is related over paired simulations, therefore, it seems likely that observed correlations will be partly due to the introduction of a proline in the region surrounding p3V/P.

**Fig 3.**
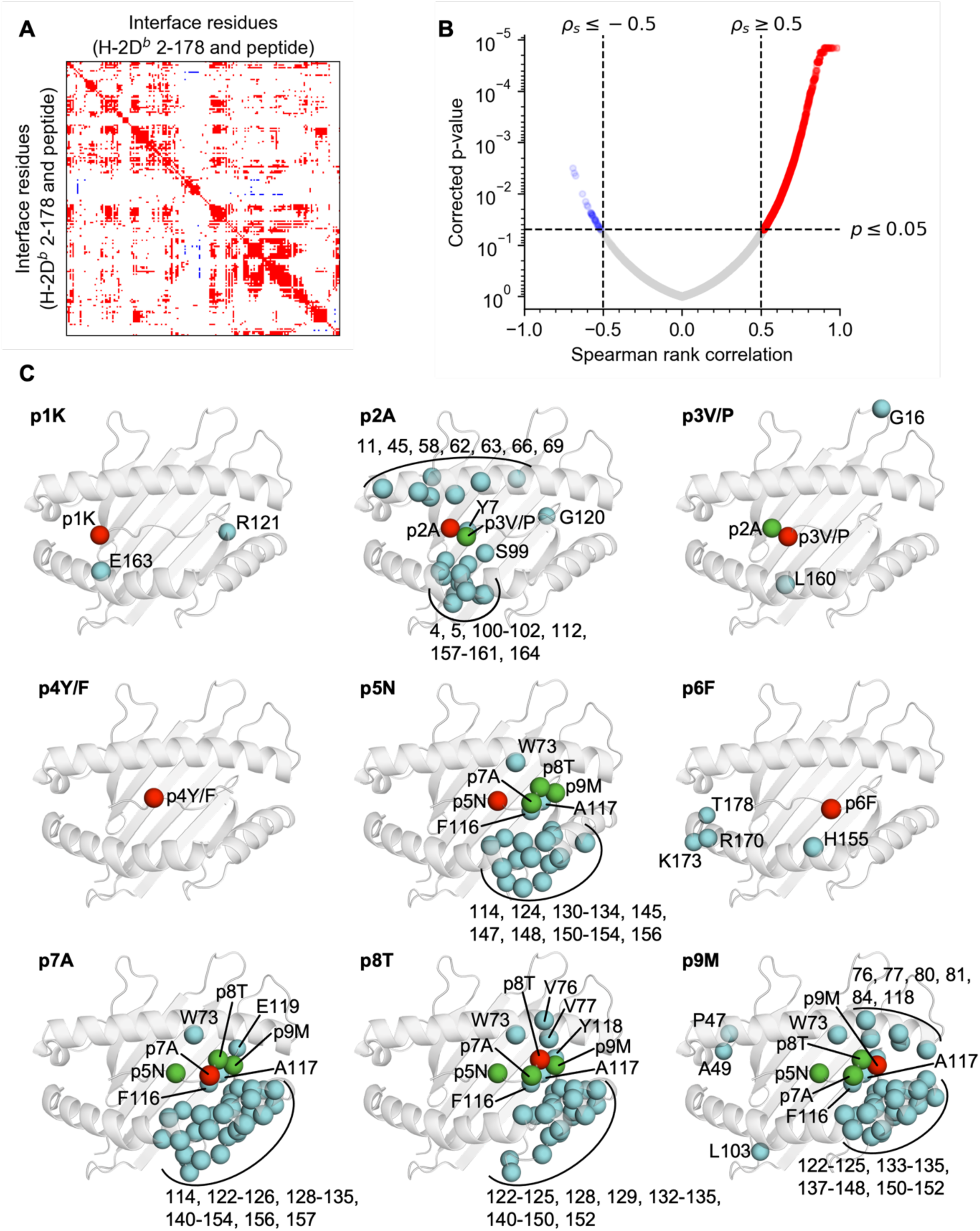
Mapping all significant pairwise correlates of peptide per-residue RMSF data onto the structure of H-2D^b^/gp33 yields spatially restricted clusters of dynamically similar residues. **A** Spearman rank correlation matrix depicting significant positive (red) and negative (blue) pairwise correlation between per-residue RMSF values at the interface. Nonsignificant values have been set to 0 (white). **B** Volcano plot of Spearman rank correlation and corrected p-values used to determine significant pairwise correlations. Note that the threshold of |ρ_s_| < 0.5 does not filter out any points in this dataset, when used in combination with a p < 0.05 threshold. **C** Significant pairwise correlates of each peptide residue (red) shown within their structural context. Correlates are colored depending on whether the residue is in the heavy chain (cyan) or the peptide (green). The backbone of H-2D^b^/gp33 (white) is shown in cartoon representation as reference.

It should be noted that no significant correlations were identified for p4Y/F (**Fig. 3C**). This may be explained by the solvent-exposed nature of the sidechain, resulting in considerably higher dynamics and freedom compared to surrounding residues, hindering strong correlations. Alternatively, it could be explained by the presently used simulation scheme, where p3P and p3V trajectories can be paired accordingly. We could investigate the dynamic effects of p4 mutations at the interface by also generating paired p4Y and p4F trajectories. Although it seems unlikely the results would be as striking compared to those of the p3P substitution given the solvent-exposed nature of the residue. In other words, we cannot adequately explain the lack of correlation with p4Y/F. As an addendum, if this lack of dynamic coupling is biophysically representative, it would suggest that the p3P subverts Y4F-dependent immune evasion by enhancing P14 TCR affinity through p6F, rather than altering the dynamics at p4Y/F.

On the other hand, the dynamics at the main anchor residues for H-2D^b^, p5N and p9M (*79*, *98*) correlated strongly with numerous residues of H-2D^b^. Although coupled residues are also present in the α1 helix and at the base of the peptide cleft, the largest cluster is found in the α2 helix. This relationship can be extended to residues p7A and p8T, which, despite not being anchor residues, also show extensive correlation with the same MHC regions. In the case of H-2D^b^/gp33 and related peptides, p2A and p3V/P act as secondary anchor residues (*44*, *71*, *73*). Our data corroborates this as p2A and p3V/P are correlated, and p2A is further linked with multiple residues along both α1 and α2 helixes. Notably, the dynamics of R62, directly, and E163, through proximity, are associated with those of p2A and p3V/P. These H-2D^b^ residues were identified to have altered conformations after the introduction of p3P and form important contacts with the P14 TCR (*62*). H155, another critical residue for TCR contacts, correlates with p6F, which closely matches our observations from the ensemble refinement (**Fig. 1D**). While H155 and p6F do not strictly cluster with the directly surrounding peptide and heavy chain residues, changes to their backbone dynamics will bias the propensity for sampling specific sidechain conformations (*99*). Additionally, p3P substitutions are stabilized by Y159 (*62*, *76*), and our data shows that the dynamics for residues 157-161, p2A, and p3V/P are related.

The coupling of residue dynamics can also be investigated using agglomerative hierarchical clustering on rank correlation distance of the per-residue RMSF data (**Fig. S16**), instead of assessing significant pairwise Spearman rank correlations. This yields very similar results, in which p3V/P and p2A cluster with Y159 on the α2 helix, H155 and p6F cluster with each other, and the rest of the α2 helix clusters with p5N and p7A (**Fig. S17**). These findings begin to paint a picture of how the p3P substitution can affect the conformational sampling of p6F, and ultimately whether the pMHC can be recognized by the P14 TCR. In particular, we propose that the altered dynamics at p3P are predominantly transmitted via the α2 helix to H155 and p6F. While promising, the methods presented thus far adopt a relatively reductionist approach to investigating pMHC dynamics at the interface. It would, therefore, prove valuable to encode the correlation of residue dynamics as a network to probe interface dynamics holistically using graph theory.

### Dynamic correlations can be used to generate a biophysically relevant network

Networks were constructed by defining each residue as a node in the network and deriving the edge weights from the matrix of significant Spearman rank correlations (**Fig. 3A**). We devised three path-based validation metrics to validate whether networks derived from the correlation data yielded biophysically relevant observations: F_1_, F_2_, and F_3_ (**Extended Methods**). F_1_ accounts for uncharacteristically distant couplings observed in all shortest paths. This is achieved by calculating the proportion of shortest paths in the network where any edge travels a larger through-space distance than the initial distance. F_2_ and F_3_ provide related measures of the proportion of shortest paths that converge directly to their respective target nodes. They measure the proportion of shortest paths where the through-space distance increases at any node on the path relative to all previous nodes or the initial node respectively. F_3_ was designed to partially address the issue of edges, which diverge from the target node yet display high edge betweenness, being counted multiple times when calculating F_2_. These metrics and the proportion of nodes in the largest connected component were calculated for each network derived from a 2D scan of network construction parameters (**Fig. S18**). Based on a compromise between minimizing F_1_ and maximizing the connected residues in the network, we constructed the network by only considering significantly correlated residues whose inter-residue C_α_ distance was less than 10 Å (**Fig. 4**). We decided not to incorporate a distance-based cost into the edge weight metric to reduce biases introduced in network construction. Furthermore, the validation metrics of networks with a distance threshold of 10 Å only improved marginally when including distance in the edge weight (**Fig. S18**).

**Fig 4.**
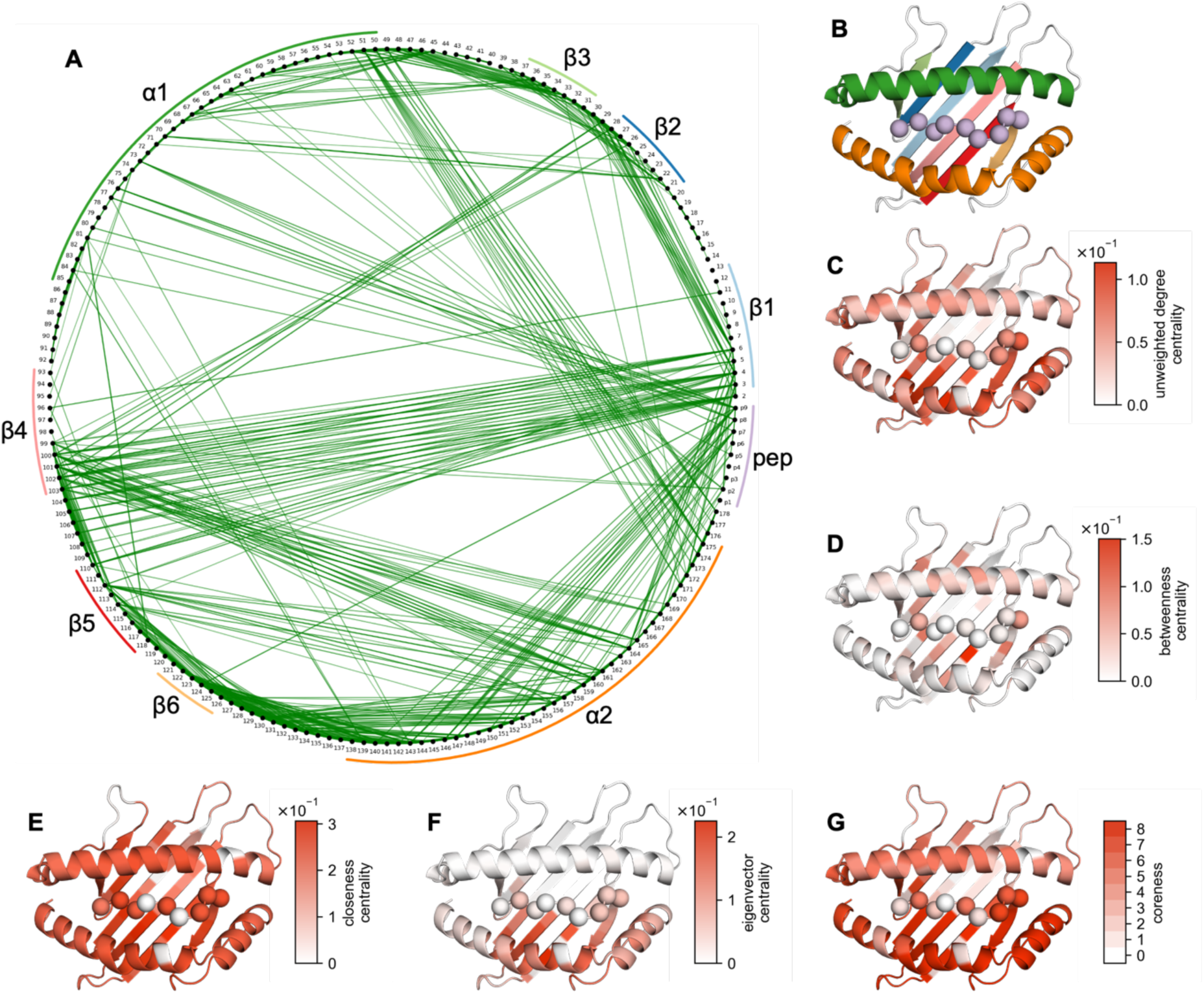
A biophysically relevant network can be derived from the correlation matrix revealing residues playing a central role in interface dynamics. **A** The network encoding interface dynamics, where each node (black) represents an interface residue and edges (green) are significant correlations between residues whose alpha-carbon distance does not surpass 10 Å. Edge transparency and width are scaled according to |ρ_s_|. Secondary structure features of the interface have been labelled. **B** Secondary structure features of the H-2D^b^/peptide interface colored according to the same label colors as in **A**. Various informative network centrality measures can be mapped to structure: **C** unweighted degree centrality, **D** betweenness centrality, **E** closeness centrality, **F** eigenvector centrality, and **G** coreness. Plots showing centrality measures per residue are provided (**Fig S19**).

An analysis of the topological properties of our biophysically relevant network produced expected observations and novel insights regarding this set of H-2D^b^-restricted gp33-associated peptides (**Figs. 4, S19**). Importantly, edges are observed between residues in neighboring secondary structure features as expected (**Figs. 4A, 4B**). Moreover, not every neighboring residue is linked, highlighting the discriminatory capacity of the network in identifying dynamically coupled residues through correlational metrics rather than reproducing a simple pairwise distance matrix. When considering the unweighted degree centrality (**Fig. 4C**) of each node, p2A, p5N, and p9M are amongst the most connected peptide residues paralleling their role as anchor positions (*44*, *71*, *73*, *79*, *98*). L114 displays the highest betweenness centrality (**Fig. 4D**), indicating it sits on the largest proportion of shortest paths in the network. It may, therefore, play a likely central role in transmitting dynamics throughout the interface. In class I pMHCs, residue 114 forms part of the F pocket for peptide binding and is found in a region whose dynamics have been linked to complex stability and peptide loading efficiency (*70*, *73*, *90*, *100–102*). In terms of the closeness centrality of each residue (**Fig. 4E**), most of the complex is similarly central, aside from loop regions and nodes outside of the largest connected component, as would be expected for a globular protein. Both the eigenvector centrality (**Fig. 4F**), which favors high degree connectivities with other highly connected nodes, and k-coreness (**Fig. 4G**) of each residue identify the α2 helix as forming an influential and core part of the network. It is not immediately evident whether the latter observation is due to conserved dynamic patterns at the pMHC interface, a system-specific result of analyzing paired p3V and p3P trajectories, or a combination thereof. Nevertheless, these findings reflect the ability of our constructed network to capture the biophysically relevant coupled dynamics at the pMHC interface. For completeness, a network derived directly from the correlation matrix, without any modification or transformation, and its corresponding topological analysis is also available (**Figs. S20, S21**).

### Network analysis unveils long-range coupling between residues p3 and p6

Using a network to describe pMHC dynamics opens the door to a suite of potential network analyses to probe the structural and biophysical properties of the pMHC interface. We present some examples such as shortest path analysis and Louvain community detection (**Fig. 5**). By mapping all shortest paths originating from p3 onto the structure of H-2D^b^/gp33, we observe that the dynamics at p3V/P are correlated to those of p2A and L160 before being transmitted across the whole interface. This transmission occurs primarily through the α1 and α2 helixes (**Fig. 5A**). A connected subgraph of this network (**Fig. 5B**) shows that the shortest path of dynamic coupling from p3 to p6 is through the α2 helix. Namely, p3P transfers its dynamics to L160, likely via its interaction with Y159 (*62*, *76*), before this propagates to the residues surrounding p6F and H155. It should be noted that p6F and H155 are not part of the largest connected component of the biophysically relevant network. However, altered backbone angle distributions could explain the increased conformational sampling of the p6F sidechain (*99*). In fact, the backbone and Ψ angles of p6F and the Φ angle of p7A display some of the largest deviations in conformational distribution after the introduction of p3P (**Figs. S9F, S10F, S11F**). Furthermore, community detection through the Louvain algorithm corroborates the dynamic coupling between p3V/P and the α2 helix, as seen in the most frequent solution obtained in 11.2% of iterations (**Fig. 5C**), and community co-occurrence frequency of each residue with p3P (**Fig. 5D**). The latter was calculated as we postulated that the low recurrence of the most frequent solution was due to several residues on community fringes that would cluster inconsistently rather than fully or mostly unique community structures with each iteration. This was confirmed by the high community co-occurrence frequency of p1K, p2A, p3V/P, and the region of the α2 helix containing Y159 and L160.

**Fig 5.**
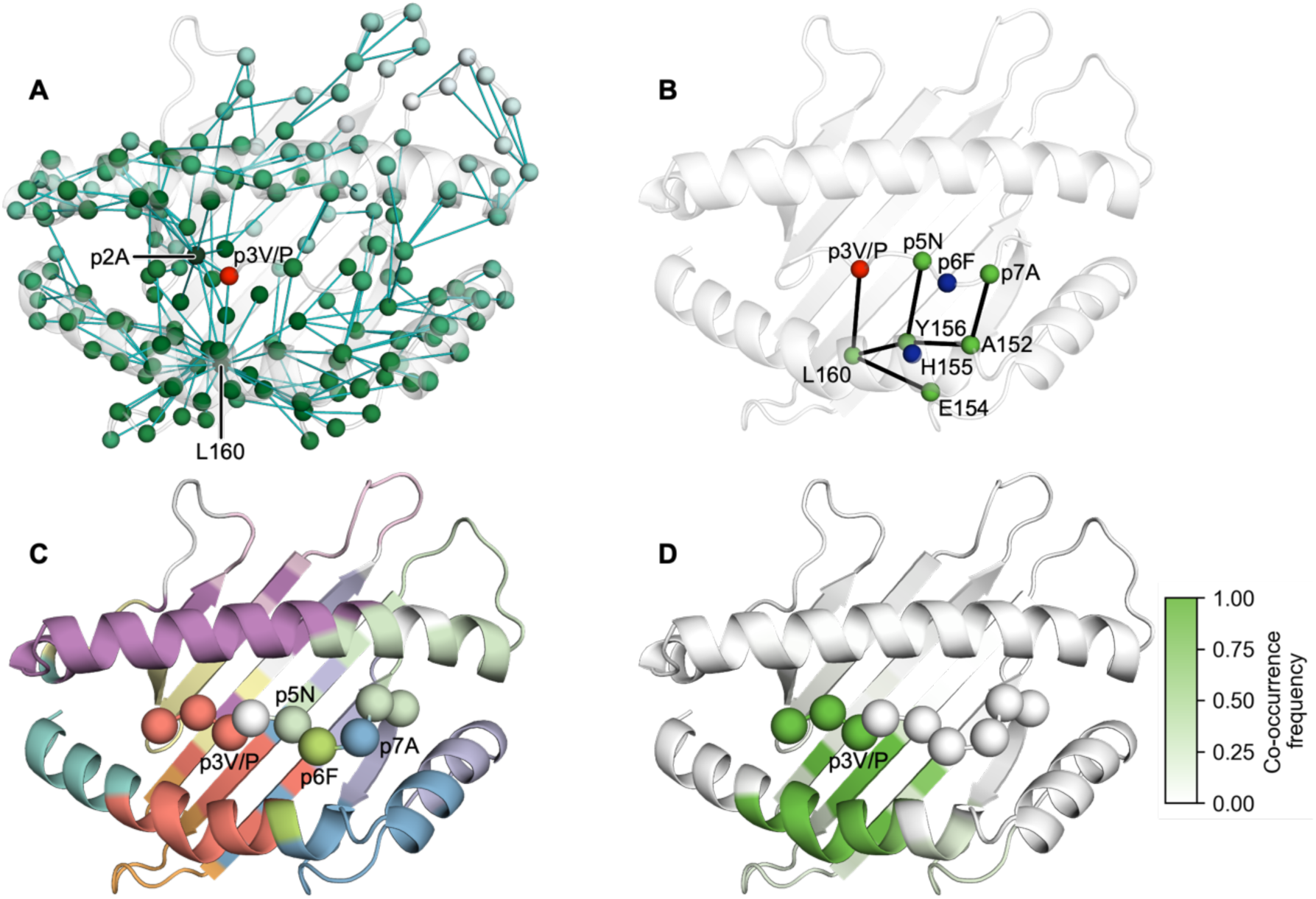
Shortest path analysis and Louvain community detection indicate that the dynamics of p3V/P are coupled through the α2 helix with the residues surrounding p6F and H155. **A** All shortest paths from p3V/P (red) to every other node in the largest connected component of the network. The dynamic signal of p3V to p3P mutation is firstly transmitted to p2A and L160 before propagating to the rest of the interface, primarily through the α2 helix. Residues are colored according to their shortest path distance from p3V/P, from green to white. Edges (teal) traversed by the shortest paths are shown. **B** Shortest paths from p3V/P to the residues neighboring p6F and H155. **C** Most frequent Louvain community detection solution obtained, representing 11.2% of solutions from 1 x 10^6^ iterations. Each community of residues has been depicted in a different color; residues that are not in any community are shown in white. The first three peptide residues cluster with the part of the α2 helix which neighbors communities containing p6F and p7A. **D** Community co-occurrence frequency of each residue with p3V/P over all Louvain community detection iterations. This demonstrates that p3V/P consistently clusters with residues on the α2 helix.

## DISCUSSION

The immunostimulatory potential of class I pMHC molecules is governed by a combination of the complex’s composition, structure, stability, and dynamics (*1*). Critically, the surface properties and dynamics at the TCR interface are altered by the loading of different peptides in the pMHC peptide-binding cleft. These dynamics extend beyond local fluctuations, resulting in multiple conformational states within the complexes (*42*, *89*, *97*, *103*)

In the context of both the unbound pMHC and P14 TCR-bound pMHC, the H-2D^b^-restricted gp33, V3P, Y4F, and PF peptides have been extensively structurally, biophysically, and functionally characterized by our group (*22*, *44*, *62*, *71–73*, *104*, *105*). The free energy surface and conformational space inhabited by the peptides in H-2D^b^/gp33, H-2D^b^/V3P, H-2D^b^/gp33/P14, and H-2D^b^/V3P/P14 were also investigated using MD simulations (*106*). Our present results suggest that altered dynamics at the third peptide residue upon p3P substitution can result in long-range modulation of conformational sampling of p6F through dynamic coupling via the α2 helix. While this mechanistic insight has been missing for this system of peptides, the dynamic interplay between the peptide and surrounding α1 and α2 domains is well established (*2*, *83–92*). Despite our analyses being primarily centered on the unbound pMHC, they corroborate well with the existing structural and functional models, which indicate that proline-APLs prime the complex for engagement by the TCR (*62*). In particular, we propose that the increased sampling of TCR-permissive sidechain conformations of p6F, observed in the crystal structure, ensemble refinement, and MD trajectories, is facilitated through long-range dynamic coupling with p3P. This may explain why V3P and PF act as a superagonist or a partial agonist respectively, in terms of P14 T cell and endogenous CD8 T cell responses compared to the full agonist gp33 or the null immune escape peptide Y4F (*62*).

As mentioned, no correlational linkages were observed between p4Y/F and any other residues. However, it is not clear whether this is due to the MD simulation pairing scheme or the increased dynamics of the solvent-exposed p4Y/F sidechain. Furthermore, the mechanism driving Y4F-dependent immune escape seems to be caused by the altered electrostatic surface potential at the interface and loss of critical P14 TCR contacts (*62*, *71*, *73*), rather than the dynamic modulation of neighboring residues. The Y4F modification abolishes the possibility of establishing hydrogen bond interactions as well as hydrophobic interactions with the sidechain of the P14 residue R97β and additional hydrogen bond interactions with the H-2Db residue E163 and the TCR residue Y101α (*62*). As such, it seems unlikely that this observation detracts from our findings. Additionally, it remains to be seen whether these long-range dynamic couplings of interfacial residues are conserved among other proline-APL systems (*76*, *79*, *82*). Other MD and crystallographic studies of the H-2D^b^-restricted melanoma-associated proline-APL of gp100 and Trh4 demonstrated that the p3P substitution results in overall rigidification at the interface (*76*, *82*).

We have demonstrated that the present MD protocol aligns well with single model and ensemble crystallographic observations of peptide dynamics and specific H-2D^b^ residues that are key for TCR recognition. However, dynamic links were largely obscured by the presence of crystal contacts along the α1 and α2 helixes. Previous ensemble refinement studies of pMHC molecules circumvent the issue by only analyzing human class I pMHC ensembles with resolutions better than 3 Å that displayed the largest RMSFs at the interface (*84*). This approach cannot completely account for contacts and would reduce our agency in choosing which system to investigate. Serendipitously, the space groups and arrangements of crystal contacts of the H-2D^b^-restricted gp33, V3P, Y4F, and PF crystals allowed for the validation of MD trajectories to establish a reasonable simulation protocol. It may now prove interesting to adapt the combined MD and dynamical network analysis workflow to other proline-APLs, such as those derived from the H-2D^b^-restricted cancer-associated gp100 (*79*) or Trh4 (*80*, *82*) peptides. These were previously inaccessible for extensive ensemble refinement analyses due to their unfavorable crystallographic contact parameters.

Using the per-residue profiles of interface dynamics, we established an intuitive and systematic workflow capable of extracting and analyzing long-range coupled dynamics at the pMHC interface. The presented strategies were inspired by established analyses of MD simulations (*107*). In the literature, correlational analyses of protein dynamics typically assess the pairwise cross-correlation of atomic displacement vectors, sometimes only of the Cα atoms, over single MD trajectories to identify concerted motions (*86*, *108–110*). Instead, we analyzed the correlations in the magnitudes of the per-residue RMSF values over multiple paired trajectories. We were interested in identifying correlated differences in RMSF profiles after introducing a dynamic perturbation in the form of a p3P mutation, instead of the subtly different exercise of identifying concerted motions of atoms along a trajectory. The abstraction of protein dynamics on a per-residue basis was initially applied to address the challenge of appropriately aligning atomic indexes from different trajectories despite the presence of p3V/P mutations and, to a lesser extent, unmodeled termini. Subsequently, the use of fluctuation magnitude, rather than displacement vectors, was incorporated to reduce the risk of signal interference due to partially independent motions of the backbone and sidechain moieties of each residue (*111*). It would prove interesting to combine our approach with more conventional dynamic cross-correlation analyses if we develop a strategy to systematically and reasonably address the misalignment of atomic indexes. Importantly, we demonstrate that a comparison of per-residue RMSF values correlates with immunogenicity (**Figs. S8, S15**), particularly around residues confirmed to be both structurally and biochemically relevant (*22*, *44*, *62*, *71–73*, *104*, *105*).

In terms of dynamical network construction, a range of different methods have been published (*86*, *112*, *113*), including those that use correlation or distance thresholds. To our knowledge, only one other study has explored pMHC interface dynamics through a combination of both MD and network analyses (*86*), however, significant differences exist between our methods. For instance, they utilize more conventional cross-correlation and suboptimal path analyses on a wide range of human HLA-A2 pMHCs to assess long-range allostery between the α3 domain and the peptide-binding cleft. We did not include the α3 domain in the downstream analyses of our MD trajectories, as we could not ensure the equilibration of interdomain motions between the α3 domain or β_2_m chain and the rest of the pMHC (*96*). This provided the additional benefit of avoiding conclusions drawn from regions that could not be readily modeled crystallographically (*62*). We further incorporated correlational significance testing and an additional optimization step with custom validation metrics to ensure our network was biophysically relevant. In the end, the network excelled at uncovering previously hidden dynamic couplings a the pMHC interface, which were not readily accessible through B-factor, ensemble refinement, or conventional MD trajectory analyses.

Furthermore, the presented current workflow could be extended to include the TCR and assess the dynamic coupling of the p3P substitution or more conventional APL mutations at anchor positions with residues in the CDR loops (*94*). If such relationships exist, they may be linked to important biophysical features that govern immunogenicity, such as the propensity for catch bond formation at the TCR interface (*48–52*). Additionally, the role of coupled dynamics in stabilizing pMHC through β_2_m (*93*, *105*) could be investigated using similar methods. Identifying conserved networks of dynamically coupled residues across various pMHCs or pMHC-TCR pairings could significantly enhance our understanding of the co-evolution and germline-encoded affinity between pMHCs and TCRs (*114*). Ultimately, this workflow, coupled with pMHC structure prediction pipelines (*115*), may also be useful in pre-screening strategies and assessing the dynamics of tumor-associated antigens and neoantigens, as well as potential APLs for peptide vaccinations and other immunotherapeutic strategies (*5*).

## METHODS

All methods are described reproducibly here; however, the full implementation is available in the supplement (**Extended Methods**).

### Ensemble refinement

Previously deposited crystal structures for unbound and P14 TCR-bound H-2D^b^/gp33, H-2D^b^/V3P, H-2D^b^/Y4F, and H-2D^b^/PF were subject to crystal contact analysis and ensemble refinement. The latter was performed with readily available tools which perform maximum-likelihood time-averaged restrained molecular dynamics simulations. The generated ensembles and corresponding per-residue RMSF profiles of interfacial residues were used to assess the crystallographic conformational sampling and dynamics of various residues and regions of interest.

### Molecular dynamics

Each complex copy present in each asymmetric unit cell of the unbound pMHC structures was used to generate an initial configuration. Furthermore, *in silico* controls to account for initial configuration bias were prepared by manually mutating p3 to a proline or valine as required. Hydrogen mass repartitioning was applied to these initial models to allow for larger 5 fs timesteps throughout the simulations. Following solvation in a cubic box containing 150 mM NaCl and waters, each system was energy-minimized, then subject to 200 ps NVT and 1 ns NPT equilibration steps. Subsequently, each simulation was run for 200 ns. Each trajectory was analyzed in a similar manner as the crystallographic ensembles by assessing the conformational sampling and RMSF profiles of residues.

### Correlation analysis

The Spearman rank correlation between all per-residue RMSF magnitudes of the pMHC interface was calculated. Significant correlations, as identified through permutation testing at 5% significance, were mapped back onto the model of copy A in the H-2D^b^/gp33 crystal structure. As an alternative, correlational clustering was achieved by applying agglomerative hierarchical clustering on the inter-residue Spearman rank correlation distances.

### Network analysis

The correlational coupling between residues was encoded as a network, by defining each interfacial residue as a node and converting significant correlations between residues to edges. The best way to define these edges mathematically to obtain a biophysically relevant network was unclear, as such the three aforementioned validation metrics were devised: F_1_, F_2_, and F_3_ (**Extended Methods**). Each of these metrics represent the proportion of shortest paths where the node-to-target distance diverges in semantically different manners. A 2D parameter scan of different C_α_ distance thresholds and cost models was used to generate multiple networks which were validated. An optimized and biophysically relevant network was chosen by minimizing F_1_, deemed the most deleterious, and complexity of the edge weight formulation, while maximizing the number of nodes in the largest connected component. This network, with a distance threshold of 10 Å and no associated cost model, was subject to standard network methods, including centrality analysis, shortest path mapping and iterative Louvain clustering.

## Author contributions

AA, BMS, CZ, EA, FSO, HGL, RS, TS, TR, and XH designed research. BMS and TR performed research. AA, BMS, CZ, TS, and TR contributed new analytic tools. BMS and TR analyzed data. AA, BMS, CZ, EA, FSO, HGL, RS, TS, TR, and XH wrote the paper.

## Data and materials availability

All data and code are available from the following Zenodo repository: (persistent link will be provided upon acceptance)

## References and Notes

1. C. Szeto, C. A. Lobos, A. T. Nguyen, S. Gras, TCR Recognition of Peptide-MHC-I: Rule Makers and Breakers. Int J Mol Sci 22, 1–26 (2020).

2. M. Wieczorek, E. T. Abualrous, J. Sticht, M. Álvaro-Benito, S. Stolzenberg, F. Noé, C. Freund, Major Histocompatibility Complex (MHC) Class I and MHC Class II Proteins: Conformational Plasticity in Antigen Presentation. Front Immunol 8, 292 (2017).

3. D. Dersh, J. Hollý, J. W. Yewdell, A few good peptides: MHC class I-based cancer immunosurveillance and immunoevasion. Nat Rev Immunol 21, 116–128 (2021).

4. R. A. Mariuzza, P. Agnihotri, J. Orban, The structural basis of T-cell receptor (TCR) activation: An enduring enigma. J Biol Chem 295, 914–925 (2020).

5. M. Candia, B. Kratzer, W. F. Pickl, On Peptides and Altered Peptide Ligands: From Origin, Mode of Action and Design to Clinical Application (Immunotherapy). Int Arch Allergy Immunol 170, 211–233 (2016).

6. M. Beerbaum, M. Ballaschk, N. Erdmann, C. Schnick, A. Diehl, B. Uchanska-Ziegler, A. Ziegler, P. Schmieder, NMR spectroscopy reveals unexpected structural variation at the protein-protein interface in MHC class I molecules. J Biomol NMR 57, 167–78 (2013).

7. A. R. Smith, J. A. Alonso, C. M. Ayres, N. K. Singh, L. M. Hellman, B. M. Baker, Structurally silent peptide anchor modifications allosterically modulate T cell recognition in a receptor-dependent manner. Proc Natl Acad Sci U S A 118, 2018125118 (2021).

8. J. Sloan-Lancaster, P. M. Allen, Altered peptide ligand-induced partial T cell activation: molecular mechanisms and role in T cell biology. Annu Rev Immunol 14, 1–27 (1996).

9. B. M. Baker, S. J. Gagnon, W. E. Biddison, D. C. Wiley, Conversion of a T cell antagonist into an agonist by repairing a defect in the TCR/peptide/MHC interface: implications for TCR signaling. Immunity 13, 475–84 (2000).

10. B. M. Baker, R. V Turner, S. J. Gagnon, D. C. Wiley, W. E. Biddison, Identification of a crucial energetic footprint on the alpha1 helix of human histocompatibility leukocyte antigen (HLA)-A2 that provides functional interactions for recognition by tax peptide/HLA-A2-specific T cell receptors. J Exp Med 193, 551–62 (2001).

11. T. K. Baxter, S. J. Gagnon, R. L. Davis-Harrison, J. C. Beck, A.-K. Binz, R. V Turner, W. E. Biddison, B. M. Baker, Strategic mutations in the class I major histocompatibility complex HLA-A2 independently affect both peptide binding and T cell receptor recognition. J Biol Chem 279, 29175–84 (2004).

12. S. J. Gagnon, O. Y. Borbulevych, R. L. Davis-Harrison, T. K. Baxter, J. R. Clemens, K. M. Armstrong, R. V Turner, M. Damirjian, W. E. Biddison, B. M. Baker, Unraveling a hotspot for TCR recognition on HLA-A2: evidence against the existence of peptide-independent TCR binding determinants. J Mol Biol 353, 556–73 (2005).

13. P. J. Miller, Y. Pazy, B. Conti, D. Riddle, E. Appella, E. J. Collins, Single MHC mutation eliminates enthalpy associated with T cell receptor binding. J Mol Biol 373, 315–27 (2007).

14. K. L. Rock, E. Reits, J. Neefjes, Present Yourself! By MHC Class I and MHC Class II Molecules. Trends Immunol 37, 724–737 (2016).

15. S. H. van der Burg, M. J. Visseren, R. M. Brandt, W. M. Kast, C. J. Melief, Immunogenicity of peptides bound to MHC class I molecules depends on the MHC-peptide complex stability. J Immunol 156, 3308–14 (1996).

16. D. H. Busch, E. G. Pamer, MHC class I/peptide stability: implications for immunodominance, in vitro proliferation, and diversity of responding CTL. J Immunol 160, 4441–8 (1998).

17. M. Harndahl, M. Rasmussen, G. Roder, I. Dalgaard Pedersen, M. Sørensen, M. Nielsen, S. Buus, Peptide-MHC class I stability is a better predictor than peptide affinity of CTL immunogenicity. Eur J Immunol 42, 1405–16 (2012).

18. M. Rasmussen, E. Fenoy, M. Harndahl, A. B. Kristensen, I. K. Nielsen, M. Nielsen, S. Buus, Pan-Specific Prediction of Peptide-MHC Class I Complex Stability, a Correlate of T Cell Immunogenicity. J Immunol 197, 1517–24 (2016).

19. C. A. Lazarski, F. A. Chaves, S. A. Jenks, S. Wu, K. A. Richards, J. M. Weaver, A. J. Sant, The kinetic stability of MHC class II:peptide complexes is a key parameter that dictates immunodominance. Immunity 23, 29–40 (2005).

20. S. Nicholls, K. P. Piper, F. Mohammed, T. R. Dafforn, S. Tenzer, M. Salim, P. Mahendra, C. Craddock, P. van Endert, H. Schild, M. Cobbold, V. H. Engelhard, P. A. H. Moss, B. E. Willcox, Secondary anchor polymorphism in the HA-1 minor histocompatibility antigen critically affects MHC stability and TCR recognition. Proc Natl Acad Sci U S A 106, 3889–94 (2009).

21. E. Spierings, S. Gras, J.-B. Reiser, B. Mommaas, M. Almekinders, M. G. D. Kester, A. Chouquet, M. Le Gorrec, J. W. Drijfhout, F. Ossendorp, D. Housset, E. Goulmy, Steric hindrance and fast dissociation explain the lack of immunogenicity of the minor histocompatibility HA-1Arg Null allele. J Immunol 182, 4809–16 (2009).

22. C. Madhurantakam, A. D. Duru, T. Sandalova, J. R. Webb, A. Achour, Inflammation-associated nitrotyrosination affects TCR recognition through reduced stability and alteration of the molecular surface of the MHC complex. PLoS One 7, e32805 (2012).

23. L. L. Jones, L. A. Colf, A. J. Bankovich, J. D. Stone, Y.-G. Gao, C. M. Chan, R. H. Huang, K. C. Garcia, D. M. Kranz, Different thermodynamic binding mechanisms and peptide fine specificities associated with a panel of structurally similar high-affinity T cell receptors. Biochemistry 47, 12398–408 (2008).

24. N. L. La Gruta, S. Gras, S. R. Daley, P. G. Thomas, J. Rossjohn, Understanding the drivers of MHC restriction of T cell receptors. Nat Rev Immunol 18, 467–478 (2018).

25. M. G. Rudolph, R. L. Stanfield, I. A. Wilson, How TCRs bind MHCs, peptides, and coreceptors. Annu Rev Immunol 24, 419–66 (2006).

26. J. J. Adams, S. Narayanan, B. Liu, M. E. Birnbaum, A. C. Kruse, N. A. Bowerman, W. Chen, A. M. Levin, J. M. Connolly, C. Zhu, D. M. Kranz, K. C. Garcia, T cell receptor signaling is limited by docking geometry to peptide-major histocompatibility complex. Immunity 35, 681–93 (2011).

27. M. Aleksic, O. Dushek, H. Zhang, E. Shenderov, J.-L. Chen, V. Cerundolo, D. Coombs, P. A. van der Merwe, Dependence of T cell antigen recognition on T cell receptor-peptide MHC confinement time. Immunity 32, 163–74 (2010).

28. C. C. Govern, M. K. Paczosa, A. K. Chakraborty, E. S. Huseby, Fast on-rates allow short dwell time ligands to activate T cells. Proc Natl Acad Sci U S A 107, 8724–9 (2010).

29. J. Huang, V. I. Zarnitsyna, B. Liu, L. J. Edwards, N. Jiang, B. D. Evavold, C. Zhu, The kinetics of two-dimensional TCR and pMHC interactions determine T-cell responsiveness. Nature 464, 932–6 (2010).

30. J. B. Huppa, M. Axmann, M. A. Mörtelmaier, B. F. Lillemeier, E. W. Newell, M. Brameshuber, L. O. Klein, G. J. Schütz, M. M. Davis, TCR-peptide-MHC interactions in situ show accelerated kinetics and increased affinity. Nature 463, 963–7 (2010).

31. C. Klammt, L. Novotná, D. T. Li, M. Wolf, A. Blount, K. Zhang, J. R. Fitchett, B. F. Lillemeier, T cell receptor dwell times control the kinase activity of Zap70. Nat Immunol 16, 961–9 (2015).

32. A. M. Kalergis, N. Boucheron, M. A. Doucey, E. Palmieri, E. C. Goyarts, Z. Vegh, I. F. Luescher, S. G. Nathenson, Efficient T cell activation requires an optimal dwell-time of interaction between the TCR and the pMHC complex. Nat Immunol 2, 229–34 (2001).

33. J. J. Boniface, Z. Reich, D. S. Lyons, M. M. Davis, Thermodynamics of T cell receptor binding to peptide-MHC: evidence for a general mechanism of molecular scanning. Proc Natl Acad Sci U S A 96, 11446–51 (1999).

34. B. E. Willcox, G. F. Gao, J. R. Wyer, J. E. Ladbury, J. I. Bell, B. K. Jakobsen, P. A. van der Merwe, TCR binding to peptide-MHC stabilizes a flexible recognition interface. Immunity 10, 357–65 (1999).

35. M. Krogsgaard, N. Prado, E. J. Adams, X. He, D.-C. Chow, D. B. Wilson, K. C. Garcia, M. M. Davis, Evidence that structural rearrangements and/or flexibility during TCR binding can contribute to T cell activation. Mol Cell 12, 1367–78 (2003).

36. J. K. Lee, G. Stewart-Jones, T. Dong, K. Harlos, K. Di Gleria, L. Dorrell, D. C. Douek, P. A. van der Merwe, E. Y. Jones, A. J. McMichael, T cell cross-reactivity and conformational changes during TCR engagement. J Exp Med 200, 1455–66 (2004).

37. R. L. Davis-Harrison, K. M. Armstrong, B. M. Baker, Two different T cell receptors use different thermodynamic strategies to recognize the same peptide/MHC ligand. J Mol Biol 346, 533–50 (2005).

38. L. K. Ely, T. Beddoe, C. S. Clements, J. M. Matthews, A. W. Purcell, L. Kjer-Nielsen, J. McCluskey, J. Rossjohn, Disparate thermodynamics governing T cell receptor-MHC-I interactions implicate extrinsic factors in guiding MHC restriction. Proc Natl Acad Sci U S A 103, 6641–6 (2006).

39. L. A. Colf, A. J. Bankovich, N. A. Hanick, N. A. Bowerman, L. L. Jones, D. M. Kranz, K. C. Garcia, How a single T cell receptor recognizes both self and foreign MHC. Cell 129, 135–46 (2007).

40. C. Mazza, N. Auphan-Anezin, C. Gregoire, A. Guimezanes, C. Kellenberger, A. Roussel, A. Kearney, P. A. van der Merwe, A.-M. Schmitt-Verhulst, B. Malissen, How much can a T-cell antigen receptor adapt to structurally distinct antigenic peptides? EMBO J 26, 1972–83 (2007).

41. K. M. Armstrong, F. K. Insaidoo, B. M. Baker, Thermodynamics of T-cell receptor-peptide/MHC interactions: progress and opportunities. J Mol Recognit 21, 275–87 (2008).

42. K. M. Armstrong, K. H. Piepenbrink, B. M. Baker, Conformational changes and flexibility in T-cell receptor recognition of peptide-MHC complexes. Biochem J 415, 183– 96 (2008).

43. S. P. Persaud, D. L. Donermeyer, K. S. Weber, D. M. Kranz, P. M. Allen, High-affinity T cell receptor differentiates cognate peptide-MHC and altered peptide ligands with distinct kinetics and thermodynamics. Mol Immunol 47, 1793–801 (2010).

44. E. B. Allerbring, A. D. Duru, H. Uchtenhagen, C. Madhurantakam, M. B. Tomek, S. Grimm, P. A. Mazumdar, R. Friemann, M. Uhlin, T. Sandalova, P.-Å. Nygren, A. Achour, Unexpected T-cell recognition of an altered peptide ligand is driven by reversed thermodynamics. Eur J Immunol 42, 2990–3000 (2012).

45. S. E. Kerry, J. Buslepp, L. A. Cramer, R. Maile, L. L. Hensley, A. I. Nielsen, P. Kavathas, B. J. Vilen, E. J. Collins, J. A. Frelinger, Interplay between TCR affinity and necessity of coreceptor ligation: high-affinity peptide-MHC/TCR interaction overcomes lack of CD8 engagement. J Immunol 171, 4493–503 (2003).

46. P. D. Holler, D. M. Kranz, Quantitative analysis of the contribution of TCR/pepMHC affinity and CD8 to T cell activation. Immunity 18, 255–64 (2003).

47. S. Gras, J. Chadderton, C. M. Del Campo, C. Farenc, F. Wiede, T. M. Josephs, X. Y. X. Sng, M. Mirams, K. A. Watson, T. Tiganis, K. M. Quinn, J. Rossjohn, N. L. La Gruta, Reversed T Cell Receptor Docking on a Major Histocompatibility Class I Complex Limits Involvement in the Immune Response. Immunity 45, 749–760 (2016).

48. S. T. Kim, K. Takeuchi, Z.-Y. J. Sun, M. Touma, C. E. Castro, A. Fahmy, M. J. Lang, G. Wagner, E. L. Reinherz, The alphabeta T cell receptor is an anisotropic mechanosensor. J Biol Chem 284, 31028–37 (2009).

49. S. T. Kim, Y. Shin, K. Brazin, R. J. Mallis, Z.-Y. J. Sun, G. Wagner, M. J. Lang, E. L. Reinherz, TCR Mechanobiology: Torques and Tunable Structures Linked to Early T Cell Signaling. Front Immunol 3, 76 (2012).

50. B. Liu, W. Chen, B. D. Evavold, C. Zhu, Accumulation of dynamic catch bonds between TCR and agonist peptide-MHC triggers T cell signaling. Cell 157, 357–368 (2014).

51. J. Hong, S. P. Persaud, S. Horvath, P. M. Allen, B. D. Evavold, C. Zhu, Force-Regulated In Situ TCR-Peptide-Bound MHC Class II Kinetics Determine Functions of CD4+ T Cells. J Immunol 195, 3557–64 (2015).

52. Y. Liu, L. Blanchfield, V. P.-Y. Ma, R. Andargachew, K. Galior, Z. Liu, B. Evavold, K. Salaita, DNA-based nanoparticle tension sensors reveal that T-cell receptors transmit defined pN forces to their antigens for enhanced fidelity. Proc Natl Acad Sci U S A 113, 5610–5 (2016).

53. D. H. Busch, E. G. Pamer, T cell affinity maturation by selective expansion during infection. J Exp Med 189, 701–10 (1999).

54. L. Malherbe, C. Hausl, L. Teyton, M. G. McHeyzer-Williams, Clonal selection of helper T cells is determined by an affinity threshold with no further skewing of TCR binding properties. Immunity 21, 669–79 (2004).

55. R. H. McMahan, J. A. McWilliams, K. R. Jordan, S. W. Dow, D. B. Wilson, J. E. Slansky, Relating TCR-peptide-MHC affinity to immunogenicity for the design of tumor vaccines. J Clin Invest 116, 2543–51 (2006).

56. J. M. Boulter, N. Schmitz, A. K. Sewell, A. J. Godkin, M. F. Bachmann, A. M. Gallimore, Potent T cell agonism mediated by a very rapid TCR/pMHC interaction. Eur J Immunol 37, 798–806 (2007).

57. S. Tian, R. Maile, E. J. Collins, J. A. Frelinger, CD8+ T cell activation is governed by TCR-peptide/MHC affinity, not dissociation rate. J Immunol 179, 2952–60 (2007).

58. D. Zehn, S. Y. Lee, M. J. Bevan, Complete but curtailed T-cell response to very low-affinity antigen. Nature 458, 211–4 (2009).

59. A. S. Chervin, J. D. Stone, P. D. Holler, A. Bai, J. Chen, H. N. Eisen, D. M. Kranz, The impact of TCR-binding properties and antigen presentation format on T cell responsiveness. J Immunol 183, 1166–78 (2009).

60. D. A. Schmid, M. B. Irving, V. Posevitz, M. Hebeisen, A. Posevitz-Fejfar, J.-C. F. Sarria, R. Gomez-Eerland, M. Thome, T. N. M. Schumacher, P. Romero, D. E. Speiser, V. Zoete, O. Michielin, N. Rufer, Evidence for a TCR affinity threshold delimiting maximal CD8 T cell function. J Immunol 184, 4936–46 (2010).

61. J. Gálvez, J. J. Gálvez, P. García-Peñarrubia, Is TCR/pMHC Affinity a Good Estimate of the T-cell Response? An Answer Based on Predictions From 12 Phenotypic Models. Front Immunol 10, 349 (2019).

62. A. D. Duru, R. Sun, E. B. Allerbring, J. Chadderton, N. Kadri, X. Han, K. Peqini, H. Uchtenhagen, C. Madhurantakam, S. Pellegrino, T. Sandalova, P.-Å. Nygren, S. J. Turner, A. Achour, Tuning antiviral CD8 T-cell response via proline-altered peptide ligand vaccination. PLoS Pathog 16, e1008244 (2020).

63. L. V Sibener, R. A. Fernandes, E. M. Kolawole, C. B. Carbone, F. Liu, D. McAffee, M. E. Birnbaum, X. Yang, L. F. Su, W. Yu, S. Dong, M. H. Gee, K. M. Jude, M. M. Davis, J. T. Groves, W. A. Goddard, J. R. Heath, B. D. Evavold, R. D. Vale, K. C. Garcia, Isolation of a Structural Mechanism for Uncoupling T Cell Receptor Signaling from Peptide-MHC Binding. Cell 174, 672–687.e27 (2018).

64. L. S. Klavinskis, J. L. Whitton, E. Joly, M. B. Oldstone, Vaccination and protection from a lethal viral infection: identification, incorporation, and use of a cytotoxic T lymphocyte glycoprotein epitope. Virology 178, 393–400 (1990).

65. M. B. Oldstone, J. L. Whitton, H. Lewicki, A. Tishon, Fine dissection of a nine amino acid glycoprotein epitope, a major determinant recognized by lymphocytic choriomeningitis virus-specific class I-restricted H-2Db cytotoxic T lymphocytes. J Exp Med 168, 559–70 (1988).

66. J. E. Gairin, M. B. Oldstone, Design of high-affinity major histocompatibility complex-specific antagonist peptides that inhibit cytotoxic T-lymphocyte activity: implications for control of viral disease. J Virol 66, 6755–62 (1992).

67. H. Pircher, D. Moskophidis, U. Rohrer, K. Bürki, H. Hengartner, R. M. Zinkernagel, Viral escape by selection of cytotoxic T cell-resistant virus variants in vivo. Nature 346, 629–33 (1990).

68. D. Moskophidis, R. M. Zinkernagel, Immunobiology of cytotoxic T-cell escape mutants of lymphocytic choriomeningitis virus. J Virol 69, 2187–93 (1995).

69. D. Hudrisier, M. B. Oldstone, J. E. Gairin, The signal sequence of lymphocytic choriomeningitis virus contains an immunodominant cytotoxic T cell epitope that is restricted by both H-2D(b) and H-2K(b) molecules. Virology 234, 62–73 (1997).

70. A. C. Tissot, C. Ciatto, P. R. Mittl, M. G. Grütter, A. Plückthun, Viral escape at the molecular level explained by quantitative T-cell receptor/peptide/MHC interactions and the crystal structure of a peptide/MHC complex. J Mol Biol 302, 873–85 (2000).

71. L. M. Velloso, J. Michaëlsson, H.-G. Ljunggren, G. Schneider, A. Achour, Determination of structural principles underlying three different modes of lymphocytic choriomeningitis virus escape from CTL recognition. J Immunol 172, 5504–11 (2004).

72. T. Sandalova, J. Michaëlsson, R. A. Harris, J. Odeberg, G. Schneider, K. Kärre, A. Achour, A structural basis for CD8+ T cell-dependent recognition of non-homologous peptide ligands: implications for molecular mimicry in autoreactivity. J Biol Chem 280, 27069–75 (2005).

73. A. Achour, J. Michaëlsson, R. A. Harris, J. Odeberg, P. Grufman, J. K. Sandberg, V. Levitsky, K. Kärre, T. Sandalova, G. Schneider, A structural basis for LCMV immune evasion: subversion of H-2D(b) and H-2K(b) presentation of gp33 revealed by comparative crystal structure analyses. Immunity 17, 757–68 (2002).

74. T. Aebischer, D. Moskophidis, U. H. Rohrer, R. M. Zinkernagel, H. Hengartner, In vitro selection of lymphocytic choriomeningitis virus escape mutants by cytotoxic T lymphocytes. Proc Natl Acad Sci U S A 88, 11047–51 (1991).

75. M. T. Puglielli, A. J. Zajac, R. G. van der Most, J. L. Dzuris, A. Sette, J. D. Altman, R. Ahmed, In vivo selection of a lymphocytic choriomeningitis virus variant that affects recognition of the GP33-43 epitope by H-2Db but not H-2Kb. J Virol 75, 5099–107 (2001).

76. H. Uchtenhagen, E. T. Abualrous, E. Stahl, E. B. Allerbring, M. Sluijter, M. Zacharias, T. Sandalova, T. van Hall, S. Springer, P.-Å. Nygren, A. Achour, Proline substitution independently enhances H-2D(b) complex stabilization and TCR recognition of melanoma-associated peptides. Eur J Immunol 43, 3051–60 (2013).

77. S. Watts, C. Wheeler, R. Morse, R. S. Goodenow, Amino acid comparison of the class I antigens of mouse major histocompatibility complex. Immunogenetics 30, 390–2 (1989).

78. D. J. Barker, G. Maccari, X. Georgiou, M. A. Cooper, P. Flicek, J. Robinson, S. G. E. Marsh, The IPD-IMGT/HLA Database. Nucleic Acids Res 51, D1053–D1060 (2023).

79. M. J. B. van Stipdonk, D. Badia-Martinez, M. Sluijter, R. Offringa, T. van Hall, A. Achour, Design of agonistic altered peptides for the robust induction of CTL directed towards H-2Db in complex with the melanoma-associated epitope gp100. Cancer Res 69, 7784–92 (2009).

80. I. Hafstrand, E. M. Doorduijn, A. D. Duru, J. Buratto, C. C. Oliveira, T. Sandalova, T. van Hall, A. Achour, The MHC Class I Cancer-Associated Neoepitope Trh4 Linked with Impaired Peptide Processing Induces a Unique Noncanonical TCR Conformer. The Journal of Immunology 196, 2327–2334 (2016).

81. E. M. Doorduijn, M. Sluijter, B. J. Querido, C. C. Oliveira, A. Achour, F. Ossendorp, S. H. van der Burg, T. van Hall, TAP-independent self-peptides enhance T cell recognition of immune-escaped tumors. J Clin Invest 126, 784–94 (2016).

82. I. Hafstrand, E. M. Doorduijn, R. Sun, A. Talyzina, M. Sluijter, S. Pellegrino, T. Sandalova, A. D. Duru, T. van Hall, A. Achour, The Immunogenicity of a Proline-Substituted Altered Peptide Ligand toward the Cancer-Associated TEIPP Neoepitope Trh4 Is Unrelated to Complex Stability. The Journal of Immunology 200, 2860–2868 (2018).

83. I. Kass, A. M. Buckle, N. A. Borg, Understanding the structural dynamics of TCR-pMHC complex interactions. Trends Immunol 35, 604–612 (2014).

84. J. Fodor, B. T. Riley, N. A. Borg, A. M. Buckle, Previously Hidden Dynamics at the TCR–Peptide–MHC Interface Revealed. The Journal of Immunology 200, 4134–4145 (2018).

85. K. Natarajan, J. Jiang, N. A. May, M. G. Mage, L. F. Boyd, A. C. McShan, N. G. Sgourakis, A. Bax, D. H. Margulies, The Role of Molecular Flexibility in Antigen Presentation and T Cell Receptor-Mediated Signaling. Front Immunol 9, 1657 (2018).

86. C. M. Ayres, E. T. Abualrous, A. Bailey, C. Abraham, L. M. Hellman, S. A. Corcelli, F. Noé, T. Elliott, B. M. Baker, Dynamically driven allostery in MHC proteins: Peptide-dependent tuning of class I MHC global flexibility. Front Immunol 10 (2019).

87. C. Szeto, J. I. Bloom, H. Sloane, C. A. Lobos, J. Fodor, D. Jayasinghe, D. S. M. Chatzileontiadou, E. J. Grant, A. M. Buckle, S. Gras, Impact of HLA-DR antigen binding cleft rigidity on T cell recognition. Int J Mol Sci 21, 1–20 (2020).

88. A. M. Buckle, N. A. Borg, Integrating Experiment and Theory to Understand TCR-pMHC Dynamics. Front Immunol 9, 2898 (2018).

89. F. K. Insaidoo, J. Zajicek, B. M. Baker, A general and efficient approach for NMR studies of peptide dynamics in class I MHC peptide binding grooves. Biochemistry 48, 9708– 9710 (2009).

90. T. Pöhlmann, R. A. Böckmann, H. Grubmüller, B. Uchanska-Ziegler, A. Ziegler, U. Alexiev, Differential peptide dynamics is linked to major histocompatibility complex polymorphism. Journal of Biological Chemistry 279, 28197–28201 (2004).

91. F. K. Insaidoo, O. Y. Borbulevych, M. Hossain, S. M. Santhanagopolan, T. K. Baxter, B. M. Baker, Loss of T cell antigen recognition arising from changes in peptide and major histocompatibility complex protein flexibility: Implications for vaccine design. Journal of Biological Chemistry 286, 40163–40173 (2011).

92. O. Serçinoğlu, P. Ozbek, Computational characterization of residue couplings and micropolymorphism-induced changes in the dynamics of two differentially disease-associated human MHC class-I alleles. J Biomol Struct Dyn 36, 724–740 (2018).

93. A. K. Binz, R. C. Rodriguez, W. E. Biddison, B. M. Baker, Thermodynamic and kinetic analysis of a peptide-class I MHC interaction highlights the noncovalent nature and conformational dynamics of the class I heterotrimer. Biochemistry 42, 4954–4961 (2003).

94. 94. D. M. Gakamsky, E. Lewitzki, E. Grell, X. Saulquin, B. Malissen, F. Montero-Julian, M. Bonneville ¶ † †, I. Pecht, “Kinetic evidence for a ligand-binding-induced conformational transition in the T cell receptor” (2007).

95. B. T. Burnley, P. V Afonine, P. D. Adams, P. Gros, Modelling dynamics in protein crystal structures by ensemble refinement. Elife 1, e00311 (2012).

96. L. Tomasiak, R. Karch, W. Schreiner, Conformational flexibility of a free and TCR-bound pMHC-I protein investigated by long-term molecular dynamics simulations. BMC Immunol 23, 36 (2022).

97. W. F. Hawse, M. M. Champion, M. V Joyce, L. M. Hellman, M. Hossain, V. Ryan, B. G. Pierce, Z. Weng, B. M. Baker, Cutting edge: Evidence for a dynamically driven T cell signaling mechanism. J Immunol 188, 5819–23 (2012).

98. H. G. Rammensee, T. Friede, S. Stevanoviíc, MHC ligands and peptide motifs: first listing. Immunogenetics 41, 178–228 (1995).

99. R. L. Dunbrack, M. Karplus, Conformational analysis of the backbone-dependent rotamer preferences of protein sidechains. Nat Struct Biol 1, 334–40 (1994).

100. S. Yanaka, T. Ueno, Y. Shi, J. Qi, G. F. Gao, K. Tsumoto, K. Sugase, Peptide-dependent conformational fluctuation determines the stability of the human leukocyte antigen class I complex. J Biol Chem 289, 24680–90 (2014).

101. D. Narzi, C. M. Becker, M. T. Fiorillo, B. Uchanska-Ziegler, A. Ziegler, R. A. Böckmann, Dynamical characterization of two differentially disease associated MHC class I proteins in complex with viral and self-peptides. J Mol Biol 415, 429–42 (2012).

102. M. Zacharias, S. Springer, Conformational flexibility of the MHC class I alpha1-alpha2 domain in peptide bound and free states: a molecular dynamics simulation study. Biophys J 87, 2203–14 (2004).

103. B. M. Baker, D. R. Scott, S. J. Blevins, W. F. Hawse, Structural and dynamic control of T-cell receptor specificity, cross-reactivity, and binding mechanism. Immunol Rev 250, 10– 31 (2012).

104. T. Sandalova, J. Michaëlsson, R. A. Harris, H.-G. Ljunggren, K. Kärre, G. Schneider, A. Achour, Expression, refolding and crystallization of murine MHC class I H-2Db in complex with human beta2-microglobulin. Acta Crystallogr Sect F Struct Biol Cryst Commun 61, 1090–3 (2005).

105. A. Achour, J. Michaëlsson, R. A. Harris, H. G. Ljunggren, K. Kärre, G. Schneider, T. Sandalova, Structural basis of the differential stability and receptor specificity of H-2Db in complex with murine versus human β2-microglobulin. J Mol Biol 356, 382–396 (2006).

106. F. Ballabio, L. Broggini, C. Paissoni, X. Han, K. Peqini, B. M. Sala, R. Sun, T. Sandalova, A. Barbiroli, A. Achour, S. Pellegrino, S. Ricagno, C. Camilloni, l- to d-Amino Acid Substitution in the Immunodominant LCMV-Derived Epitope gp33 Highlights the Sensitivity of the TCR Recognition Mechanism for the MHC/Peptide Structure and Dynamics. ACS Omega 7, 9622–9635 (2022).

107. T. Ichiye, M. Karplus, Collective motions in proteins: a covariance analysis of atomic fluctuations in molecular dynamics and normal mode simulations. Proteins 11, 205–17 (1991).

108. R. A. Estabrook, J. Luo, M. M. Purdy, V. Sharma, P. Weakliem, T. C. Bruice, N. O. Reich, Statistical coevolution analysis and molecular dynamics: identification of amino acid pairs essential for catalysis. Proc Natl Acad Sci U S A 102, 994–9 (2005).

109. H. Kamberaj, A. van der Vaart, Correlated motions and interactions at the onset of the DNA-induced partial unfolding of Ets-1. Biophys J 96, 1307–17 (2009).

110. K. Kasahara, I. Fukuda, H. Nakamura, A novel approach of dynamic cross correlation analysis on molecular dynamics simulations and its application to Ets1 dimer-DNA complex. PLoS One 9, e112419 (2014).

111. J. E. Fuchs, B. J. Waldner, R. G. Huber, S. von Grafenstein, C. Kramer, K. R. Liedl, Independent Metrics for Protein Backbone and Side-Chain Flexibility: Time Scales and Effects of Ligand Binding. J Chem Theory Comput 11, 851–60 (2015).

112. W. Yan, J. Zhou, M. Sun, J. Chen, G. Hu, B. Shen, The construction of an amino acid network for understanding protein structure and function. Amino Acids 46, 1419–39 (2014).

113. J. Salamanca Viloria, M. F. Allega, M. Lambrughi, E. Papaleo, An optimal distance cutoff for contact-based Protein Structure Networks using side-chain centers of mass. Sci Rep 7, 2838 (2017).

114. E. J. Steele, R. A. Lindley, Regulatory T cells and co-evolution of allele-specific MHC recognition by the TCR. Scand J Immunol 91, e12853 (2020).

115. V. Mikhaylov, C. A. Brambley, G. L. J. Keller, A. G. Arbuiso, L. I. Weiss, B. M. Baker, A. J. Levine, Accurate modeling of peptide-MHC structures with AlphaFold. Structure 32, 228–241.e4 (2024).

