## Supplementary Information for "From Structure to Immunogenicity: Decoding Correlated Dynamics at the Peptide MHC interface to Understand TCR Recognition"

### EXTENDED METHODS

#### Atomic coordinates and ensemble refinement of complexes

Fully refined models for all pMHC and TCR/pMHC complexes were obtained from the following Protein Data Bank (PDB) entries: 1S7U, 4NSK, 1S7X, 3TBY, 5TJE, 5TIL, and 5M02 (*62*, *71*, *73*). In *Coot* (*116*), alternative conformations with lower occupancy were removed, and the model was re-refined using *phenix.refine* (*117*, *118*) from the PHENIX 1.19.1-4122 program suit (*119*). The models and phases from the refinement were used as input for the ensemble refinement. The ensemble refinement was performed according to previously published work (*95*, *120*, *121*). *phenix.ensemble_refinement* generates the ensembles via a maximum-likelihood time-averaged restrained molecular dynamics simulation; default parameters were used except for the pTLS parameters, which were optimized for each refinement with values of 1.0, 0.9, 0.8, and 0.7.

#### Structural and ensemble analysis and graphics

The per-residue root-mean-square fluctuations (RMSF) of every ensemble model were obtained using the *ens_tools.py* PyMOL script included with *phenix.ensemble_refinement* (*120*). Structural analyses and representations were performed using PyMOL 2.4.2 (Schrödinger).

#### MD simulations of the H-2D$^b$/peptide complexes

All simulations were performed based on previously published protocols (*106*, *122*) using GROMACS 2021.4 (*123*). The initial configurations were obtained from each individual pMHC molecule in the asymmetric unit cell of the following PDB entries: 1S7U, 4NSK, 1S7X, and 3TBY. Gaps and missing sidechains were modeled in *Coot* with the help of the deposited electron density maps, and, if required, the first molecule in the H-2D$^b$/gp33 crystal structure as reference. Missing termini were not modeled. To account for the bias introduced by our choice of initial configuration, p3 was mutated manually in *Coot* to valine or proline as required, to generate the initial configurations for another set of MD simulations. Real-space refinement in *Coot* was used to remove model distortions in p3 and the immediately preceding and succeeding residues. Using the Amber99SB-ILDN forcefield (*124*) and SPC/E water model (*125*), the topology was generated through the *gmx pdb2gmx* command passed with the *-heavyh* flag. The latter allows for the use of 5 fs timesteps in MD runs through the hydrogen mass repartitioning scheme (*126*) when constraining all bonds with the LINCS algorithm (*127*). Each system was solvated in 150 mM NaCl in a cubic box with periodic boundary conditions and a minimum solute-to-box distance of 1.1 nm. Charges were neutralized by adding $Na^+$ and $Cl^-$ counterions. Following steepest-descent energy-minimization, each system was equilibrated, first a 200 ps NVT simulation with protein position restraints, then a 1 ns NPT simulation. Boltzmann-distributed velocities were assigned at the beginning of the NVT equilibration. Short-range van der Waals and Coulomb interactions were implemented with a 1.0 cutoff, using potential shifting to minimize cutoff truncation artifacts. Long-range electrostatics were handled with the Particle-Mesh Ewald method (*128*). Long-range dispersion corrections for energy and pressure were also applied. Two thermostats, one for the solvent and one for the solute, were implemented through the velocity scaling algorithm (*129*). For the NPT equilibration, pressure was controlled with a Parrinello-Rahman barostat (*130*, *131*). After equilibration, the system was simulated for 200 ns using the same parameters as the NPT equilibration with system coordinates being saved every 100 ps. Each trajectory was gathered and centered on the H-2D$^b$/peptide complex and each frame was roto-translationally fitted to the backbone of the first frame. All simulations were validated by monitoring the root-mean-square deviations compared to the input structure, radius of gyration, and visual inspection of each trajectory for large, unexplained divergences. Additionally, the temperature, pressure, density, and energies were confirmed to be constant.

**Code environments for dynamic analysis of MD trajectories**

R and Python 3 were used to perform analyses on the MD data. The R analyses were run in an R 4.4.1 (*132*) environment with the readxl 1.4.3 (*133*), foreach 1.5.2 (*134*), doParallel 1.0.17 (*135*), tidyverse 2.0.0 (*136*), and ggplot2 3.5.1 (*137*) packages installed. All Python 3 analyses were run using JupyterLab 4.0.6 notebooks (*138*) in a Conda 24.9.1 environment (Anaconda) with Python 3.10.12 (Python Software Foundation) and the NumPy 1.26.0 (*139*), SciPy 1.13.1 (*140*), pandas 2.1.1 (*141*), Matplotlib 3.8.0 (*142*), scikit-learn 1.5.1 (*143*), NetworkX 3.3 (*144*), Biopython 1.81 (*145*), MDTraj 1.10.0 (*146*), and Multiprocess 0.70.15 (*147*) libraries installed.

**Trajectory analysis and graphics**

The per-residue RMSF values were obtained from each trajectory using the energy-minimized structure as reference through the *gmx rmsf* command with the *-res* flag. This RMSF data was subsequently analyzed and plotted using custom R scripts. To visually compare the dynamics across different trajectories, the *gmx rmsf* command was used again to convert the atomic RMSF to anisotropic B-factors, which were written to the B-factor column of a PDB file with the energy-minimized coordinates. This was visualized in PyMOL 3.0.3 (Schrödinger) using stick and ellipsoid representations. The backbone and dihedral angle distributions of each peptide residue throughout all trajectories were obtained using MDTraj.

**Correlational analysis of the RMSF data obtained from MD simulations**

The Spearman rank correlation coefficient ($\rho_s$) of the per-residue RMSF data was calculated for all pairwise combinations of residues to assess their monotonic relationship.

Subsequently, the coefficients were converted to the following test statistic (*t*), where *n* denotes the number of simulations:

$$t = \rho_s \sqrt{\frac{n-2}{1-{\rho_s}^2}}$$

To assess the significance of the observed correlation coefficients, the corresponding p-values were obtained through a two-tailed comparison of the test statistics against the null distribution. This null distribution was generated from test statistics calculated from the correlation of 5 x $10^7$ pseudo-randomly paired resamples of two sequential integer arrays from 1 to 26 inclusive. Both the two-tailed comparison and creation of the null distribution were achieved using refactored code from the *scipy.stats.permutation_test* function and examples described in the SciPy documentation (*140*, *148*). Multiple testing corrections were applied by adjusting the p-values using the Benjamini-Hochberg procedure to control the estimated false discovery rate at 5% significance (*149*).

As an alternative approach, hierarchical agglomerative clustering was applied to the per-residue RMSF data. The clustering was achieved with the UPGMA algorithm (*150*) using the pairwise correlation distance of the ranked RMSF data, equivalent to $1 - \rho_s$, as the distance metric. The threshold for clustering leaves on the dendrogram was determined by identifying a cophenetic distance threshold which included p3V/P in one of the clusters.

**Correlational analysis of per-residue RMSF data and immunogenicity**

The per-residue RMSF values were grouped according to immunogenicity, where p3P trajectories were considered to be more immunogenic than p3V trajectories. This aligns with the published biophysical and functional data (*62*). For each residue of interest, the rank biserial correlation of RMSF values with immunogenicity was calculated. Subsequently, residues with the largest magnitudes of correlation were further subjected to permutation testing to determine the probability of observing the same result through random chance. The permutation test was performed using the same test statistic as before, however, due to the small number of comparisons, no generalized null distribution was generated. Instead, a null distribution was generated from 1 x $10^6$ pseudo-random resamples of the paired RMSF and immunogenicity data for each residue tested.

**Dynamic correlation network construction and validation**

Undirected networks with weighted edges were constructed from the correlation matrix of per-residue RMSF data. In these networks (*G*), all interface residues were assigned as nodes (*V*) and edges and their respective weights (*E*) were derived from the correlation matrix:

$$G = (V, E)$$

A plausible general edge weight metric was defined to later optimize the construction of a biophysically relevant network:

$$E[u,v,n,t] = \begin{cases} \dfrac{|\rho_S[u,v]|}{(r_{C\alpha}[u,v])^n}, r_{C\alpha}[u,v] < t \wedge p_{adj}[u,v] < 0.05 \\ \emptyset, \text{otherwise} \end{cases}$$

*u* and *v* are any two nodes in the network ($u,\ v \in V$), $\rho_s$ is their Spearman rank correlation, $r_{C\alpha}$ is their through-space $C_\alpha$ distance derived from the crystal structure of H-2D$^b$/gp33, and $p_{adj}$ is their Benjamini-Hochberg corrected p-value. Both *n* and *t* are parameters to be optimized and represent the inverse power of distance and distance threshold respectively.

A 2D scan of possible *n* and *t* values was performed to generate different networks. Custom shortest-path-based metrics were defined to help validate each network and assess their biophysical relevance to probing network dynamics. All shortest paths (*P*) for the network *G* were calculated using Dijkstra's algorithm (*151*):

$$\forall P \in G, P = (v_{\text{start}}, v_1, \dots, v_n, v_{\text{end}})$$

These shortest paths were used to calculate the proportion of shortest paths with any through-space edge distance greater than the initial distance ($F_1$), the proportion of shortest paths where the through-space distance increases relative to the preceding nodes ($F_2$), and the proportion of shortest paths where the through-space distance increases relative to the initial distance ($F_3$):

$$F_1[G] = \frac{\sum_{P \in G} f_1[P]}{\sum_{P \in G}}, \text{where } f_1[P] = \begin{cases} 1, r_{C\alpha}[v_n, v_{n+1}] > r_{C\alpha}[v_{\text{start}}, v_{\text{end}}] \\ 0, \text{otherwise} \end{cases}$$

$$F_2[G] = \frac{\sum_{P \in G} f_2[P]}{\sum_{P \in G}}, \text{where } f_2[P] = \begin{cases} 1, r_{C\alpha}[v_n, v_{n-1}] < r_{C\alpha}[v_n, v_{n+1}] \\ 0, \text{otherwise} \end{cases}$$

$$F_3[G] = \frac{\sum_{P \in G} f_3[P]}{\sum_{P \in G}}, \text{where } f_3[P] = \begin{cases} 1, r_{C\alpha}[v_n, v_{\text{end}}] > r_{C\alpha}[v_{\text{start}}, v_{\text{end}}] \\ 0, \text{otherwise} \end{cases}$$

In the end, a biophysically relevant network was chosen based on a compromise between minimizing $F_1$ and edge weight complexity, while maximizing the number of nodes present in the largest connected component (> 90%). $F_2$ and $F_3$ were not directly used in network optimization but are, nonetheless, informative for understanding network properties.

**Network analysis**

Our chosen biophysically relevant ($n = 0$, $t = 10$ Å) and significant correlation network ($n = 0$, $t = 52$ Å) were subject to a suit of standard analyses to investigate network features. Note that the largest observed pairwise $C_\alpha$ distance was 51.1 Å. The topological properties of the networks, such as node centrality, were assessed using built-in functions from NetworkX and mapping these values back onto the crystallographic structure of H-2D$^b$/gp33. For all centrality measures, except for degree centrality and coreness, edge weight (*E*) or edge distance ($E^{-1}$) were incorporated as required. The shortest paths originating from p3V/P, as identified previously using Dijkstra's algorithm, were also mapped to the structure. Lastly, the community structure of dynamics at the pMHC interface was also assessed using the Louvain community detection algorithm (*152*). Due to the heuristic nature of this approach, the Louvain algorithm was iteratively performed on the network with 1 x $10^6$ pseudo-random seeds. Both the most frequent solution and community co-occurrence frequency of each residue with p3V/P were mapped to H-2D$^b$/gp33.

### SUPPLEMENTARY FIGURES AND TABLES

**Table S1. Ensemble refinement statistics for each crystal structure.** [a] $\Delta R_{free}$ was calculated by subtracting the ensemble refinement $\Delta R_{free}$ from the published $\Delta R_{free}$. Multiple conformations were removed from H-2D$^b$/PF/P14 before ensemble refinement.

| Complex | PDB code | Space group | Resolution | $R_{free}$ | | $\Delta R_{free}$[a] | Ensemble size | Nr. AU molecules |
|---|---|---|---|---|---|---|---|---|
| | | | | PDB | Ensemble | | | |
| H-2D$^b$/gp33 | 1S7U | P 1 $2_1$ 1 | 2.20 Å | 0.262 | 0.280 | 0.018 | 27 | 4 |
| H-2D$^b$/V3P | 4NSK | C 1 2 1 | 2.60 Å | 0.283 | 0.267 | -0.016 | 30 | 1 |
| H-2D$^b$/Y4F | 1S7X | P 1 $2_1$ 1 | 2.41 Å | 0.254 | 0.246 | -0.008 | 24 | 4 |
| H-2D$^b$/PF | 3TBY | P 1 $2_1$ 1 | 2.50 Å | 0.315 | 0.311 | -0.004 | 18 | 4 |
| H-2D$^b$/ gp33/P14 | 5TJE | P $2_1$ $2_1$ $2_1$ | 3.20 Å | 0.314 | 0.334 | 0.020 | 14 | 2 |
| H-2D$^b$/ V3P/P14 | 5TIL | P $2_1$ $2_1$ $2_1$ | 2.83 Å | 0.275 | 0.295 | 0.020 | 14 | 2 |
| H-2D$^b$/PF/P14 | 5M02 | C 1 2 1 | 1.75 Å | 0.214 | 0.194 | -0.020 | 47 | 1 |

**Table S2. All polar contacts of each crystal copy in the TCR-unbound pMHCs.** All residues with favorable or unfavorable polar contacts within 4 Å are indicated.

| Complex | Copy | Chain | Polar contacts (<4.0 Å) |
|---|---|---|---|
| H-2Db/gp33 | A | H-2Db | E31 E58 E61 R62 Q65 K68 G69 Q70 Y84 Y85 N86 Q87 S88 A89 G90 G91 S92 H93 D106 R108 E119 R121 D137 M138 K186 T214 P267 |
| | | β2m | N42 E74 T75 T77 E89 D96 |
| | | gp33 | p4Y p5N |
| | B | H-2Db | Q54 E61 R62 Q65 G69 Q70 Y84 Y85 N86 Q87 S88 A89 G90 G91 S92 H93 R108 E119 R121 E137 M138 K173 R194 K196 E223 Y262 P267 |
| | | β2m | N42 K44 E74 T75 T77 E89 P90 D96 |
| | | gp33 | p4Y p5N |
| | C | H-2Db | R62 R79 Y84 Y85 N86 Q87 S88 A89 G90 G91 S92 W107 R108 E119 R121 D137 M138 K146 S150 N174 K186 R194 S195 E223 E264 |
| | | β2m | Q2 K19 N42 E74 T75 P90 T92 Y94 W95 D96 |
| | | gp33 | p4Y p8T |
| | D | H-2Db | G16 L17 E18 E53 Q54 R62 R79 Y84 Y85 N86 Q87 S88 A89 G90 G91 S92 H93 W107 R108 E119 R121 D137 M138 A139 K146 Q149 N176 K186 E223 E268 |
| | | β2m | Q2 N42 E74 T75 T92 Y94 W95 |
| | | gp33 | p4Y p8T |
| H-2Db/V3P | A | H-2Db | Q54 E58 Q65 R79 N80 N86 S88 G90 G91 R108 R121 A136 D137 M138 Q141 R144 K146 Q149 S150 H155 G162 E166 R170 K173 N220 E222 P267 E268 H269 R273 |
| | | β2m | Q2 N42 T75 D76 D85 S86 M87 A88 E89 W95 |
| | | V3P | p4Y p8T |
| H-2Db/Y4F | A | H-2Db | E41 E58 Q65 G69 Y84 Y85 N86 Q87 S88 A89 G90 G91 S92 H93 D106 W107 E119 R121 D137 M138 K173 R194 E223 P267 |
| | | β2m | N42 E74 T75 T77 |
| | | Y4F | p5N |
| | B | H-2Db | E41 R62 G69 Q70 Y84 Y85 N86 Q87 S88 A89 G90 G91 H93 E119 R121 D137 M138 E166 R194 S195 K196 E223 Y262 P267 E268 |
| | | β2m | N42 E74 T75 T77 D96 R97 |
| | | Y4F | p5N |
| | C | H-2Db | G16 E18 Q54 R75 R79 N80 Y84 Y85 N86 Q87 S88 A89 G90 G91 S92 H93 W107 R108 E119 R121 A136 D137 M138 A139 K146 K173 K186 S195 E222 |
| | | β2m | Q2 K19 G43 E74 T75 |
| | | Y4F | p8T |
| | D | H-2Db | E18 E53 Y84 Y85 N86 Q87 S88 A89 G90 G91 S92 H93 W107 R108 E119 R121 D137 M128 A136 K146 S150 N174 T178 S195 H263 P267 E268 |
| | | β2m | H13 K19 E74 T75 W85 T92 |
| | | Y4F | p8T |
| H-2Db/PF | A | H-2Db | E58 R62 Q65 K68 G69 Q70 Q72 Y84 Y85 N86 Q87 S88 G90 G91 S92 H93 W107 E119 R121 D137 M138 N176 R181 R194 T214 P267 |
| | | β2m | E16 N42 T75 T77 E89 K91 T92 |
| | | PF | p5N |
| | B | H-2Db | R62 Q65 G69 Q70 Q72 Y84 Y85 N86 Q87 S88 G90 G91 S92 H93 W107 E119 R121 D137 M138 T178 R181 R194 E223 H263 P267 |
| | | β2m | E16 N42 E74 T77 E89 K91 T92 Y94 W95 |
| | | PF | p5N |
| | C | H-2Db | E18 R62 Q72 R79 Y84 Y85 N86 Q87 S88 G90 G91 S92 H93 S105 W107 R108 E119 R121 D137 M138 K146 N176 R181 K186 P267 |
| | | β2m | N17 E74 T75 E89 Y94 R97 |
| | | PF | p8T |
| | D | H-2Db | E18 R62 Q72 Y84 Y85 N86 Q87 S88 G90 G91 S92 H93 W107 R108 E119 R121 D137 M138 K146 S150 G175 R181 K186 P267 |
| | | β2m | Q2 N17 T73 E74 T75 S86 E89 Y94 W95 D96 R97 |
| | | PF | p8T |

**Table S3. All polar contacts of each crystal copy in the TCR-bound pMHCs.** All residues with favorable or unfavorable polar contacts within 4 Å are indicated.

| Complex | Copy | Chain | Polar contacts (<4.0 Å) |
|---|---|---|---|
| H-2D$^b$/gp33/P14 | A | TCRα | S17 Q16 T19 T25 V60 D62 K117 |
| H-2D$^b$/gp33/P14 | A | TCRβ | H39 T53 K121 V122 L124 E126 K175 N178 Y179 R198 N227 S229 E231 A230 |
| H-2D$^b$/gp33/P14 | A | H-2D$^b$ | L17 E18 N86 K103 D106 W107 R145 H169 T214 N220 G221 E264 L266 P267 E268 R273 |
| H-2D$^b$/gp33/P14 | A | β2m | K19 E36 Q38 N42 G43 K44 K45 R81 Y94 |
| H-2D$^b$/gp33/P14 | A | gp33 | – |
| H-2D$^b$/gp33/P14 | B | TCRα | Q1 K3 E4 K5 V60 D62 R76 K117 |
| H-2D$^b$/gp33/P14 | B | TCRβ | D36 H39 R42 D51 T53 E54 K175 E176 S177 N178 Y181 V122 L124 N159 G160 R198 H200 Q226 R235 D237 |
| H-2D$^b$/gp33/P14 | B | H-2D$^b$ | L17 E18 D106 W107 R144 E148 K173 T182 D183 T214 N220 G221 L266 P267 E268 |
| H-2D$^b$/gp33/P14 | B | β2m | P20 N21 E36 Q38 K44 R81 |
| H-2D$^b$/gp33/P14 | B | gp33 | – |
| H-2D$^b$/V3P/P14 | A | TCRα | Q16 S17 T19 T25 V60 S61 D62 K117 |
| H-2D$^b$/V3P/P14 | A | TCRβ | D36 T37 Y48 D51 K63 K175 N178 Y179 K121 V122 L124 E126 R198 N227 I228 S229 E231 R235 |
| H-2D$^b$/V3P/P14 | A | H-2D$^b$ | L17 E18 Y84 Y85 N86 D106 W107 R108 R145 K173 T214 N220 G221 E223 L266 P267 |
| H-2D$^b$/V3P/P14 | A | β2m | K19 E36 Q38 K44 K45 T75 D76 R81 Y94 |
| H-2D$^b$/V3P/P14 | A | V3P | – |
| H-2D$^b$/V3P/P14 | B | TCRα | Q2 E4 H6 D7 V60 D62 |
| H-2D$^b$/V3P/P14 | B | TCRβ | D36 R42 D51 T53 E54 K63 V122 S123 L124 E126 N159 G160 Y174 K175 E176 N178 R198 R202 Q226 D237 |
| H-2D$^b$/V3P/P14 | B | H-2D$^b$ | L17 E18 D106 W107 R108 R144 E148 K173 R181 T182 T214 N220 G221 E223 L266 P267 |
| H-2D$^b$/V3P/P14 | B | β2m | H13 P20 N21 E36 Q38 K44 K45 T75 R81 Y94 |
| H-2D$^b$/V3P/P14 | B | V3P | – |
| H-2D$^b$/PF/P14 | A | TCRα | V60 S61 D62 K63 R134 Q136 T155 |
| H-2D$^b$/PF/P14 | A | TCRβ | T13 D51 T53 E54 E112 D113 R115 K129 N134 N158 N178 Y179 R198 E218 R202 K222 Q226 S229 A236 |
| H-2D$^b$/PF/P14 | A | H-2D$^b$ | L17 E18 E53 Q54 E58 D106 W107 R108 K131 A135 A136 D137 M138 Q141 R145 H169 K157 K173 H191 G197 T214 N220 G221 E222 E223 Q255 N256 Y262 L266 P267 |
| H-2D$^b$/PF/P14 | A | β2m | E36 Q38 N42 K44 K45 E74 R81 Y94 |
| H-2D$^b$/PF/P14 | A | PF | – |

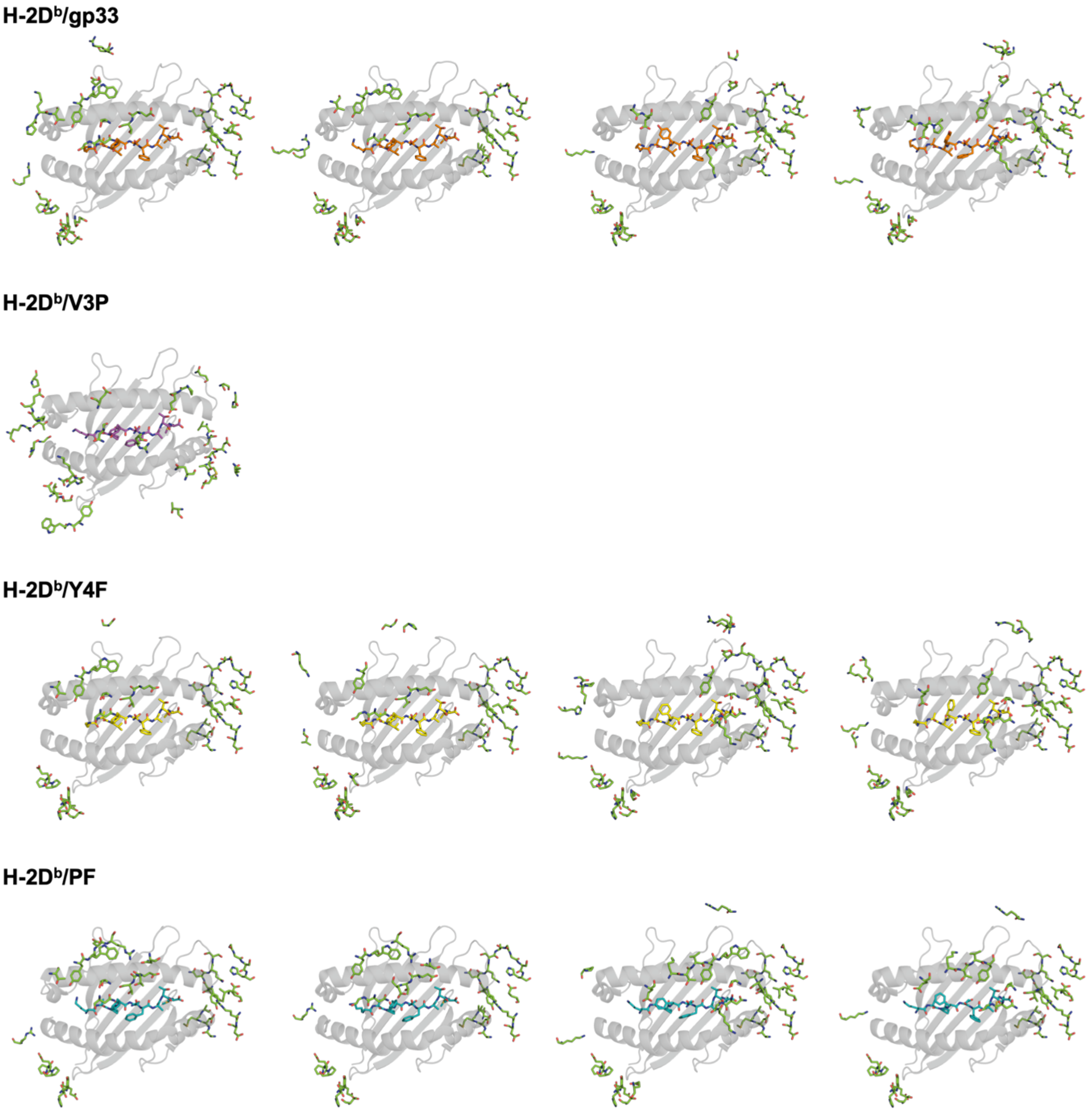


**Fig. S1. The crystal packing geometry at the pMHC interface is similar for H-2D$^b$/gp33, H-2D$^b$/Y4F, and H-2D$^b$/PF.** The α1 and α2 domains are shown as a transparent gray cartoon. Each peptide is shown in stick representation, with gp33 (orange), V3P (purple), Y4F (yellow), and PF (teal) color accordingly. Any residues not part of the respective crystal copy within 4 Å (green) are in stick representation.

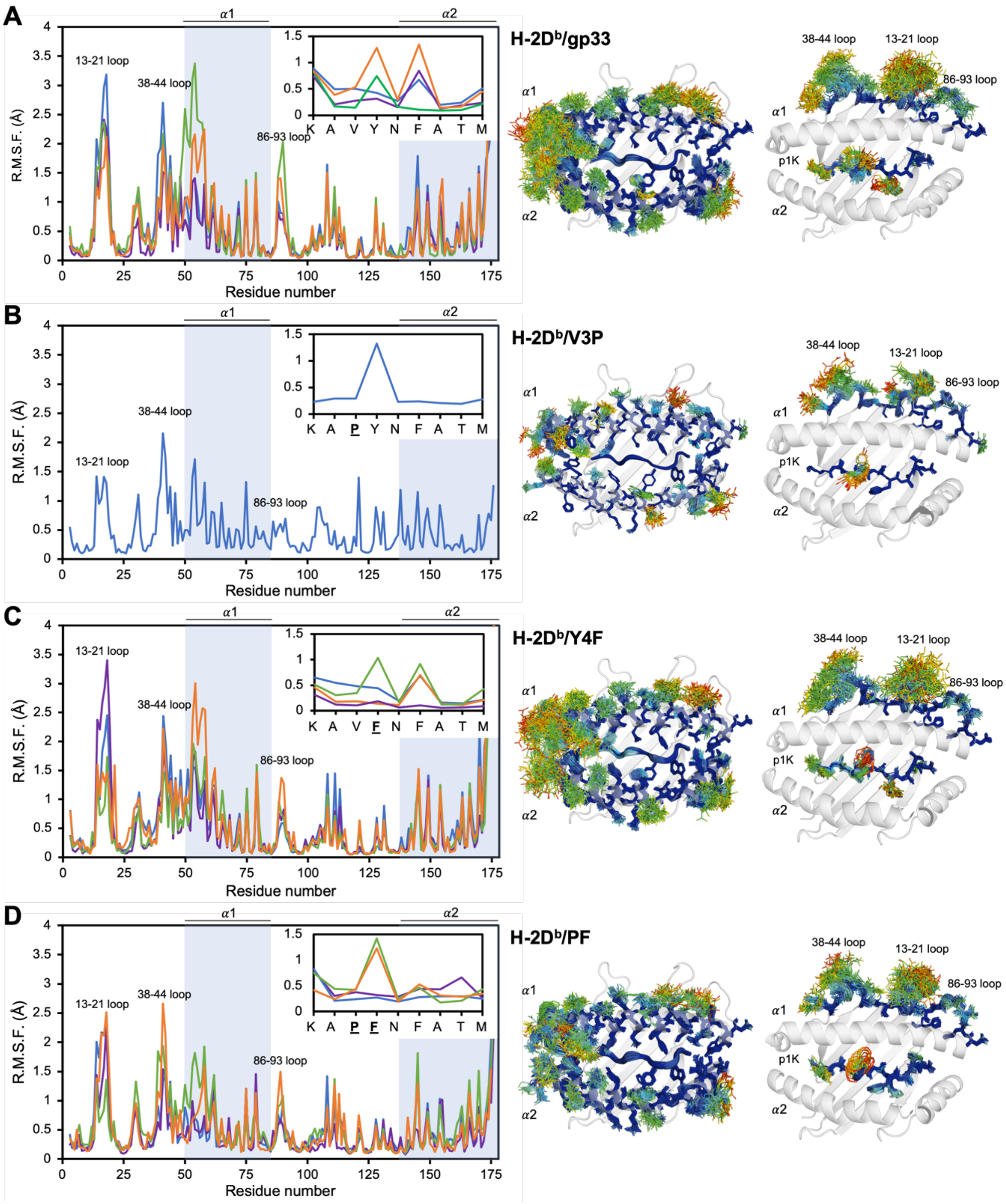


**Fig. S2. MHC and peptide conformational flexibility are reduced when proline is positioned at p3.** The per-residue RMSF values, obtained from ensemble refinement, are reported for residues 1 to 176 of MHC and all peptide residues (inset). MHC helixes and most dynamic loop regions are labeled. Data are reported using different line colors for every molecule in the asymmetric unit (blue for copy A, purple for copy B, green for copy C and orange for copy D). Ensemble structural regions are represented as sticks and colored by RMSF, ranging from 0 (deep blue) to 4 (bright red) Å. pMHC structures represent the complex before P14 binding. **A** A high degree of mobility is seen in H-2D$^b$/gp33, particularly at the beginning of the α1 helix, and by the loops, including residues 13-21 and 38-44. **B** The pMHC fluctuations are drastically reduced in H-2D$^b$/V3P. The only residue of the peptide exploring multiple conformations is p4Y. **C** A similar degree of dynamics is observed in H-2D$^b$/Y4F compared to H-2D$^b$/gp33. However, p4F and p6F display reduced flexibility. **D** The dynamics of the pMHC are reduced in H-2D$^b$/PF, despite p4F regaining mobility.

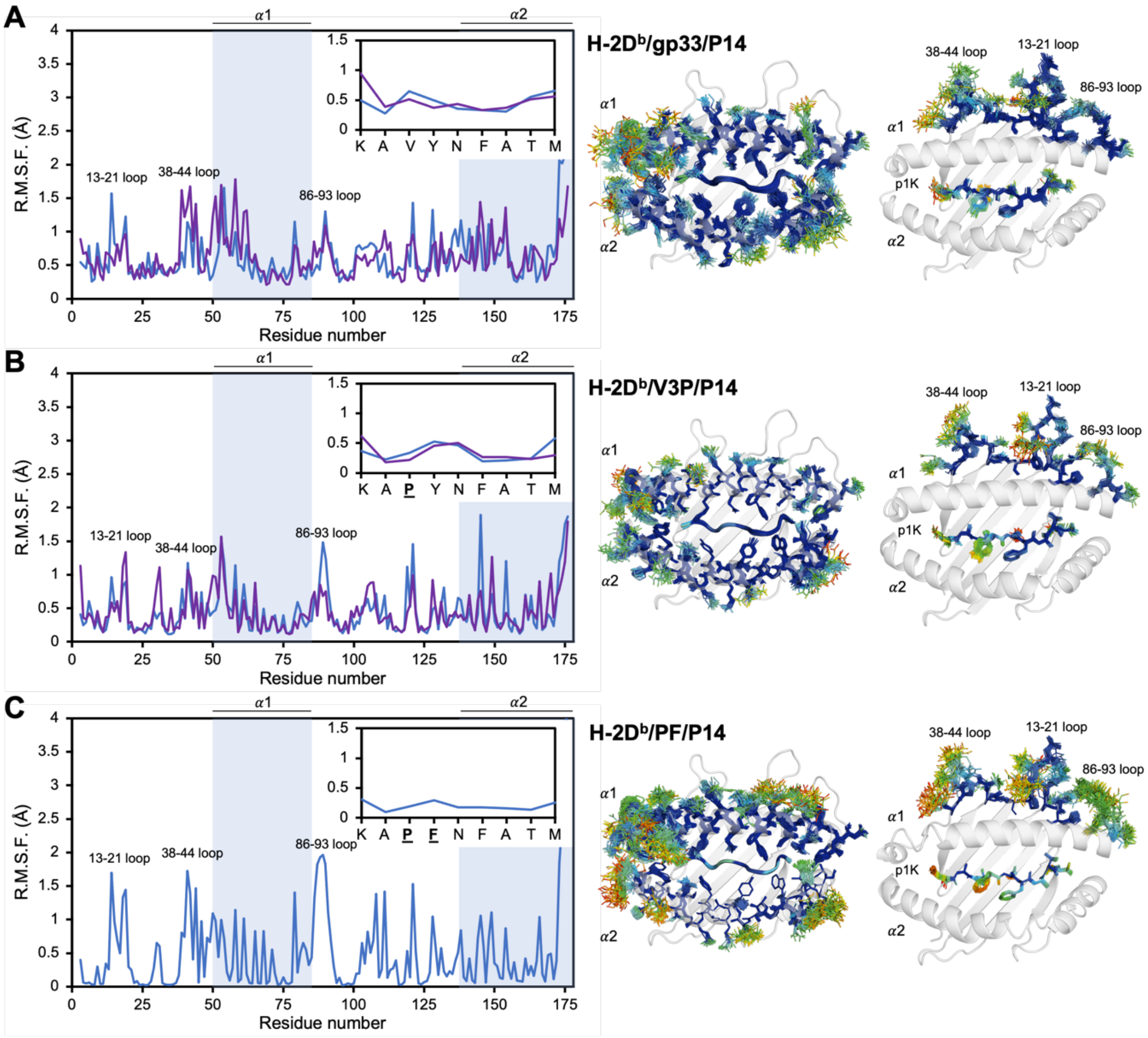


**Fig. S3. MHC and peptide conformational flexibility are reduced when the TCR P14 binds to the complexes.** The per-residue RMSF values, obtained from ensemble refinement, are reported for residues 1 to 176 of MHC and all peptide residues (inset). MHC helixes and the most flexible and dynamic loop regions are labeled. Data are plotted with different line colors for every molecule in the asymmetric unit (blue for copy A, purple for copy B). Ensemble structural regions are represented as sticks and colored by RMSF, ranging from 0 (deep blue) to 4 (bright red) Å. All pMHC structures represent the complex after P14 binding. **A** Reduced dynamics are observed in H-2D$^b$/gp33 after P14 binding. **B** The fluctuations in the pMHC are further reduced in H-2D$^b$/V3P.

**C** Increased mobility is visible around the region of α1 helix in H-2D$^{b}$/PF, particularly in the loop containing residues 86-96. Overall, the pMHC complexes bound to P14 display reduced dynamics compared to the free pMHC complexes.

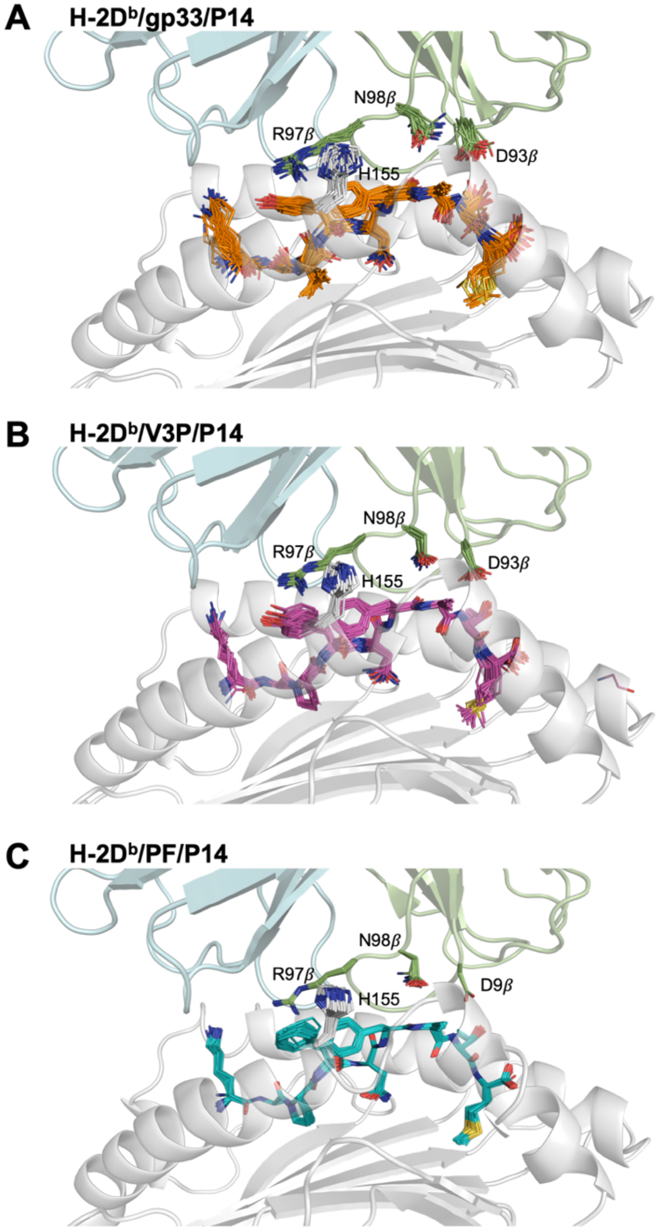


**Fig. S4. The TCR-peptide interface is rigid and is no observable relationship between pMHC-TCR affinity and dynamics in this region of the ternary complex. A**, **B**, **C** Cartoon representation of the MHC heavy chain (grey) and P14 α (light blue) and β (green) chains. Peptide residues, a key interacting residue of the MHC (H155), and TCR (D93β, R97β, and N98β) are shown as sticks. The conformations of the labeled residues are similar in all ensembles, with H-2D$^b$/gp33/P14 displaying the greatest degree of fluctuations.

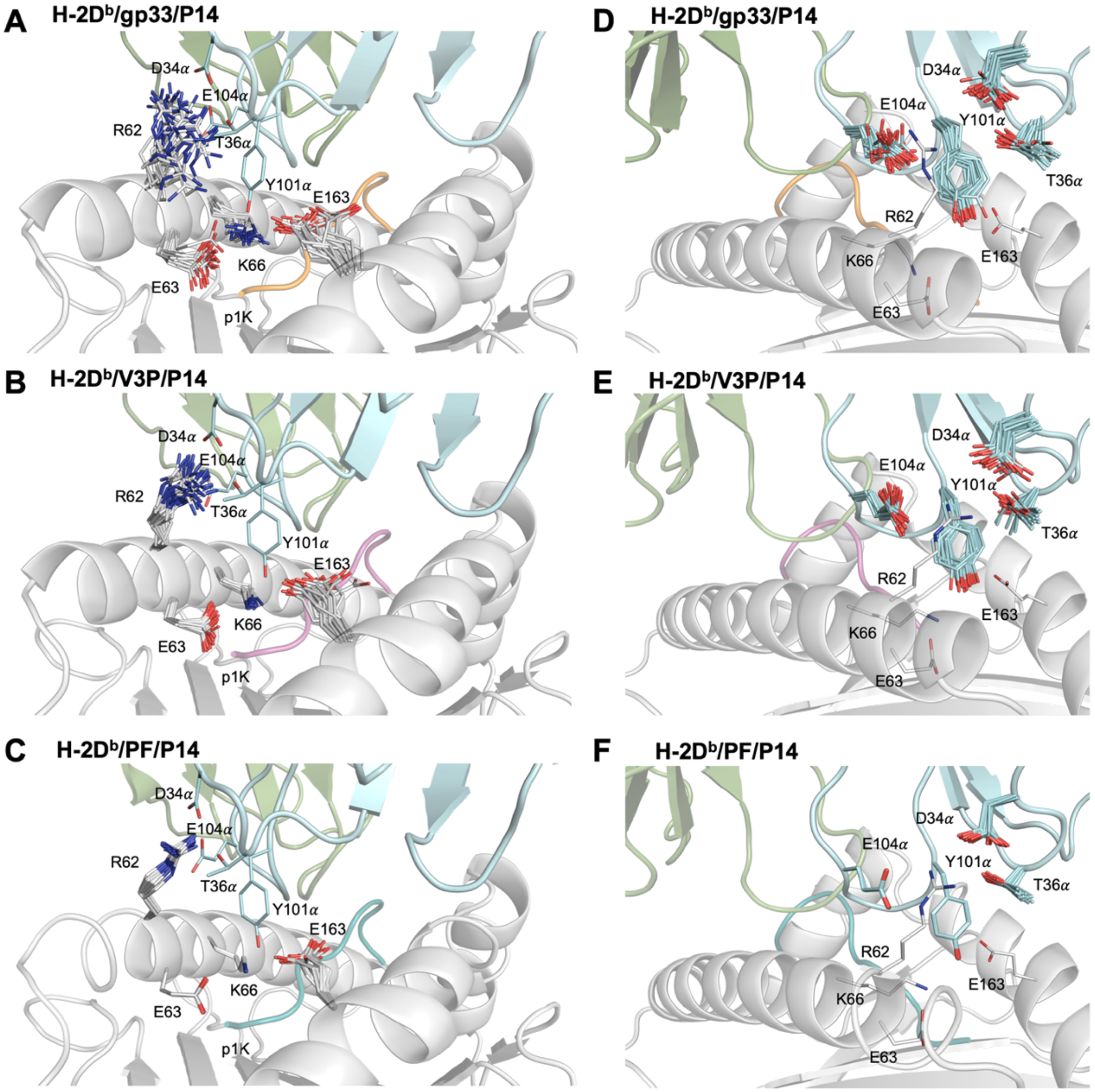


**Fig. S5. The conformational rigidity at the pMHC-TCR interface does not correlate with the affinity of the TCR-pMHC interactions.** **A**, **B**, **C**, **D**, **E**, **F** Cartoon representation of the heavy chain (grey), peptides and P14 α (light blue) and β (green) chains. Key interacting residues of the MHC (R62, E63, K66, E163) and the TCR (D34α, T36α, Y101α, E104α) are shown as sticks. Sidechain ensembles of MHC heavy chain residues are shown in **A**, **B**, and **C**, while sidechain ensembles of TCR α chain residues are shown in **D**, **E**, and **F**. The conformation of the highlighted residues is similar in all ensembles of complexes, with H-2D$^b$/gp33/P14 and H-2D$^b$/V3P/P14 having R62 and E163 displaying higher conformational flexibility.

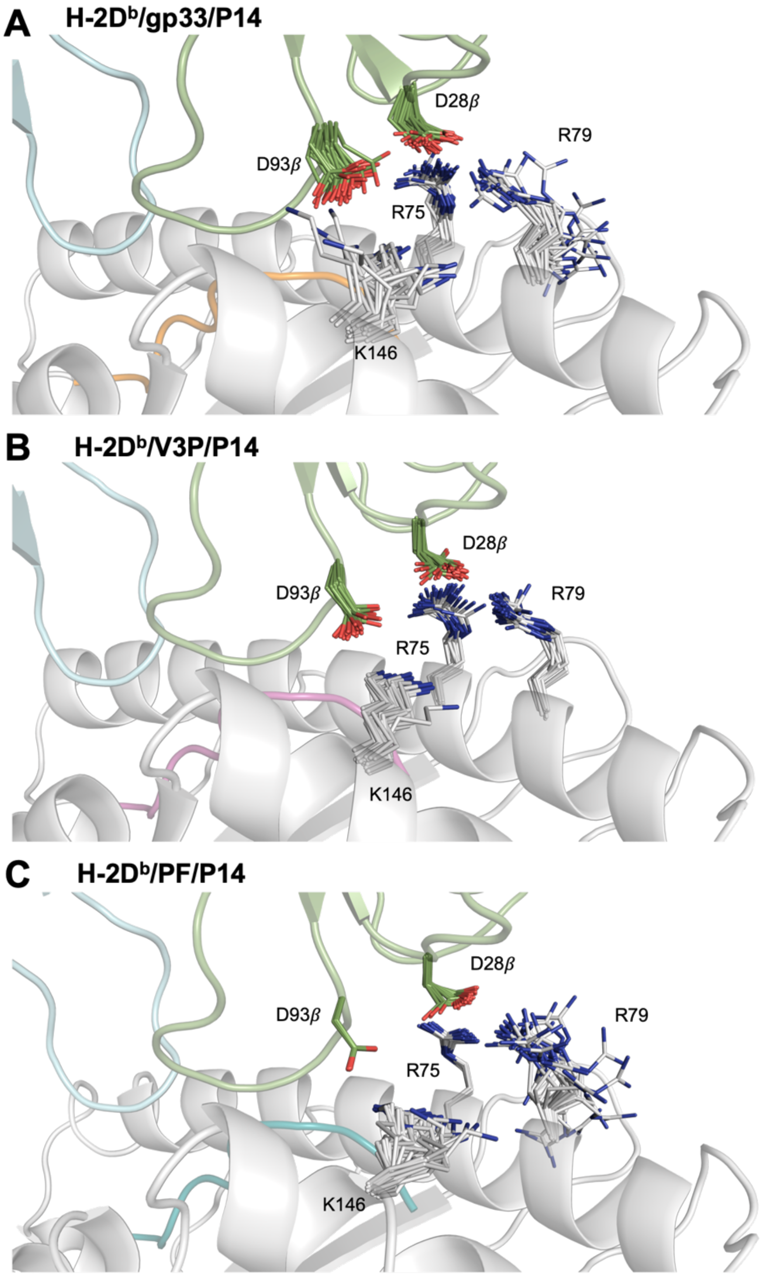


**Fig. S6. The conformational rigidity at the pMHC-TCR interface does not correlate with the affinity of the TCR-pMHC interactions. A**, **B**, **C** Cartoon representation of the heavy chain (grey), peptides, and P14 α (light blue) and β (green) chains. The sidechains of the key interacting MHC (R75, R79, K146) and TCR (D93β, D28β) residues are shown as sticks. R79 and K146 sidechains in H-2D$^b$/gp33/P14 and H-2D$^b$/PF/P14 complexes display increased conformational flexibility compared to H-2D$^b$/V3P/P14.

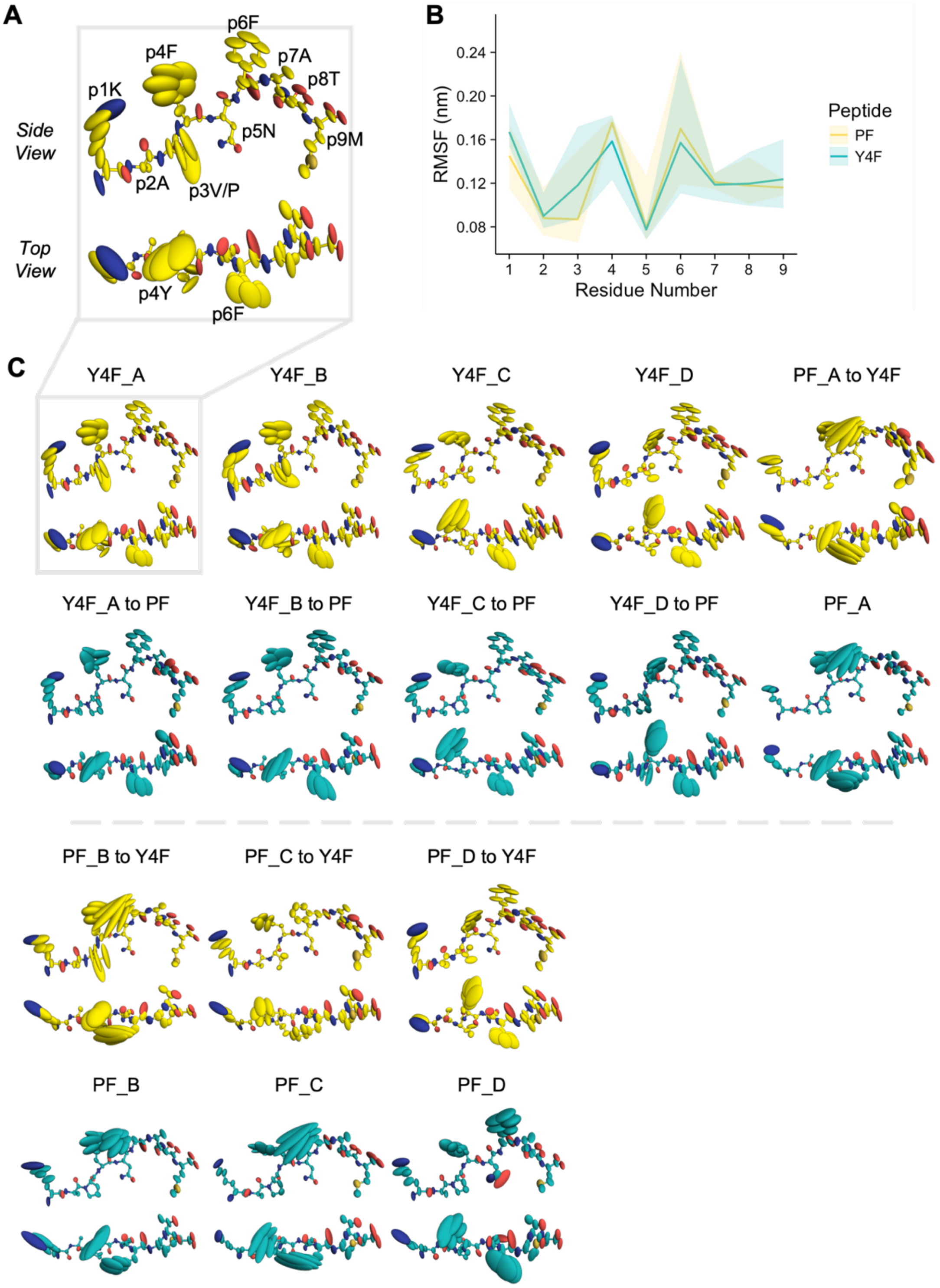


**Fig. S7. Overview of the conformational sampling of the Y4F and PF peptides throughout the MD trajectories reveals that the proline substitution leads to rigidification at p3 and increased dynamics at p6F.** Stick and ellipsoid representation of MHC-bound Y4F (yellow) and

PF (purple). Ellipsoids are scaled and shaped according to the anisotropic atomic temperature factors derived from each trajectory and mapped to their respective crystallographic models. **A** Residue labeling of side and top views of the peptide as reference. **B** Average per-residue RMSF values over all trajectories for Y4F and PF. The colored transparent envelopes depict the full range of observed trajectory RMSF values for each peptide. **C** Conformational sampling of the peptides observed in each trajectory. Paired simulations are organized vertically, with one trajectory derived from the initial crystallographic model and the other derived from an *in silico* mutation between p3V and p3P.

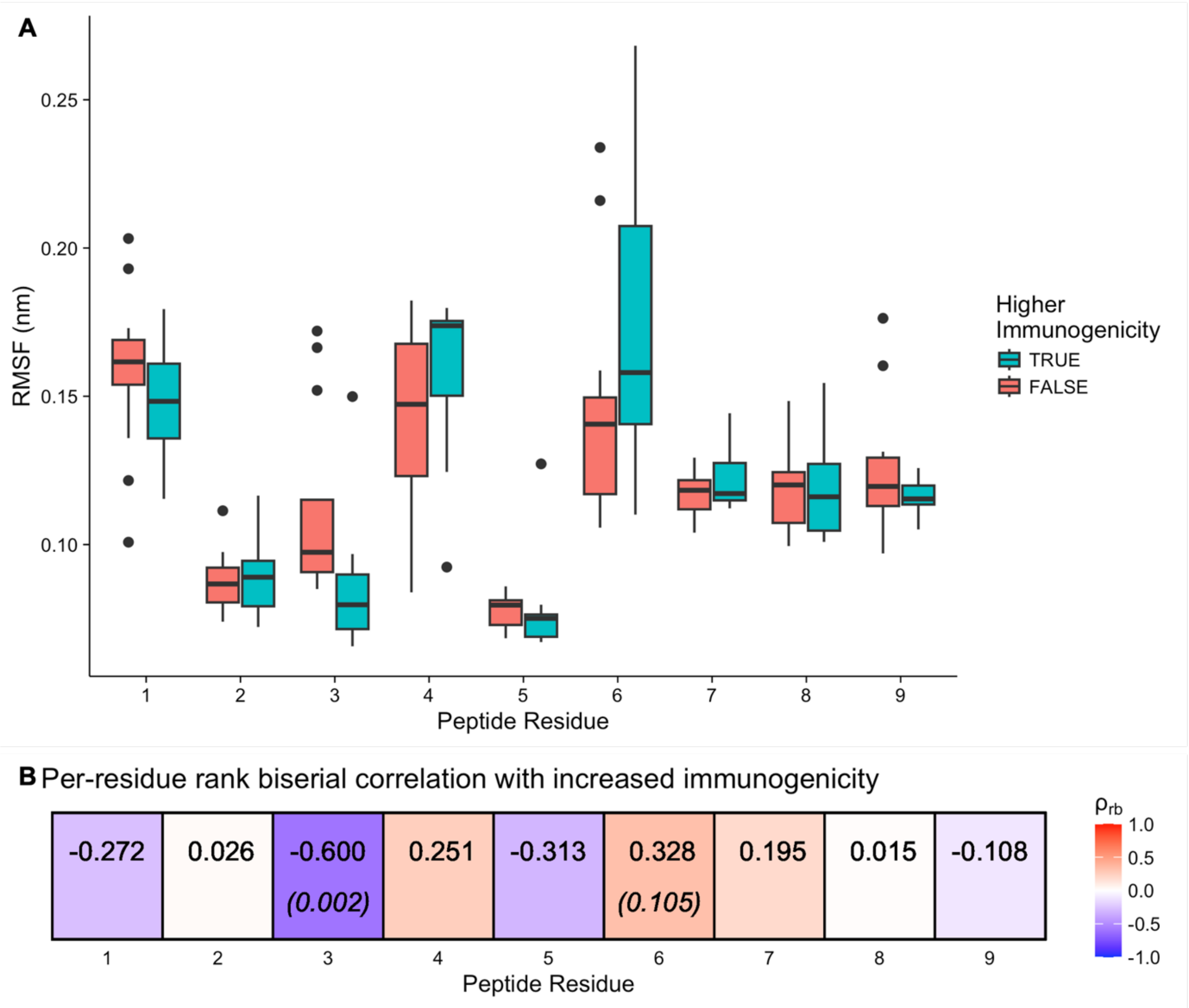


**Fig. S8. MD-derived per-residue peptide dynamics correlate with immunogenicity.** MD trajectories were grouped according to their relative immunogenicity, that is the p3P trajectories were considered as more immunogenic than their paired p3V trajectory counterparts, reflecting previously published findings (*62*). **A** Boxplots summarizing the observed RMSF values per trajectory. Visually, p3V/P and p6F display the most striking differences. **B** Rank biserial correlation of per-residue RSMF values and immunogenicity. The uncorrected p-values, if calculated and obtained via permutation testing, are indicated in parentheses. In this comparison, p3V/P and p6F are also the strongest correlates of immunogenicity.

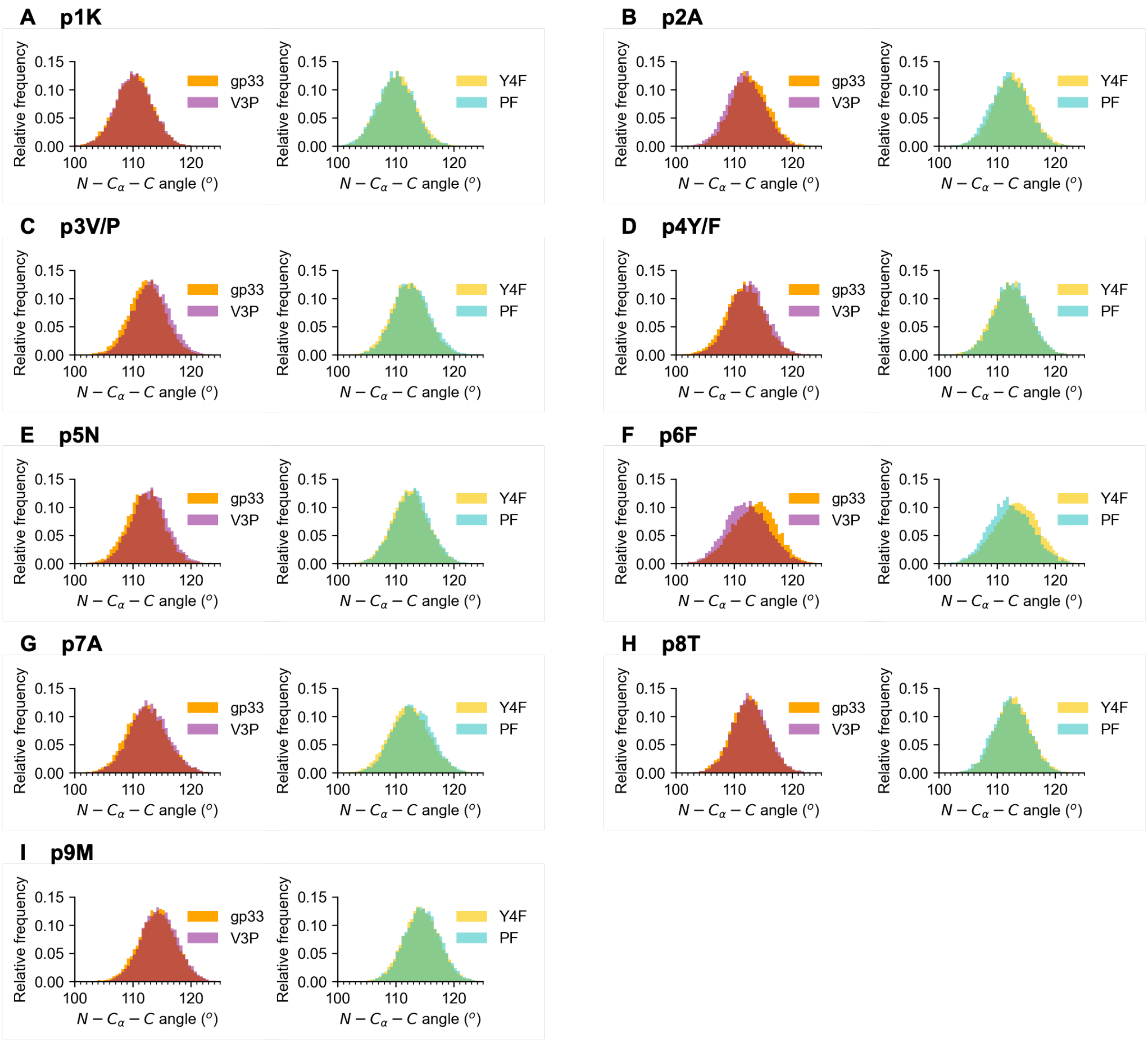


**Fig. S9. The observed backbone angles (N-Cα-C) of each peptide residue throughout the MD trajectories imply that the altered dynamics at p3 are not transmitted to p6 via the peptide backbone. A, B, C, D, E, F, G, H, I** Relative frequency histograms of the backbone angles of each indicated peptide residue. MD trajectories were grouped according to peptide. Only p6F (**F**) shows a distinct difference in backbone angle when comparing p3V with p3P peptides.

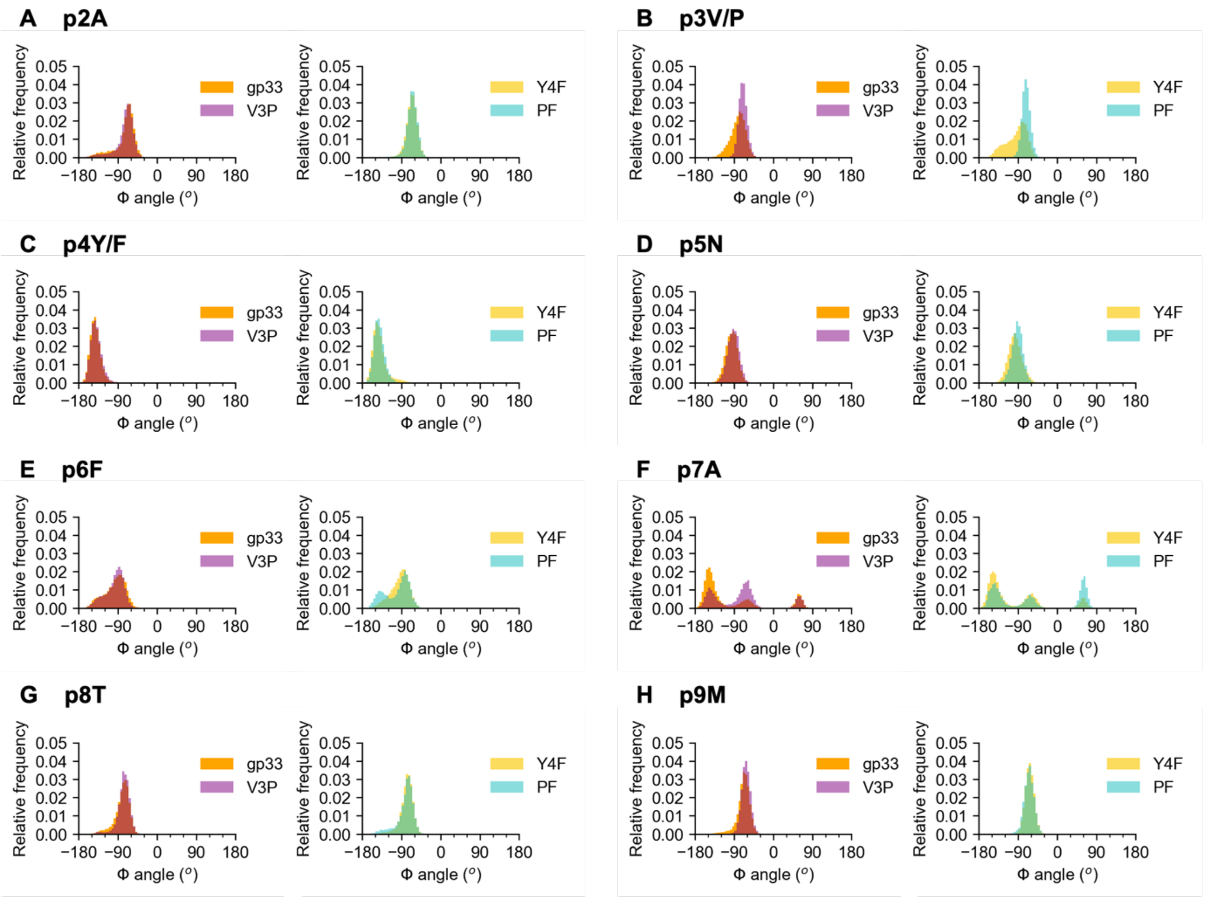


**Fig. S10. The observed Φ dihedral angles (C-N-C$_{\alpha}$-C) of each peptide residue throughout the MD trajectories imply that the altered dynamics at p3 are not transmitted to p6 via the peptide backbone. A**, **B, C, D, E, F, G, H** Relative frequency histograms of the Φ angles of each indicated peptide residue. MD trajectories were grouped according to peptide. The Φ angles of p3V/P (**B**) and p7A (**F**) show distinct differences in conformational sampling. Most notably, p4Y/F (**C**) and p5N (**D**) remain similar, regardless of the presence of a proline or valine at the third peptide position.

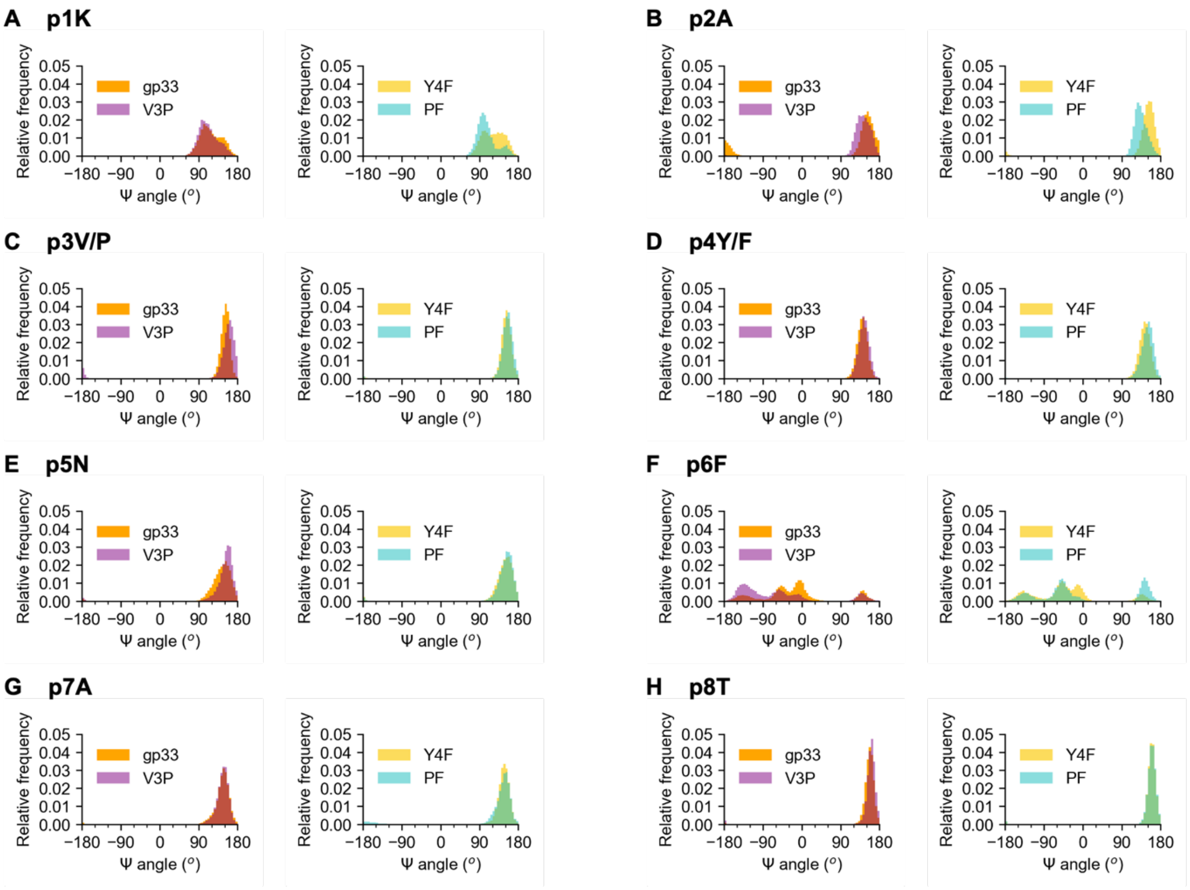


**Fig. S11 The observed Ψ dihedral angles (N-$C_\alpha$-C-N) of each peptide residue throughout the MD trajectories imply that the altered dynamics at p3 are not transmitted to p6 via the peptide backbone. A**, **B, C, D, E, F, G, H** Relative frequency histograms of the Ψ angles of each indicated peptide residue. MD trajectories were grouped according to peptide. The Ψ angles of p2A (**B**) and p6F (**F**) show distinct differences in conformational sampling. Most notably, p4Y/F (**D**) and p5N (**E**) remain similar, regardless of the presence of a proline or valine at the third peptide position.

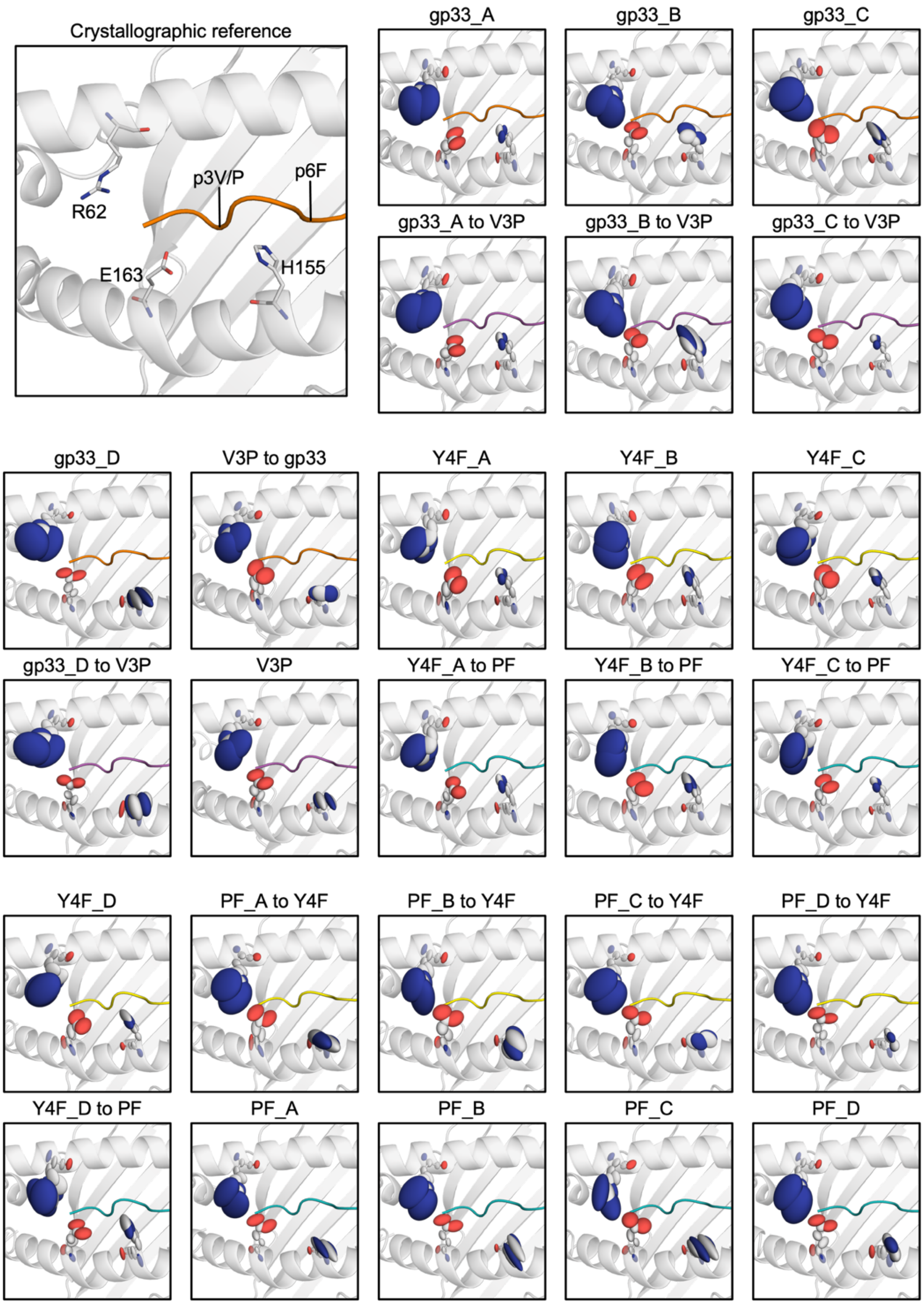


**Fig. S12. Visual overview of R62, H155, and E163 sidechain dynamics throughout all trajectories.** Stick and ellipsoid representation of each sidechain of interest. Ellipsoids are scaled and shaped according to the anisotropic atomic temperature factors derived from each trajectory and mapped to their respective crystallographic models. The backbone of H-2D$^b$ (white), gp33 (orange), V3P (purple), Y4F (yellow), and PF (teal) are shown in cartoon representation. Trajectories are paired vertically, one derived from the initial crystallographic model and the other derived from an *in silico* mutation between p3V and p3P. No relationship between the p3P substitution and R62 sidechain dynamics is visually evident. On average, increased conformational sampling of H155 is observed in the presence of p3P. In the trajectories where H155 is more dynamic, p6F is generally more dynamic (**Figs. 2, S7**). E163, which is near p3 in the structure, displays smaller fluctuations in the presence of p3P compared to p3V.

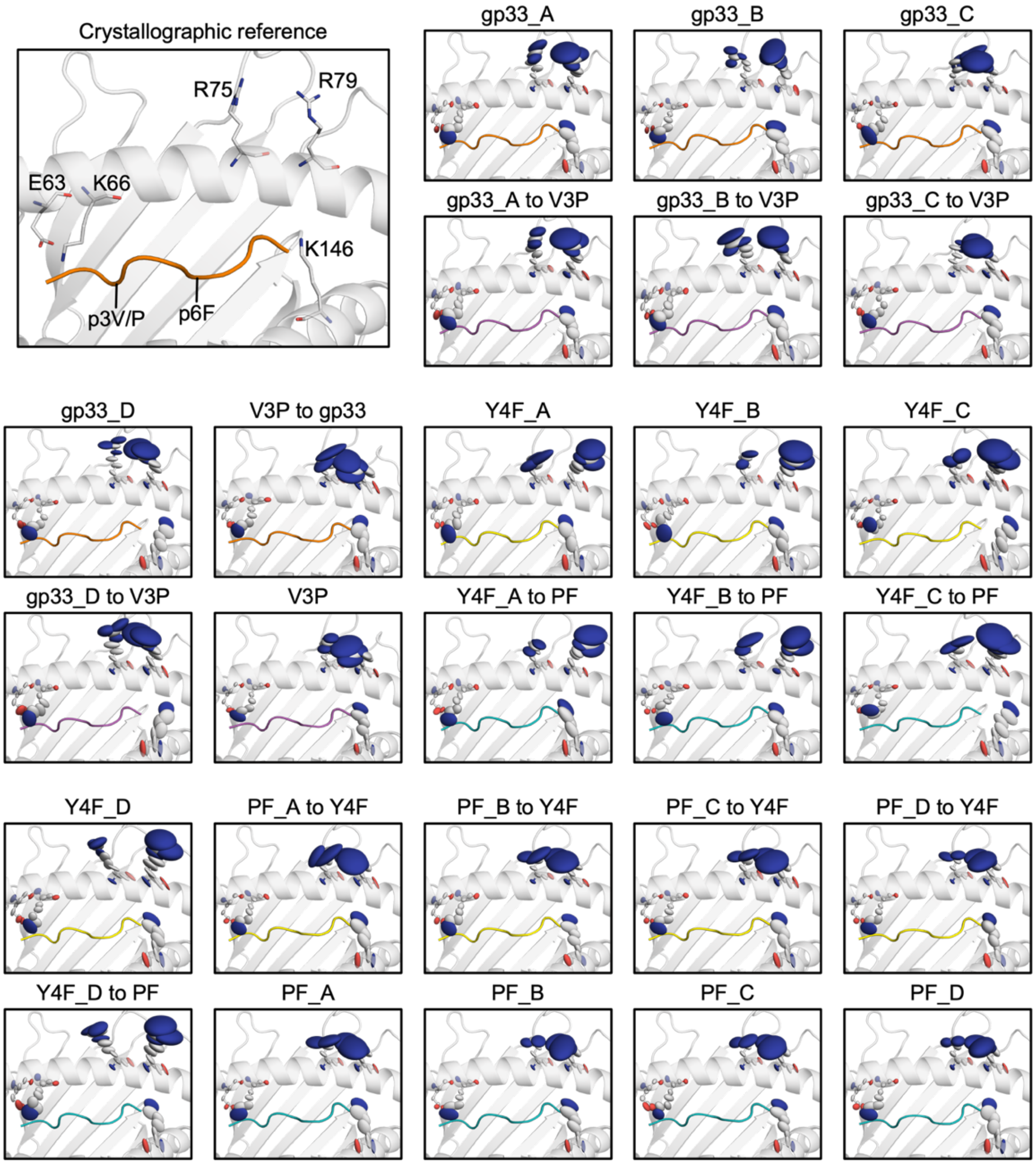


**Fig. S13. Visual overview of E63, K66, R75, R79 and K146 sidechain dynamics throughout all trajectories.** Stick and ellipsoid representation of each sidechain of interest. Ellipsoids are scaled and shaped according to the anisotropic atomic temperature factors derived from each trajectory and mapped to their respective crystallographic models. The backbone of H-2D$^b$ (white), gp33 (orange), V3P (purple), Y4F (yellow), and PF (teal) are shown in cartoon representation. Trajectories are paired vertically, one derived from the initial crystallographic model and the other derived from an *in silico* mutation between p3V and p3P. Visually, no consistent differences in sidechain fluctuations for these residues of interest when comparing p3V and p3P trajectories were identified.

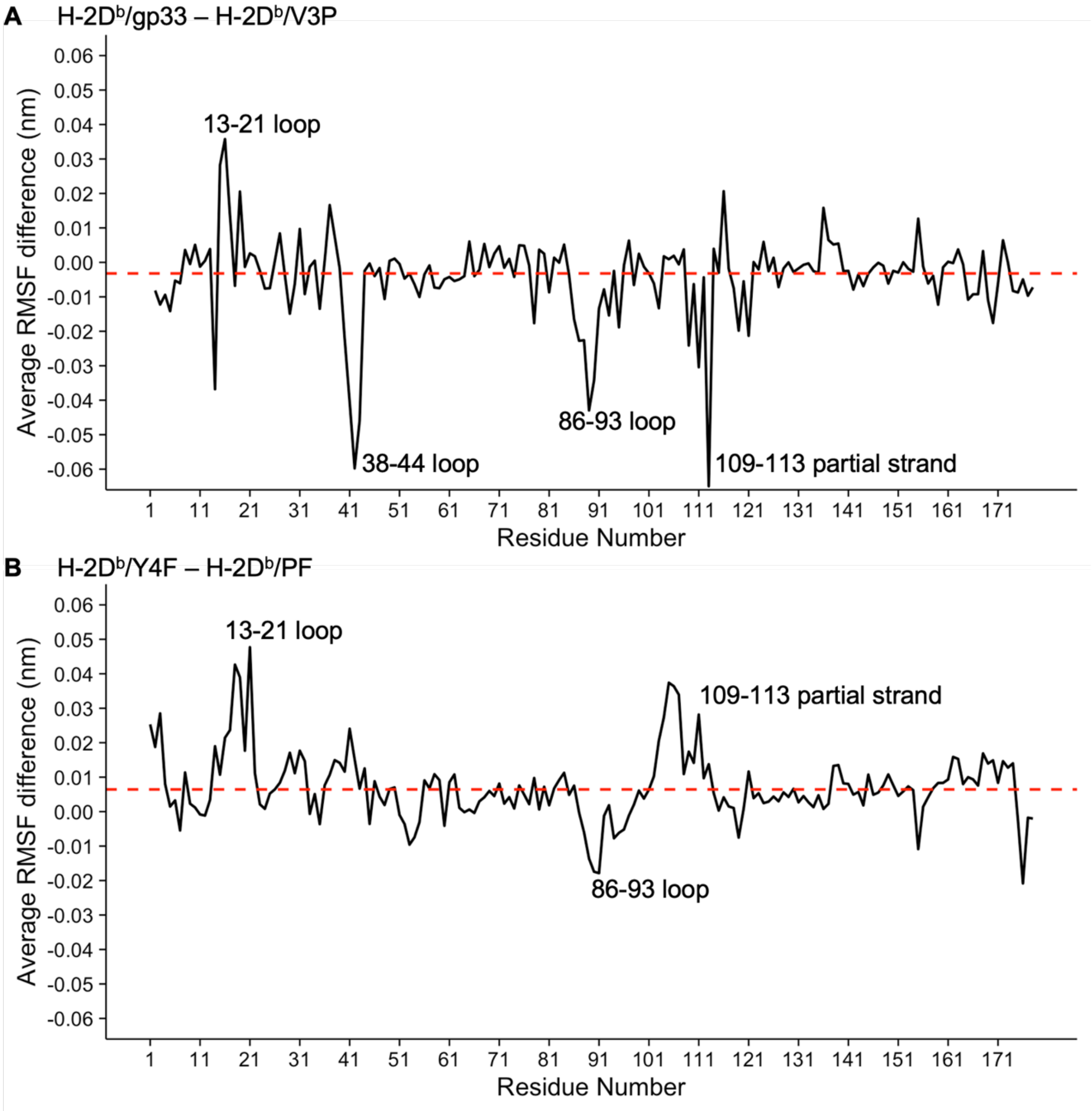


**Fig. S14. Differences in average per-residue RMSF values for H-2D$^b$.** MD trajectories were grouped according to peptide to obtain the average per-residue RMSF. The difference in average per-residue RMSF was calculated by subtracting the values of comparable trajectories: **A** H-2D$^b$/gp33 – H-2D$^b$/V3P and **B** H-2D$^b$/Y4F – H-2D$^b$/PF. Dashed red lines denote the average difference. The regions of interest identified in ensemble refinements (**Figs. S2, S3**) are labeled.

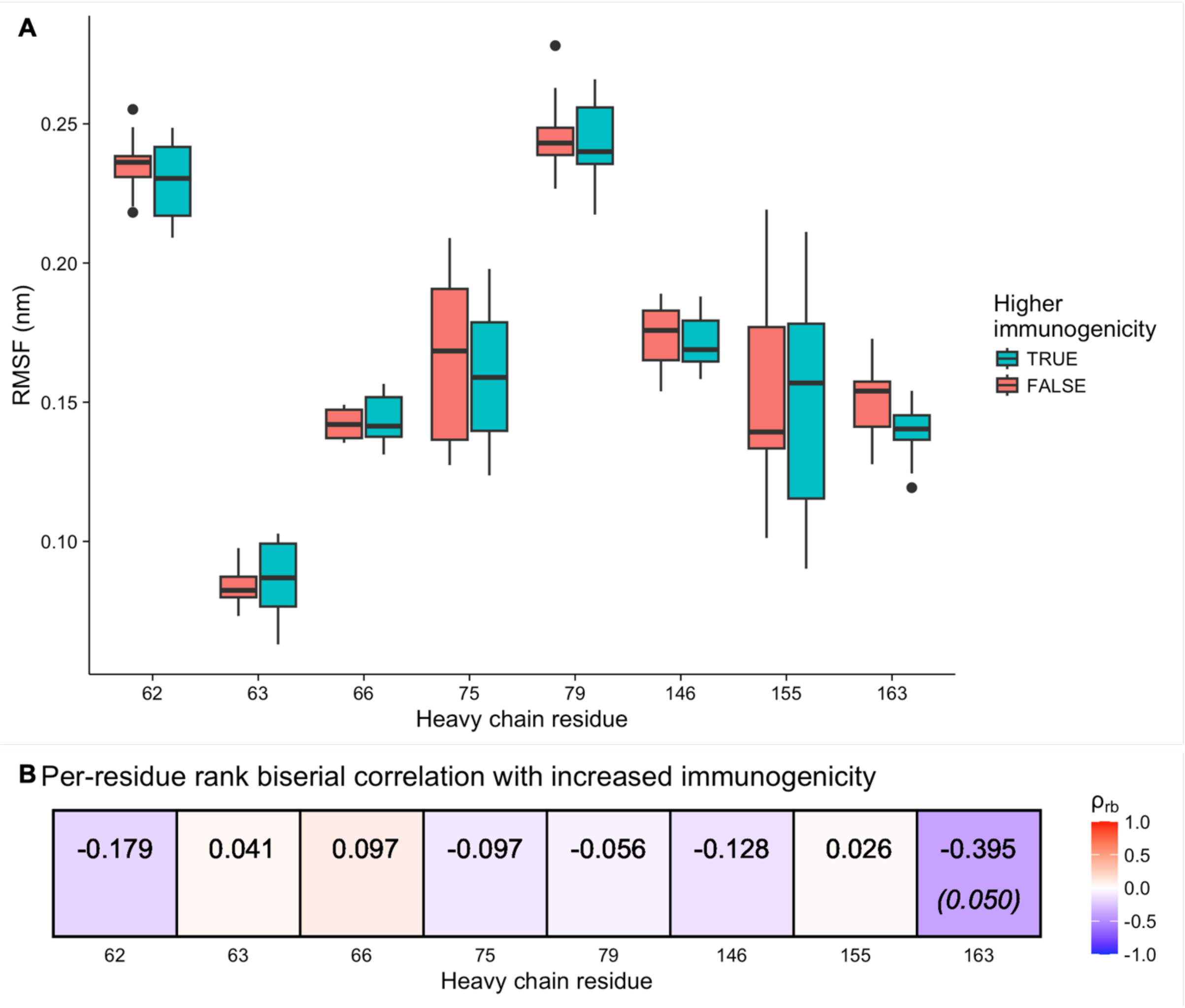


**Fig. S15. Assessing the correlation of H-2D$^b$ dynamics with immunogenicity.** MD trajectories were grouped according to their relative immunogenicity, that is the p3P trajectories were considered more immunogenic than their paired p3V trajectory counterparts, reflecting previously published findings (*62*). **A** Boxplots summarizing the observed RMSF values per trajectory. Visually, E163 displays the largest difference. **B** Rank biserial correlation of per-residue RSMF values and immunogenicity. The uncorrected p-values, if calculated and obtained via permutation testing, are indicated in parentheses. In this comparison, E163 is also the strongest correlate of immunogenicity.

**Table S4. Residues that are significantly correlated with each peptide position.** Table summarizes identified significant correlates in **Fig. 3**.

| Peptide residue | Significantly correlated residues |
|---|---|
| p1K | R121 E163 |
| p2A | S4 M5 Y7 A11 Y45 E58 R62 E63 K66 G69 S99 G100 C101 D102 G112 G120 K157 A158 Y159 L160 E161 C164 p3V/P |
| p3V/P | G16 L160 p2A |
| p4Y/F | – |
| p5N | W73 L114 F116 A117 I124 L130 K131 T132 W133 T134 R145 W147 E148 S150 G151 A152 A153 E154 Y156 p7A p8T p9M |
| p6F | H155 R170 K173 T178 |
| p7A | W73 L114 F116 A117 E119 D122 Y123 I124 A125 L126 E128 D129 L130 K131 T132 W133 T134 A135 A140 A141 I142 T143 R144 R145 K146 W147 E148 Q149 S150 G151 A152 A153 E154 Y156 K157 p5N p8T p9M |
| p8T | W73 V76 S77 F116 A117 Y118 D122 Y123 I124 A125 E128 D129 T132 W133 T134 A135 A140 Q141 I142 T143 R144 R145 K146 W147 E148 Q149 S150 A152 p5N p7A p9M |
| p9M | P47 A49 W73 V76 S77 N80 L81 Y84 L103 F116 A117 Y118 D122 Y123 I124 A125 W133 T134 A135 D137 M138 A139 A140 Q141 I142 T143 R144 R145 K146 W147 E148 S150 G151 A152 p5N p7A p8T |

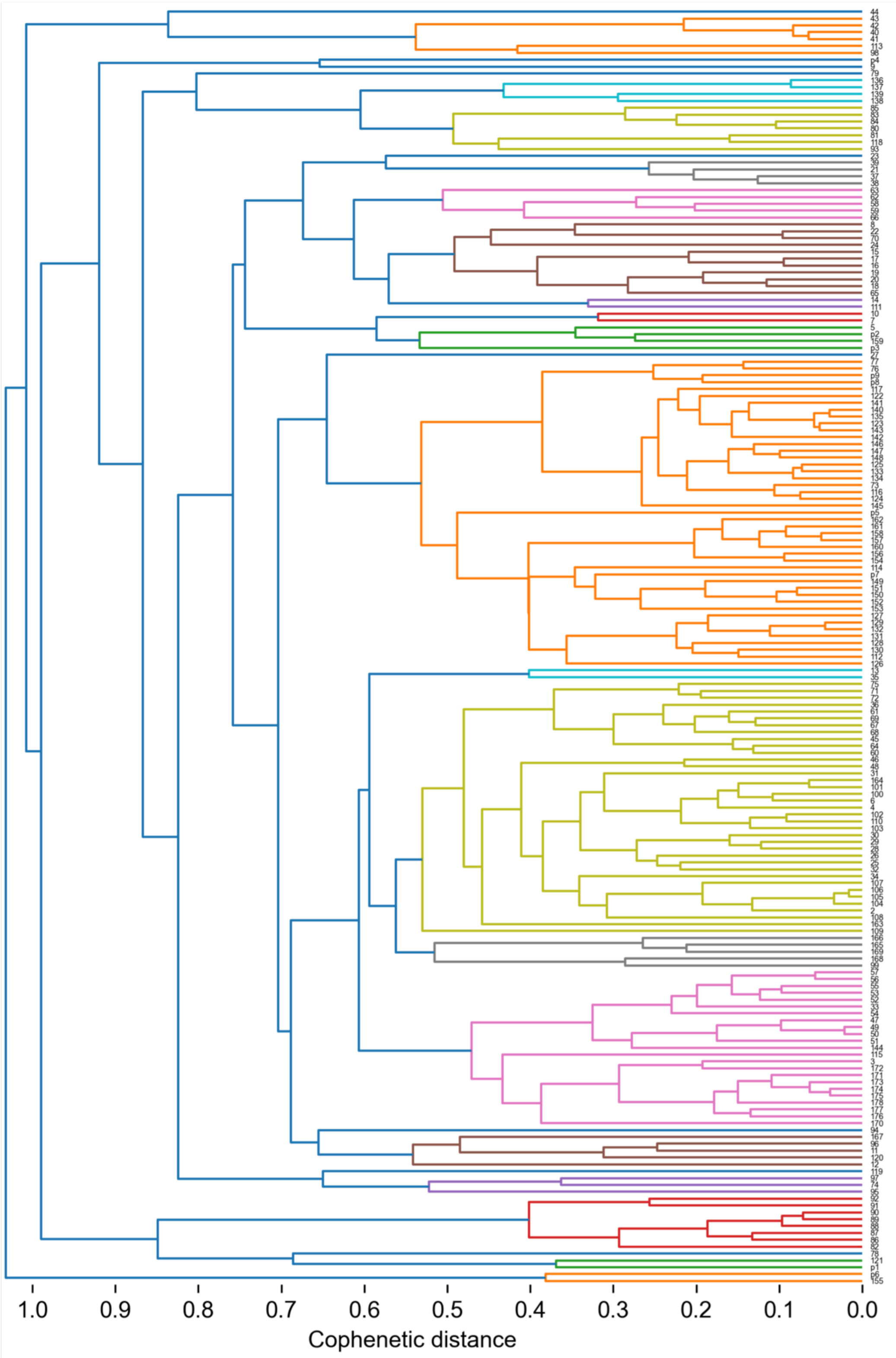


**Fig. S16. Agglomerative hierarchical clustering dendrogram derived from the raw Spearman rank correlations of per-residue RMSF values.** The dendrogram was constructed using the UPGMA algorithm, using pairwise correlation distance as the distance metric. Dendrogram leaves are clustered and colored using a cophenetic distance threshold of 0.55, a distance which allows p3V/P to be a member of one of the clusters.

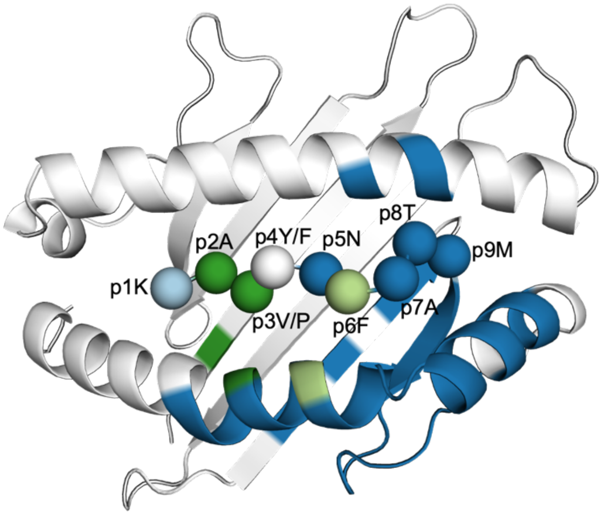


**Fig. S17. Clusters from agglomerative hierarchical clustering containing peptide residues mapped back onto the structure of H-2D$^b$/gp33.** In this mapping, p3V/P and p2A can be seen to cluster with Y159 on the α2 helix and M5 on the β1 strand. Both p5N and p7A form part of a larger cluster including the neighboring regions around Y159 on the α2 helix. H155 and p6F form their own cluster.

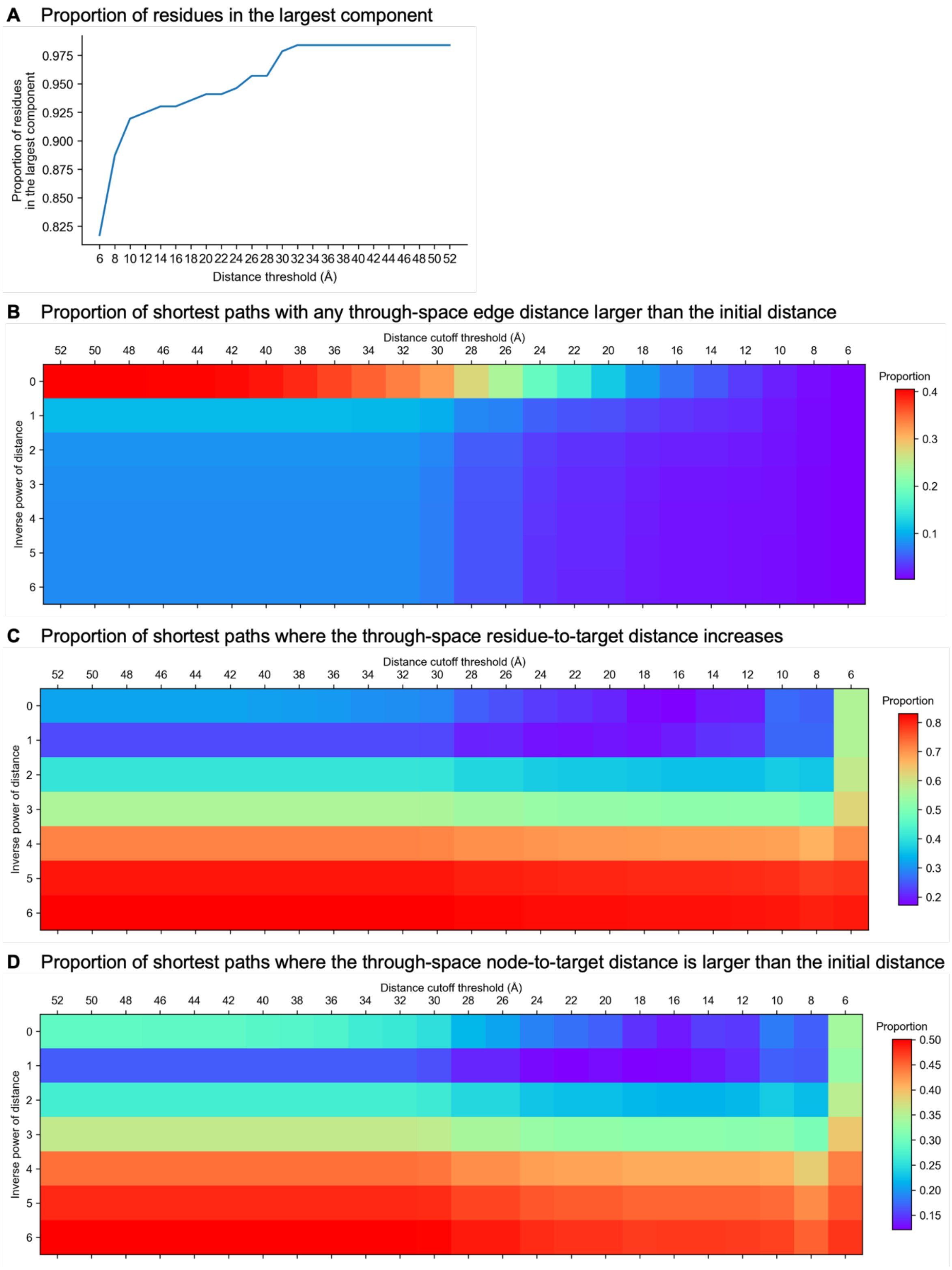


**Fig. S18. Overview of the network validation statistics used to determine which edge weight metric yielded the most biophysically relevant network.** A 2D scan of different distance cutoff thresholds and inverse powers of distance for network construction was performed. Note that a distance threshold of 52 Å is larger than any pairwise through-space $C_\alpha$ distance at the interface (51.1 Å). **A** Proportion of residues that remain in the largest connected component of the network using different distance thresholds. **B** Proportion of shortest paths where any through-space edge distance along the path is greater than the initial distance between the start and end nodes ($F_1$). The biophysically relevant network was chosen by minimizing edge weight complexity and **B** while maximizing **A** (>90%). **C** Proportion of shortest paths where the through-space node-to-target distance increases ($F_2$). **D** Proportion of shortest paths where any through-space node-to-target distance along the path is larger than the initial distance ($F_3$).

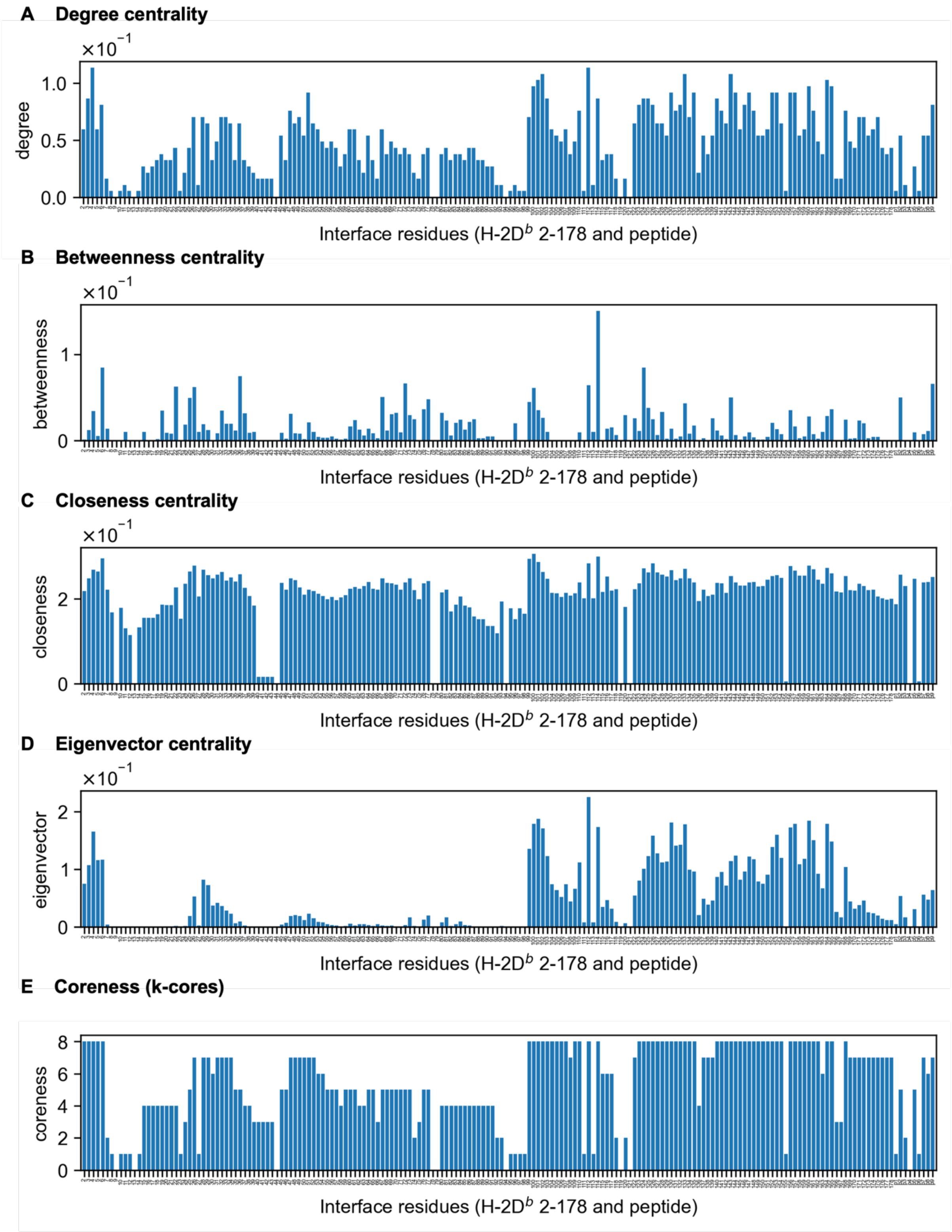


**Fig. S19. Centrality measures for each residue in the biophysically relevant network. A**, **B**, **C**, **D**, **E** The different centrality measures presented in **Fig. 4** are plotted accordingly for reference.

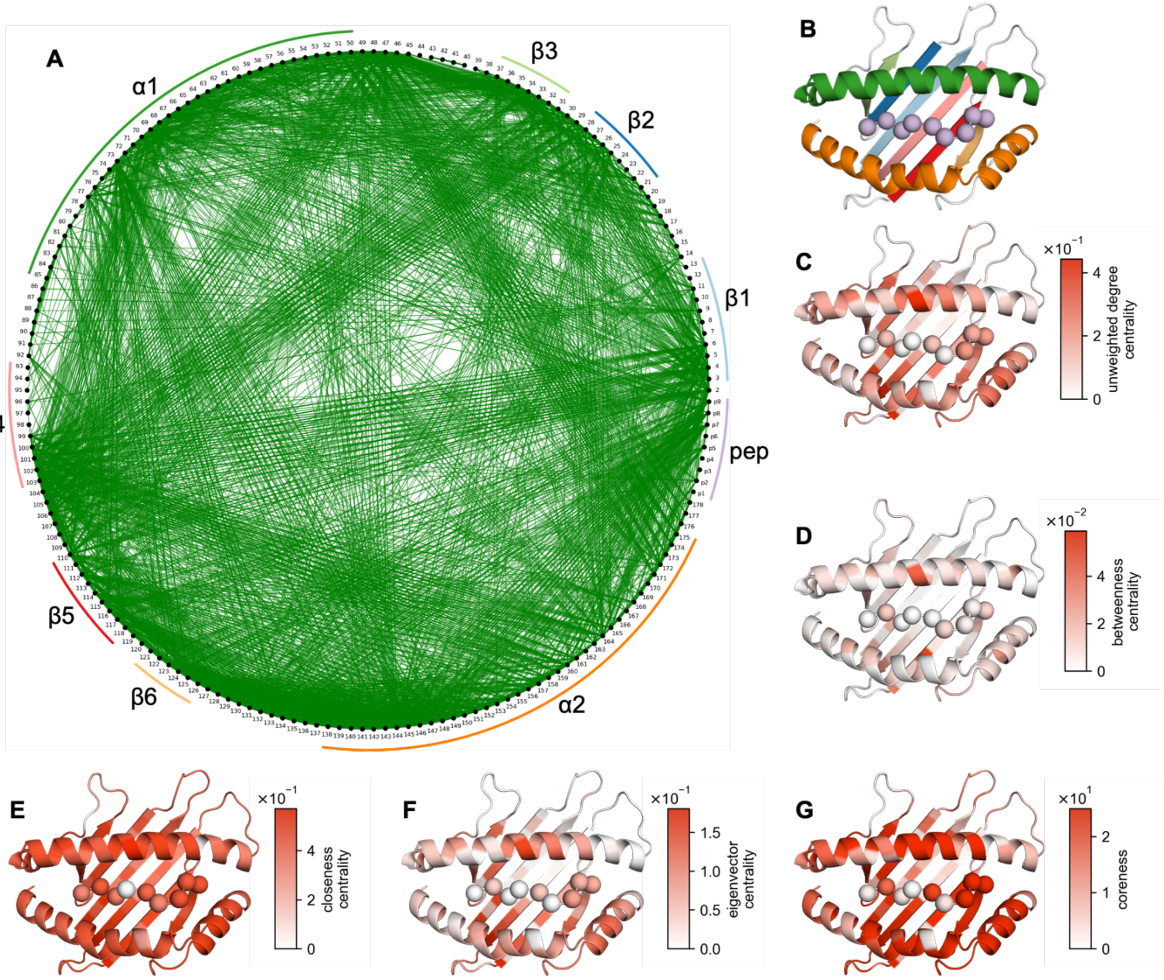


**Fig. S20. Network generated from the correlation matrix without implementing inter-residue through-space distance thresholds. A** The network encoding interface dynamics, where each node (black) represents an interface residue and edges (green) are significant correlations between residues whose alpha-carbon distance does not surpass 10 Å. Edge transparency and width are scaled according to $|\rho_s|$. Secondary structure features of the interface have been labeled. **B** Secondary structure features of the H-2D$^b$/peptide interface are colored according to the same label colors as in **A**. Various informative network centrality measures can be mapped to structure: **C** unweighted degree centrality, **D** betweenness centrality, **E** closeness centrality, **F** eigenvector centrality, and **G** coreness. Plots showing centrality measures per residue are provided (**Fig. S21**).

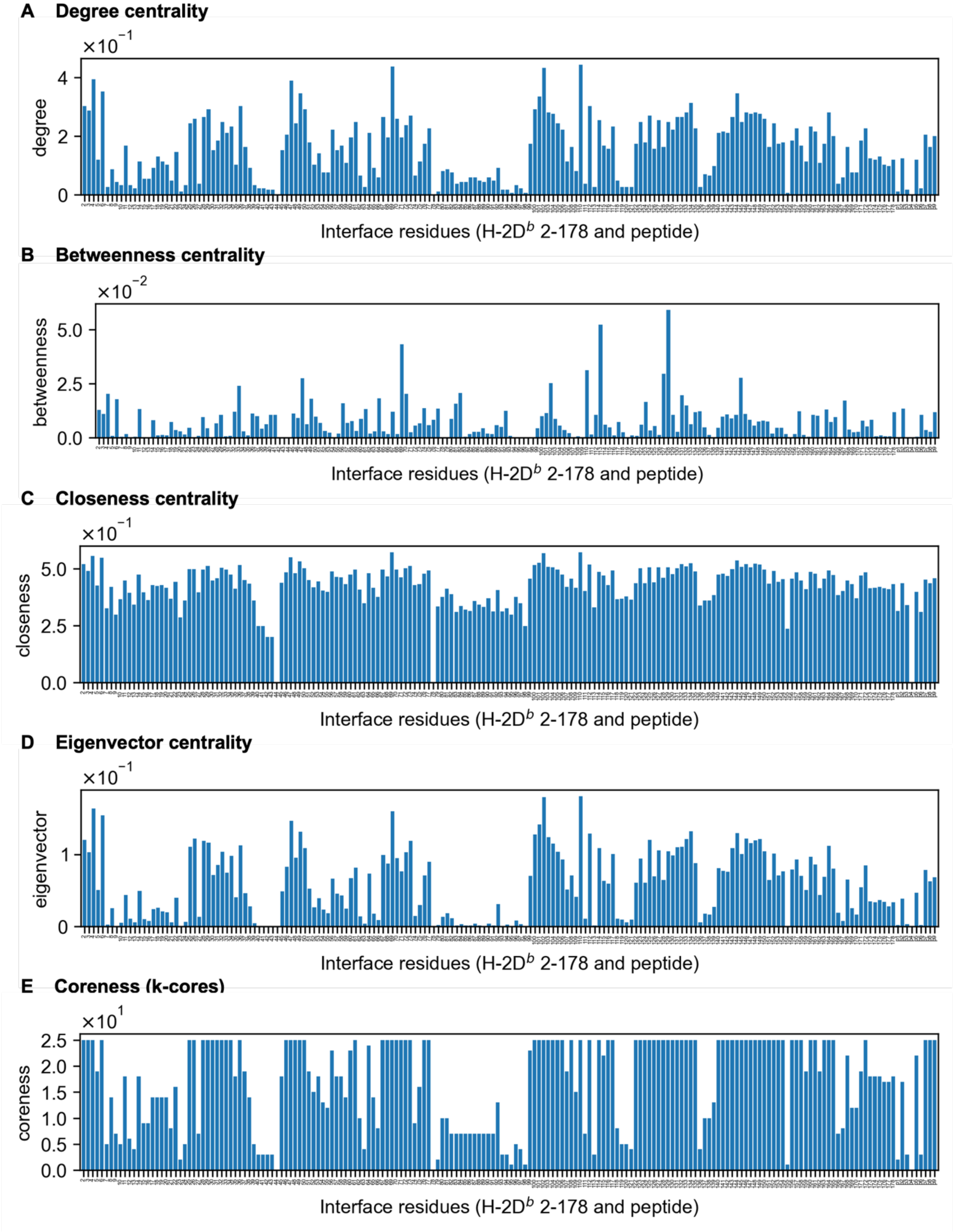


**Fig. S21. Centrality measures for each residue in the correlation network. A**, **B**, **C**, **D**, **E** The different centrality measures presented (**Fig. S20**) plotted accordingly for reference.

### References

1. A. Achour, J. Michaëlsson, R. A. Harris, J. Odeberg, P. Grufman, J. K. Sandberg, V. Levitsky, K. Kärre, T. Sandalova, G. Schneider, A structural basis for LCMV immune evasion: subversion of H-2D(b) and H-2K(b) presentation of gp33 revealed by comparative crystal structure analyses. *Immunity* **17**, 757–68 (2002).
2. L. M. Velloso, J. Michaëlsson, H.-G. Ljunggren, G. Schneider, A. Achour, Determination of structural principles underlying three different modes of lymphocytic choriomeningitis virus escape from CTL recognition. *J Immunol* **172**, 5504–11 (2004).
3. A. D. Duru, R. Sun, E. B. Allerbring, J. Chadderton, N. Kadri, X. Han, K. Peqini, H. Uchtenhagen, C. Madhurantakam, S. Pellegrino, T. Sandalova, P.-Å. Nygren, S. J. Turner, A. Achour, Tuning antiviral CD8 T-cell response via proline-altered peptide ligand vaccination. *PLoS Pathog* **16**, e1008244 (2020).
4. P. Emsley, B. Lohkamp, W. G. Scott, K. Cowtan, Features and development of Coot. *Acta Crystallogr D Biol Crystallogr* **66**, 486–501 (2010).
5. P. V Afonine, R. W. Grosse-Kunstleve, N. Echols, J. J. Headd, N. W. Moriarty, M. Mustyakimov, T. C. Terwilliger, A. Urzhumtsev, P. H. Zwart, P. D. Adams, Towards automated crystallographic structure refinement with phenix.refine. *Acta Crystallogr D Biol Crystallogr* **68**, 352–67 (2012).
6. D. Liebschner, P. V Afonine, M. L. Baker, G. Bunkóczi, V. B. Chen, T. I. Croll, B. Hintze, L. W. Hung, S. Jain, A. J. McCoy, N. W. Moriarty, R. D. Oeffner, B. K. Poon, M. G. Prisant, R. J. Read, J. S. Richardson, D. C. Richardson, M. D. Sammito, O. V Sobolev, D. H. Stockwell, T. C. Terwilliger, A. G. Urzhumtsev, L. L. Videau, C. J. Williams, P. D. Adams, Macromolecular structure determination using X-rays, neutrons and electrons: recent developments in Phenix. *Acta Crystallogr D Struct Biol* **75**, 861–877 (2019).
7. P. D. Adams, P. V Afonine, G. Bunkóczi, V. B. Chen, I. W. Davis, N. Echols, J. J. Headd, L.-W. Hung, G. J. Kapral, R. W. Grosse-Kunstleve, A. J. McCoy, N. W. Moriarty, R. Oeffner, R. J. Read, D. C. Richardson, J. S. Richardson, T. C. Terwilliger, P. H. Zwart, PHENIX: a comprehensive Python-based system for macromolecular structure solution. *Acta Crystallogr D Biol Crystallogr* **66**, 213–21 (2010).
8. B. T. Burnley, P. V Afonine, P. D. Adams, P. Gros, Modelling dynamics in protein crystal structures by ensemble refinement. *Elife* **1**, e00311 (2012).

9. B. T. Burnley, P. Gros, phenix.ensemble_refinement: a test study of apo and holo BACE1. *Computational Crystallography Newsletter* **4**, 51–58 (2013).
10. N. Ploscariu, T. Burnley, P. Gros, N. M. Pearce, Improving sampling of crystallographic disorder in ensemble refinement. *Acta Crystallogr D Struct Biol* **77**, 1357–1364 (2021).
11. J. Lemkul, From Proteins to Perturbed Hamiltonians: A Suite of Tutorials for the GROMACS-2018 Molecular Simulation Package [Article v1.0]. *Living J Comput Mol Sci* **1** (2019).
12. F. Ballabio, L. Broggini, C. Paissoni, X. Han, K. Peqini, B. M. Sala, R. Sun, T. Sandalova, A. Barbiroli, A. Achour, S. Pellegrino, S. Ricagno, C. Camilloni, l- to d-Amino Acid Substitution in the Immunodominant LCMV-Derived Epitope gp33 Highlights the Sensitivity of the TCR Recognition Mechanism for the MHC/Peptide Structure and Dynamics. *ACS Omega* **7**, 9622–9635 (2022).
13. M. J. Abraham, T. Murtola, R. Schulz, S. Páll, J. C. Smith, B. Hess, E. Lindahl, GROMACS: High performance molecular simulations through multi-level parallelism from laptops to supercomputers. *SoftwareX* **1–2**, 19–25 (2015).
14. K. Lindorff-Larsen, S. Piana, K. Palmo, P. Maragakis, J. L. Klepeis, R. O. Dror, D. E. Shaw, Improved side-chain torsion potentials for the Amber ff99SB protein force field. *Proteins* **78**, 1950–8 (2010).
15. H. J. C. Berendsen, J. R. Grigera, T. P. Straatsma, The missing term in effective pair potentials. *J Phys Chem* **91**, 6269–6271 (1987).
16. C. W. Hopkins, S. Le Grand, R. C. Walker, A. E. Roitberg, Long-Time-Step Molecular Dynamics through Hydrogen Mass Repartitioning. *J Chem Theory Comput* **11**, 1864–74 (2015).
17. B. Hess, H. Bekker, H. J. C. Berendsen, J. G. E. M. Fraaije, LINCS: A linear constraint solver for molecular simulations. *J Comput Chem* **18**, 1463–1472 (1997).
18. U. Essmann, L. Perera, M. L. Berkowitz, T. Darden, H. Lee, L. G. Pedersen, A smooth particle mesh Ewald method. *J Chem Phys* **103**, 8577–8593 (1995).
19. G. Bussi, D. Donadio, M. Parrinello, Canonical sampling through velocity rescaling. *J Chem Phys* **126**, 014101 (2007).
20. M. Parrinello, A. Rahman, Polymorphic transitions in single crystals: A new molecular dynamics method. *J Appl Phys* **52**, 7182–7190 (1981).

21. S. Nosé, M. L. Klein, Constant pressure molecular dynamics for molecular systems. *Mol Phys* **50**, 1055–1076 (1983).
22. R Core Team, "R: A Language and Environment for Statistical Computing" (Vienna, Austria, 2023); https://www.R-project.org/.
23. H. Wickham, J. Bryan, "readxl: Read Excel Files" (2023); https://CRAN.R-project.org/package=readxl.
24. Microsoft, S. Weston, "foreach: Provides Foreach Looping Construct" (2022); https://CRAN.R-project.org/package=foreach.
25. Microsoft, S. Weston, "doParallel: Foreach Parallel Adaptor for the 'parallel' Package" (2022); https://CRAN.R-project.org/package=doParallel.
26. H. Wickham, M. Averick, J. Bryan, W. Chang, L. D. McGowan, R. François, G. Grolemund, A. Hayes, L. Henry, J. Hester, M. Kuhn, T. L. Pedersen, E. Miller, S. M. Bache, K. Müller, J. Ooms, D. Robinson, D. P. Seidel, V. Spinu, K. Takahashi, D. Vaughan, C. Wilke, K. Woo, H. Yutani, Welcome to the tidyverse. *J Open Source Softw* **4**, 1686 (2019).
27. H. Wickham, *Ggplot2: Elegant Graphics for Data Analysis* (Springer-Verlag New York, 2016; https://ggplot2.tidyverse.org).
28. T. Kluyver, B. Ragan-Kelley, F. Pérez, B. E. Granger, M. Bussonnier, J. Frederic, K. Kelley, J. B. Hamrick, J. Grout, S. Corlay, P. Ivanov, D. Avila, S. Abdalla, C. Willing, Jupyter Development Team, "Jupyter Notebooks - a publishing format for reproducible computational workflows" in *Positioning and Power in Academic Publishing: Players, Agents and Agendas*, F. Loizides, B. Schmidt, Eds. (IOS Press, 2016; https://api.semanticscholar.org/CorpusID:36928206), pp. 87–90.
29. C. R. Harris, K. J. Millman, S. J. van der Walt, R. Gommers, P. Virtanen, D. Cournapeau, E. Wieser, J. Taylor, S. Berg, N. J. Smith, R. Kern, M. Picus, S. Hoyer, M. H. van Kerkwijk, M. Brett, A. Haldane, J. F. Del Río, M. Wiebe, P. Peterson, P. Gérard-Marchant, K. Sheppard, T. Reddy, W. Weckesser, H. Abbasi, C. Gohlke, T. E. Oliphant, Array programming with NumPy. *Nature* **585**, 357–362 (2020).
30. P. Virtanen, R. Gommers, T. E. Oliphant, M. Haberland, T. Reddy, D. Cournapeau, E. Burovski, P. Peterson, W. Weckesser, J. Bright, S. J. van der Walt, M. Brett, J. Wilson, K. J. Millman, N. Mayorov, A. R. J. Nelson, E. Jones, R. Kern, E. Larson, C. J. Carey, İ.

Polat, Y. Feng, E. W. Moore, J. VanderPlas, D. Laxalde, J. Perktold, R. Cimrman, I. Henriksen, E. A. Quintero, C. R. Harris, A. M. Archibald, A. H. Ribeiro, F. Pedregosa, P. van Mulbregt, SciPy 1.0 Contributors, SciPy 1.0: fundamental algorithms for scientific computing in Python. *Nat Methods* **17**, 261–272 (2020).

31. W. McKinney, "Data Structures for Statistical Computing in Python" in *Proceedings of the 9th Python in Science Conference* (2010; https://doi.curvenote.com/10.25080/Majora-92bf1922-00a), pp. 56–61.
32. J. D. Hunter, Matplotlib: A 2D Graphics Environment. *Comput Sci Eng* **9**, 90–95 (2007).
33. F. Pedregosa, G. Varoquaux, A. Gramfort, B. Michel V. and Thirion, O. Grisel, M. Blondel, R. Prettenhofer P. and Weiss, V. Dubourg, J. Vanderplas, A. Passos, D. Cournapeau, M. Brucher, M. Perrot, E. Duchesnay, Scikit-learn: Machine Learning in Python. *Journal of Machine Learning Research* **12**, 2825–2830 (2011).
34. A. A. Hagberg, D. A. Schult, P. J. Swart, "Exploring Network Structure, Dynamics, and Function using NetworkX" in *Proceedings of the 7th Python in Science Conference* (2008; https://doi.curvenote.com/10.25080/TCWV9851), pp. 11–15.
35. P. J. A. Cock, T. Antao, J. T. Chang, B. A. Chapman, C. J. Cox, A. Dalke, I. Friedberg, T. Hamelryck, F. Kauff, B. Wilczynski, M. J. L. de Hoon, Biopython: freely available Python tools for computational molecular biology and bioinformatics. *Bioinformatics* **25**, 1422–1423 (2009).
36. R. T. McGibbon, K. A. Beauchamp, M. P. Harrigan, C. Klein, J. M. Swails, C. X. Hernández, C. R. Schwantes, L.-P. Wang, T. J. Lane, V. S. Pande, MDTraj: A Modern Open Library for the Analysis of Molecular Dynamics Trajectories. *Biophys J* **109**, 1528–1532 (2015).
37. M. McKerns, L. Strand, T. Sullivan, A. Fang, M. Aivazis, "Building a Framework for Predictive Science" in *Proceedings of the 10th Python in Science Conference* (2011; https://doi.curvenote.com/10.25080/Majora-ebaa42b7-00d), pp. 76–86.
38. B. Phipson, G. K. Smyth, Permutation P-values Should Never Be Zero: Calculating Exact P-values When Permutations Are Randomly Drawn. *Stat Appl Genet Mol Biol* **9**, Article39 (2010).

39. Y. Benjamini, Y. Hochberg, Controlling the False Discovery Rate: A Practical and Powerful Approach to Multiple Testing. *J R Stat Soc Series B Stat Methodol* **57**, 289–300 (1995).
40. D. Müllner, Modern hierarchical, agglomerative clustering algorithms. (2011).
41. E. W. Dijkstra, A note on two problems in connexion with graphs. *Numer Math (Heidelb)* **1**, 269–271 (1959).
42. V. D. Blondel, J.-L. Guillaume, R. Lambiotte, E. Lefebvre, Fast unfolding of communities in large networks. *Journal of Statistical Mechanics: Theory and Experiment* **2008**, P10008 (2008).
